# Bifurcation in space: Functional modularity and distributed cognition in the neocortex

**DOI:** 10.1101/2023.06.04.543639

**Authors:** Xiao-Jing Wang, Junjie Jiang, Roxana Zeraati, Aldo Battista, Julien Vezoli, Henry Kennedy, Ulises Pereira-Obilinovic

**Author notes:** These authors contributed equally to this work (Xiao-Jing Wang, Junjie Jiang).

## Abstract

Recent reports of widespread neural representations challenge the notion that brain areas are specialized in distinct aspects of cognition. Here we tackle this challenge, using connectome-based neocortex models endowed with macroscopic gradients of neurobiological properties. We show that a bifurcation (an abrupt change of behavior) occurring locally in the cortical space gives rise to specialization of a subset of areas, that defines a multi-regional functional module, for subjective decision making or working memory coding. We found that mnemonic activity exhibits an inverted-V shaped pattern of timescales across the cortical hierarchy, which represents an experimentally testable prediction. A plethora of bifurcations in space coexist, corresponding to a variety of functional modules not well predicted by network analysis of structural modules; their associated timescale profiles could be observed in animals performing tasks that require different mental processes. These findings suggest bifurcation in space as a fundamental principle for understanding brain’s modular organization.

**Teaser:** The theory of bifurcation in space mechanistically accounts for the emergence of functional specialization of cortical areas dedicated to distinct cognitive functions that is compatible with distributed neural representations in a multiregional cortex.

## Introduction

Technical advances are enabling neuroscientists to measure calcium signals or record the spiking activity of large populations of single cells in behaving animals, opening a new era for the investigation of distributed neural computation across the multiregional brain (Wang 2022). Experimental reports of widespread neural signals of task-relevant information (Chen et al. 2024; Khilkevich et al. 2024; International Brain Laboratory et al. 2025; Findling et al. 2025). raised the question of how the notion of functional specialization/modularity (Fodor 1983; Kanwisher 2025) must be understood in terms of distributed processes in a multi-regional brain.

Consider, for instance, perceptual decision-making with two alternatives A and B, such as a classic random dots motion direction discrimination task (Newsome et al. 1989; Gold and Shadlen 2007). In an error trial, sensory evidence is in support of option A, but the brain’s judgment is option B. While early visual areas encode veridically visual inputs, neural correlates of subjective decision are observed in a number of brain regions that include posterior parietal and prefrontal cortices (Kim and Shadlen 1999; Roitman and Shadlen 2002; Bondy et al. 2025). How does a multi-regional functional module for subjective decisions emerge? This question is especially puzzling under the assumption that the cortex is composed of repeats of a canonical local circuit (Douglas and Martin 2004) and endowed with an abundance of long-range inter-areal connections (Markov et al. 2014; Harris et al. 2019; Gămănuţ et al. 2018).

We have previously developed large-scale cortical models of distributed working memory, a basic cognitive function, in both macaque monkey (Mejias and Wang 2022; Froudist-Walsh et al. 2021) and mouse (Ding et al. 2024) on the basis of recently available area-to-area connectivity data. Critically, unlike the conventional view of a cortical network as a graph where parcellated areas (nodes) are identical and differ from each other only by their connections, our model incorporated area-to-area heterogeneities, which represents a reappraisal of the canonical local circuit principle (Douglas and Martin 2007). These variations display macroscopic gradients along an axis such as the anatomically defined cortical hierarchy (Wang 2020). Model simulations revealed an abrupt transition that separates cortical areas exhibiting information-coding self-sustained persistent activity from those that do not. In a single instance we mentioned that this observed phenomenon in model simulations was “akin to a bifurcation in cortical space” (Mejias and Wang 2022), which was repeated in a review (Wang 2022). However, this idea has remained purely speculative up to now.

The mathematical term “bifurcation” denotes the sudden onset of a qualitatively novel behavior under a graded change of the properties of a dynamical system (Strogatz 2016). For instance, a single neuron displays an abrupt transition from a steady-state membrane potential to repetitive firing of spikes as the intensity of an input current is gradually increased above a threshold level. The idea of bifurcation in space implies a bifurcation localized at a position in the cortical mantle rather than for the entire dynamical system, robustly without the need of parameter tuning. In this work, we mathematically prove the concept of bifurcation in space, and we show that it may be realized even under the condition that no isolated area can generate persistent activity so that the observed working memory representation must be a collective phenomenon emerging from long-distance inter-areal connection loops, thereby extending the concept of synaptic reverberation (Goldman-Rakic 1995; Amit 1995; Wang 2001) to the large-scale multi-regional brain.

The model reproduces the observation that during working memory of visual motion information, the middle temporal (MT) area does not, while its monosynaptic projection target, the medial superior temporal (MST) area does, show persistent activity during a mnemonic delay in macaque monkeys (Mendoza-Halliday et al. 2014). Furthermore, we show functional modularity for subjective decision-making in the macaque cortex, thereby demonstrating bifurcation in space beyond working memory.

Notably, we found that the timescales of neural firing fluctuations are maximal at the spatial location of a sharp transition separating areas devoid of persistent activity and those that exhibit robust persistent activity. Consequently, cortex displays an inverted-V shaped pattern of time constants of neural fluctuations during a mnemonic state, in striking contrast to the previous finding of a roughly monotonic increase of time constants along the cortical hierarchy during a resting state (Chaudhuri et al. 2015; Murray et al. 2014; Song et al. 2024). The phenomenon disappears when a connectome-based model of macaque or mouse operates in an “everything, everywhere” regime, demonstrating the inverted V-shaped pattern of timescales as a signature of functional modularity. This model prediction can be tested experimentally using a working memory task while recording from many cortical areas.

Furthermore, a single large-scale neocortex model (with fixed parameter values) displays simultaneously many bifurcations in space, each associated with a multiregional functional module mathematically identified as a spatial attractor. We show that these functional modules defined in terms of spatial attractors generally cannot be predicted by structural modules defined by graph theory, namely using a community detection algorithm applied to inter-areal connectivity matrix. These multi-regional modules can potentially subserve distinct internally driven brain functions.

On the basis of these findings, we propose that functional modularity can be defined as a selective subset of cortical areas, which are not necessarily spatially congruent, engaged in a distinct brain function. This theory reconciles functional specialization with distributed neural representation and processing across multiple brain regions.

## Results

### A connectome-based monkey cortex model displays bifurcation in space during decision-making and working memory

We start with an extension of a connectome-based macaque cortex model (Mejias and Wang 2022; Froudist-Walsh et al. 2021) to 41 parcellated areas (Fig. 1a, Fig. S1). The model incorporates hitherto unpublished connectivity data on area MST, specifically motivated by physiological observations that neurons in MST, but not their main monosynaptic afferents from the adjacent MT area, display delay period activity in a working memory task (Mendoza-Halliday et al. 2014). In a simple version of our model, suitable for stimulus-selective decision-making and working memory, each area has two selective excitatory populations A and B, and one inhibitory population (Mejias and Wang 2022). Both long-range and local recurrent excitation strength follow a macroscopic gradient proportional to the hierarchical position of each area (Chaudhuri et al. 2015; Wang 2020).

**Figure 1.**
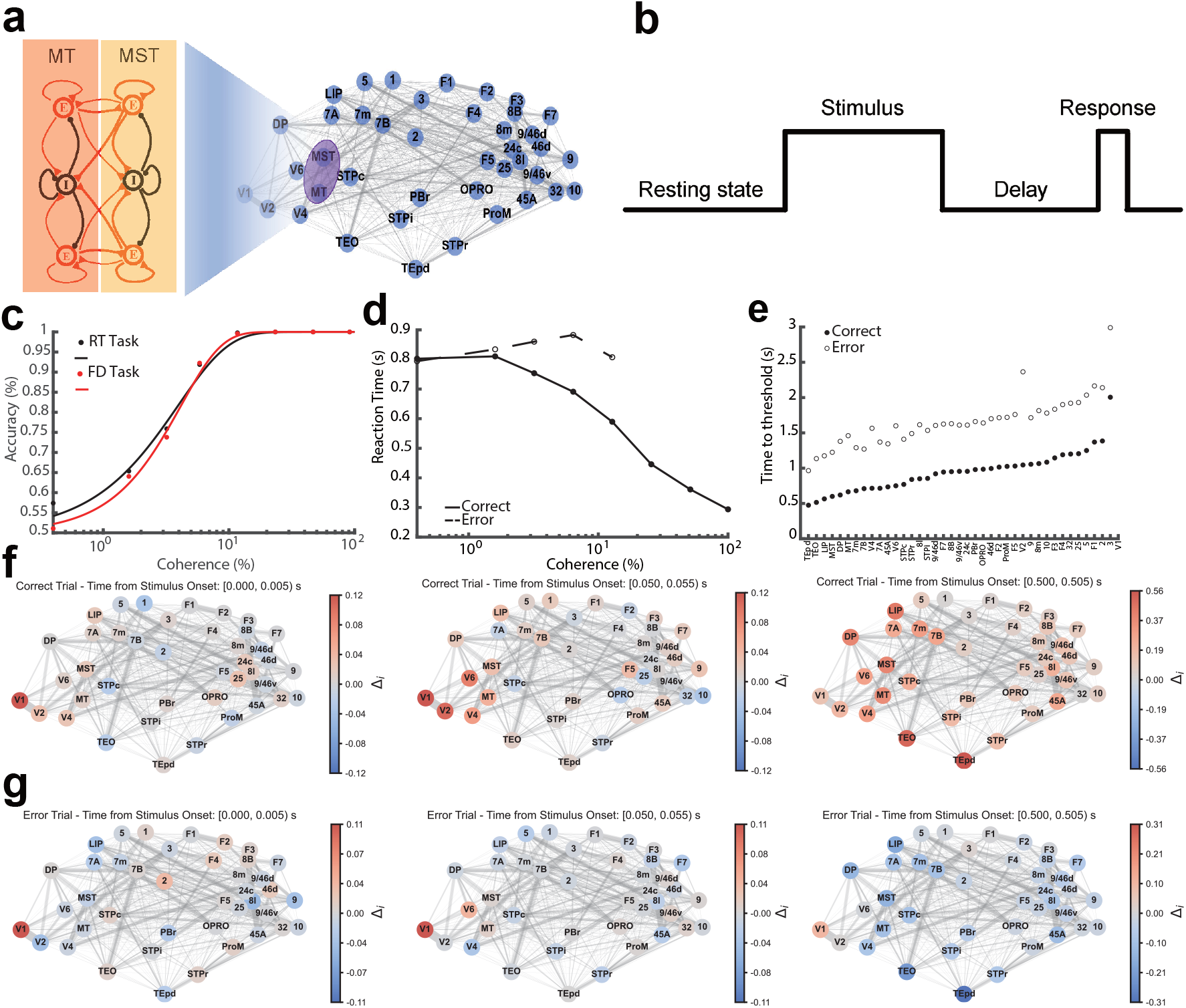
Connectome-based model of macaque monkey cortex for distributed perceptual decision-making and working memory. (a) Model scheme. Right: experimentally measured connections between areas as a graph (weight is indicated by line thickness); Left: local and long-range connections between two areas (illustrated by MT and MST) via directed projections to excitatory and inhibitory neural pools. (b) Timeline of a model simulated task that requires both perceptual decision-making and working memory. The model is initially in a resting state. A stimulus is presented for a fixed duration, with the task difficulty parametrically varied from trial to trial by the relative difference in the inputs onto the two stimulus-selective neural pools in V1, and a judgment is produced by winner-take-all attractor dynamics in a subset of cortical areas (functional module for subjective choice). The choice must be internally stored in working memory over a delay period. The memory-guided response could be an eye saccade or moving a hand towards one of two choice targets, not explicitly modeled here. In reaction time (RT) task simulations, the response is generated when Δ_*LIP*_ (*t*) exceeds a preset positive or negative threshold for choice A or B, respectively. (c) Psychometric function (accuracy) for the RT and FD tasks and (d) chronometric function (average reaction times in correct and error trials) of the model, compatible with monkey behavior in random dots motion direction discrimination experiments (Mazurek et al. 2003). (e) Rank-ordered time-to-threshold for all brain areas. (f, g) Activity maps across different brain areas exhibit distinct structures in correct (f) and error (g) trials depending on the time from the stimulus onset (indicated in panel titles). Red indicates areas (indexed by *i*) signaling 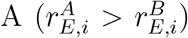 when input to excitatory population A is larger, and blue indicates areas signaling 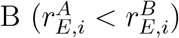. The location of areas on graph maps is computed by projecting 3D location of each area in the inflated surface map to 2D, based on Markov et al. (2014). For the temporal dynamics of the activity maps, see Movie 1 (correct trials) and Movie 2 (error trials).

We simulated the model for a task that depends on perceptual decision-making and working memory. In a random-dots motion direction discrimination experiment (Roitman and Shadlen 2002; Gold and Shadlen 2007), task difficulty varies from trial to trial by changing the motion coherence, the fraction of dots moving coherently from 100% to 0%. The subject has to decide whether the net motion direction is A (e.g., left) or B (right) (two-alternative forced choice task). In the model, motion coherence is implemented by the relative strength of external inputs into the two (A and B) selective excitatory neural populations in V1 (Wang 2002; Wong and Wang 2006), hence neural signals in V1 encode the physical stimulation, whereas a subset of other areas in the multi-regional model are expected to determine a categorical choice A or B. The latter is dissociable from sensory evidence when the motion coherence is zero and in error trials.

As in the monkey experiments (Roitman and Shadlen 2002), we simulated two versions of the direction discrimination task. In the reaction time (RT) version, a decision is read out when the accumulated information reaches a threshold (for instance, when the neural activity in the lateral intraparietal area (LIP) ramps up to a preset level (Roitman and Shadlen 2002).A different decision readout from a multi-regional system is conceivable). In the fixed duration (FD) version, a stimulus is presented for two seconds, followed by a delay period when the choice must be held in working memory to guide a later behavioral response (Fig. 1b). This task thus engages both decision-making and working memory. The psychometric functions of the model for the RT and FD tasks (Fig. 1c) are comparable to the observed monkey performance.

To quantify the complex neural dynamics during decision-making in space and time, we computed the normalized difference between the firing rates 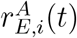 and 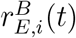 of two excitatory populations, as 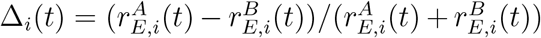 in each parcellated area *i* of our model. In model simulations of the RT task, the RT is read out when Δ_*i*_(*t*) in LIP reaches a (positive or negative) threshold (see Methods). In accordance with experimentally observed behavior the average RT as a function of coherence is shown in Fig. 1d, with longer RTs in error (open circle) than in correct (filled circle) trials.

Fig. 1f and g (see also Movies 1, 2) show the spatiotemporal dynamics of Δ_*i*_(*t*) in a correct trial and an error trial, respectively, in model simulations with evidence in favor of A. At the stimulus onset (left panel), the input to neural population A is larger than to B, and Δ(*t*) is positive in V1 (red); the signal propagates to higher areas where strong recurrent dynamics lead to a categorical winner (A or B) (middle and right panels). This was quantified by the time-to-threshold that varies from area to area (Fig. 1e); areas TEpd, TEO, and LIP are the first to reach the threshold in the correct (Fig. 1f, Movie 1) and error (Fig. 1g, Movie 2) trials. Compared to the correct trial, in the error trial, there is a period (Fig. 1g, middle panel) when some areas, such as V1 and MT, represent the veridical evidence A (with positive Δ_*i*_(*t*), red). In contrast, others signal the subjective choice B (with negative Δ_*i*_(*t*), blue). This suggests a bifurcation in space that unfolds in time. Late in time, MT also reflects the subjective choice (Fig. 1g, right panel), reproducing the observed choice correlate in MT neurons of behaving monkeys (Britten et al. 1996). Delayed choice signals in MT have been suggested to result from top-down inputs (Wimmer et al. 2015) that naturally occur in our connectome-based multiregional cortex model.

#### The inverted-V shaped profile of time constants in macaque monkey and mouse

The FD task allowed us to examine working memory of a choice across a delay period in the absence of external stimulation. The model’s performance in the FD task is similar to that of the RT task (Fig. 1c). The connectome-based model of macaque cortex (Fig. 2a) exhibits the coexistence of a resting state (a firing rate around 1.5 *Hz* for all areas) and stimulus-selective persistent activity states encoding working memory of A or B (Fig. 2b, Fig. 8a, Fig. S2a-e). The persistent activity state involves multiple areas and is dynamically stable against brief, small perturbations; we call such an internal state a *spatial attractor* that defines a multi-regional working memory module. Capturing macaque monkey physiological experimental observations (Leavitt et al. 2017), association cortical areas but not early sensory areas are engaged in stimulus-selective persistent firing during working memory (Fig. 2b). The area MST exhibits sustained activity throughout the delay period, whereas MT does not manifest such persistent activity, as evidenced in Fig. 2b. Thus, our model successfully reproduces the sharp transition in the cortical hierarchy observed by Mendoza-Halliday et al. (2014) (Mendoza-Halliday et al. 2014), and offers a general account of the emergence of functional specialization for working memory (Leavitt et al. 2017).

**Figure 2.**
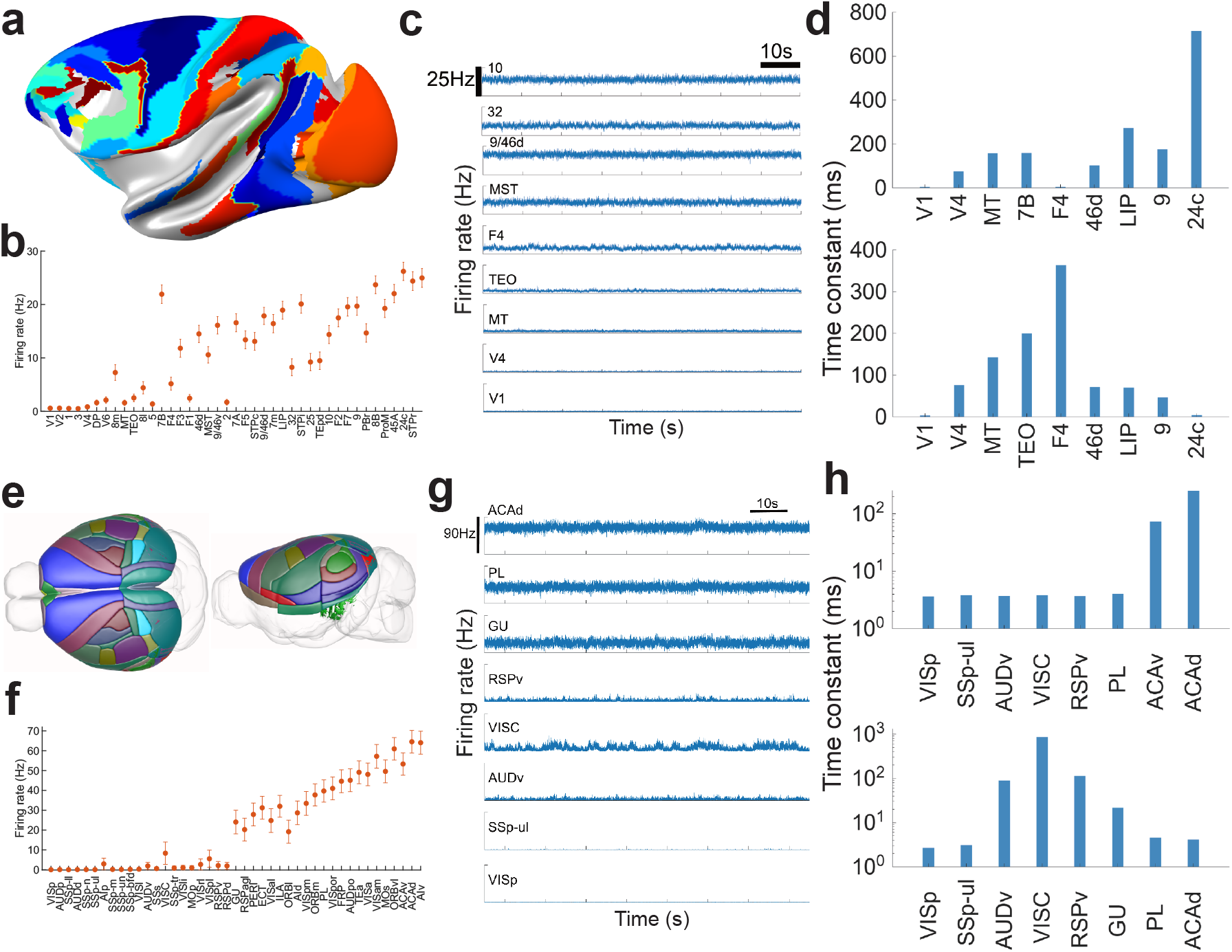
Bifurcation in space of connectome-based cortical models of the macaque monkey (a-d) and mouse (e-i). (a) Lateral view of macaque neocortex surface with areas in color included in the model. (b) Firing rate of 41 brain areas, ranked by the hierarchical position in a working memory delay period. (c) Firing rate time series of 9 sample brain areas. (d) Time constants of neural fluctuations in 9 selected brain areas for resting (upper panel) and delay period working memory (lower panel) states. (e) Superior and lateral view of mouse cortex surface with model areas in color. (f) Working memory presentation is distributed in a subset of areas (functional module), shown with the firing rate of persistent activity plotted against the hierarchical position of 43 brain areas during the delay period. (g) Firing rate time series of 8 sample areas. (h) Time constants of neural fluctuations in 8 selected areas in resting state (upper panel) and during mnemonic delay period (bottom panel). During working memory, the time scales of neural activity fluctuations exhibit an inverted-V shaped pattern along the hierarchy, with a peak indicating critical slowing down near the sharp transition locally in space.

We examined the timescale pattern along the cortical hierarchy in both the resting state and the working memory state of our model, by calculating the autocorrelation function of stochastic neural activity from which a timescale is extracted (see Methods). In the resting state prior to stimulus onset, the time constants dominating neural fluctuations were roughly a monotonic function of the hierarchy (Fig. 2d, upper panel, see also Fig. S2e, left), in agreement with previous modeling (Chaudhuri et al. 2015) and empirical (Murray et al. 2014; Song et al. 2024; Shi et al. 2025) findings. When the analysis was repeated for the mnemonic delay period, we found that the timescale of stochastic neuronal activity showed a peak, with short time constants for areas both at high and low hierarchical levels. The peak is located around the sharp transition in hierarchical space where the working memory module emerges (Fig. 2d, lower panel; see also Fig. S2e, right). The contrast is striking for area F4 (a premotor area) which shows a short timescale at rest and a long timescale during working memory, and area 24c (part of the anterior cingulate cortex) for which the the opposite holds.

Recall that the same model system, in the absence of external input, exhibits spontaneous activity prior to stimulus presentation and persistent activity during the delay period. The above results show that the timescale pattern across a large-scale cortex depends on the internal state of a strongly recurrent nonlinear neural circuit. For a working memory state, the time constant of neural fluctuations can become very long near the transition (see also the next Section), a phenomenon akin to “critical slowing down” in statistical physics, namely fluctuations occur over unusually long timescales near a phase transition such as melting of ice (Scheffer 2009; Tredicce et al. 2004). However, the analogy is inadequate in three substantial ways. First, the transition here is not for the entire system and does not require varying a control parameter (such as tuning the temperature to 0 degree Celsius for the solid-to-liquid transition of H_2_O), although what particular area displays the maximal time constant of mnemonic firing fluctuations depends on model details. Second, the transition occurs locally, in spite of the abundance of long-range interareal interactions (in contrast to local interactions between molecules in water). Third, a transition is defined for a specific internal state corresponding to a spatial attractor. As we will see later, there are many distinct spatial attractors coexisting in the same model system.

We then considered if critical slowing down also occurs during a mnemonic delay period in a connectome-based large-scale model of the mouse cortex (Ding et al. 2024). The model contains 43 cortical areas in the common coordinate framework v3 atlas (Oh et al. 2014) (Fig. 2e) with a quantified hierarchy. It incorporated experimentally measured heterogeneity of the density of parvalbumin (PV)-expressing interneurons (Kim et al. 2017); the model showed that an area with a lower PV density is more excitable and prone to display elevated persistent activity. The mouse model also exhibits the coexistence of a resting state and persistent activity states suitable to subserve working memory function (Fig. 2f, Fig. S2f). An autocorrelation analysis of the neural firing fluctuations showed a rougly increasing trend along the cortical hierarchy of time constants in the resting state (Fig. 2h, upper panel), and an inverted-V shaped profile of time constants during working memory (Fig. 2h, lower panel, see also Fig. S2g, right).

Functional modularity is not always guaranteed. It is robust against parameter changes, but can disappear under certain conditions. One key parameter controls the relative strengths of a top-down projection onto excitatory versus inhbitory neurons in a target area. When that ratio is increased, inter-areal excitatory-to-excitatory reverberation becomes more powerful, and all areas including primary sensory areas like V1 now display elevated persistent activity compared to the resting state in our models for mouse (Fig. 3a, left panel) and monkey (Fig. 3b, left panel). In this “everything everywhere” scenario of distributed working memory coding, delay period activity is no longer characterized by the inverted-V shaped pattern of timescales of neural fluctuations (Fig. 3a, right panel for the mouse model; Fig. 3b, right panel for the macaque model). This result suggests that the inverted-V pattern of timescales represents a manifestation of the presence of functional modularity for working memory. For instance, it is possible that under certain conditions the mouse cortex shows no functional modularity, and the inverted-V shaped pattern of timescales would be absent. This strong model prediction is testable experimentally.

**Figure 3.**
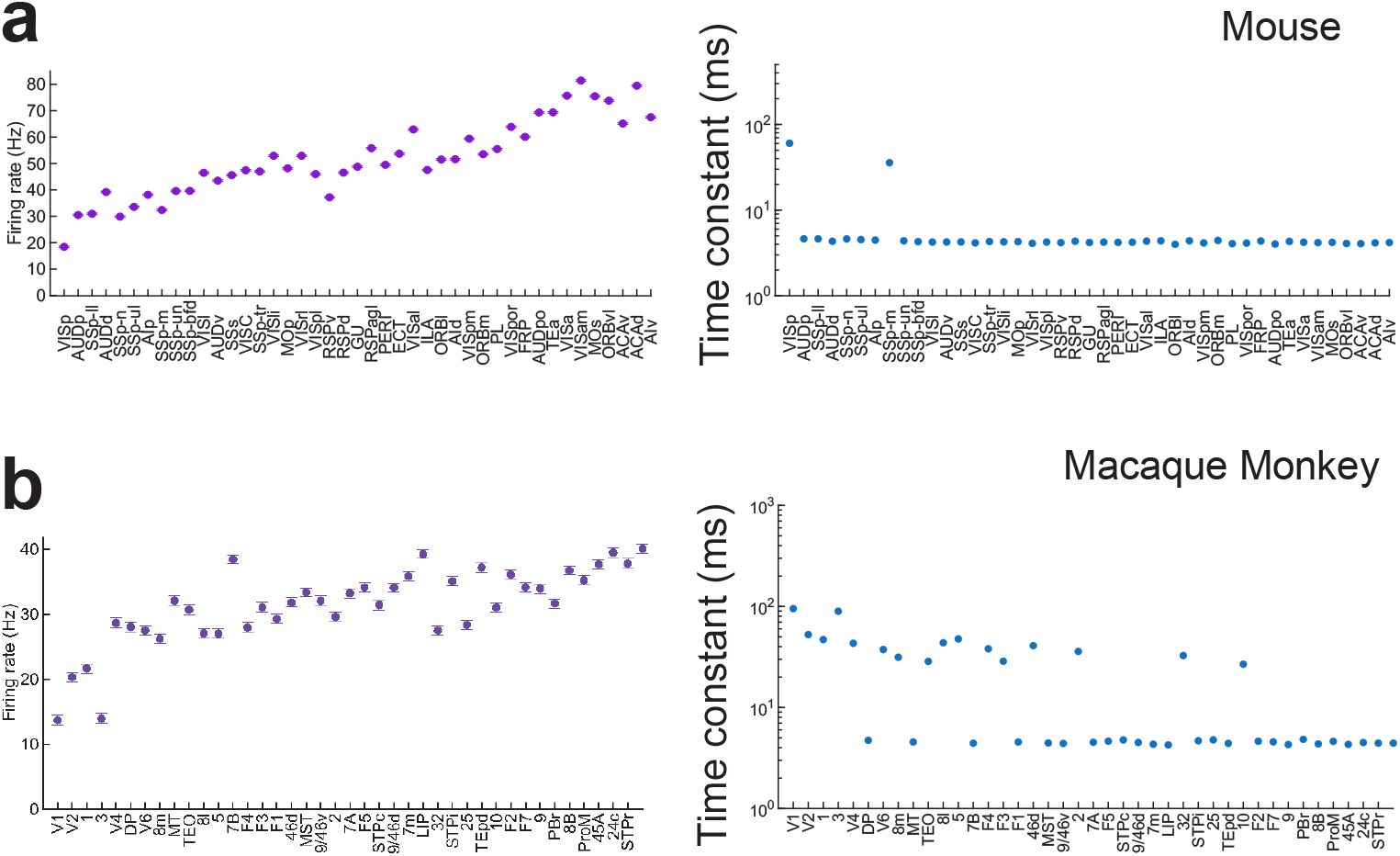
With a few parameter changes, functional modularity is abolished in the mouse model (a) and macaque monkey model (b). Working memory representation engages all areas during the delay period in an “everthing, everywhere” regime (left panels). The time scales of mnemonic persistent activity fluctuations no longer show an inverted-V shapped pattern along the cortical hierarchy (right panels). The parameter *Z* controls the relative strengths of a top-down projection to recipient excitatory and inhibitory neurons, respectively, in areas at lower hierarchical positions; a large *Z* increases exciatory-to-excitatory interactions that involve early sensory areas and could abolish functional modularity. *Z* = 2 here compared to 0.6 for the mouse model and 0.8 for the macaque model, respectively in Figure 2. Other parameter changes are found in the Method.

#### A generative model for the mammalian neocortex

To investigate the mathematics of bifurcation in space, we need to “zoom in” close to an abrupt transition point, similar to a phase transition such as the melting of ice when the temperature is adjusted to precisely zero degree Celsius, but here at a particular spatial location across the cortical hierarchy separating areas engaged in a working memory representation from those that are not. However, a connectome-based model of the mouse or macaque monkey cortex has a relatively small number of areas; thus, the minimal distance between any pair of areas along the hierarchy cannot be reduced as much as desired. To overcome this limitation, we used a generative model of a spatially embedded mammalian cortex (Fig. 4a) (Song et al. 2014) that shares the same statistical distributions empirically observed in the interareal connectivity (Ercsey-Ravasz et al. 2013; Markov et al. 2014; Song et al. 2014). In the model, the number of brain areas can be arbitrarily large, enabling examination of the bifurcation phenomenon close to a transition point. Moreover, the model can be used to produce multiple *realizations* of mammalian neocortical networks by random sampling of connectivity matrices (see Fig. 4b for one network realization), which enabled us to assess the robustness of our results over multiple network realizations.

**Figure 4.**
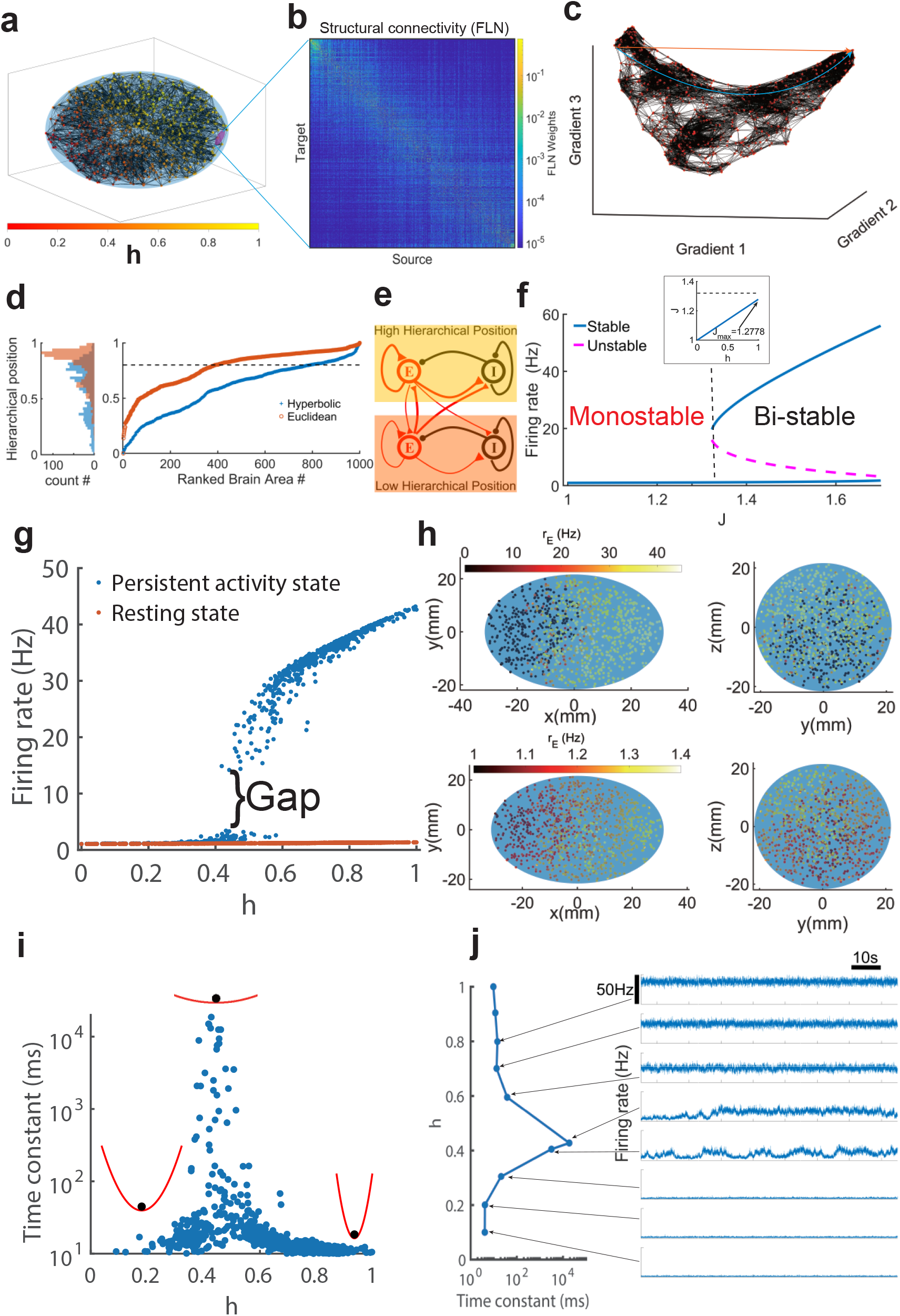
Spatially embedded generative model of a cortex. (a) Network connectivity of 1000 parcellated cortical areas produced by a generative model of the mammalian cortical connectivity (Song et al. 2014), which is spatially dependent, directed, and weighted. (b) Connectivity matrix of the model. (c) Three-dimensional diffusion map embedding of the generative cortical network. Red dots: 1000 cortical areas, black lines: the nearest neighbor links in the embedding space. The axes correspond to the first three principal gradients from the diffusion map embedding (see Methods). Using the diffusion map, the hierarchical position of any area is defined based on Euclidean (brown) or hyperbolic (blue) distance to a starting area at the bottom of the hierarchy (see Methods). (d) Hyperbolic metric yields a more linear increase than Euclidean metric along the hierarchy. (e) Local circuit model scheme for each cortical region, with recurrently connected excitatory (E) and inhibitory (I) populations. The strength of local and long-range connections is indicated by line thickness. (f) Bifurcation diagram of an isolated cortical circuit (see Eq. (7) in SI). In the large-scale system, the local E-to-E and E-to-I weights are scaled by a factor *J*, which displays a macroscopic gradient as a function of the hierarchical position *h* (Inset), with the maximal value *J* below the threshold level to induce bistability. (g) Resting (brown) and persistent activity states (blue) are shown with firing rates plotted against areas ranked by hierarchical position. In the persistent activity state, a subset of areas represents working memory and are separated from the rest of the system by a firing rate gap. (h) Front view (left) and side view (right) of the spatial distribution of the persistent activity state (top) and resting state (bottom) in the generative model’s ellipsoid. (i) The time constant of all brain areas for the persistent activity state of panel (b). Potential plots illustrate schematically the idea that neural activity state is more stable for areas at both low and high hierarchical positions (left and right) and marginally stable near the bifurcation (middle), giving rise to an inverted-V-shaped pattern of time scales. (j) Left, time constant of 10 selected brain areas; right, firing rate time series of 8 selected cortical areas for the cortical network in the persistent activity state.

We applied the diffusion map embedding (Coifman and Lafon 2006; Margulies et al. 2016) to our model. In the embedding space obtained through this nonlinear dimensionality reduction method, closer areas share more paths connecting them with stronger connections, while areas further apart share fewer paths and weaker connections (see Methods). Interestingly, the embedding of the generated connectivity conforms to a low-dimensional hyperbolic shape (see Fig. 4c). The cortical area at one of the tips of the hyperbolic shape was chosen as the start of a hierarchy. We found that for the hyperbolic distance (the distance defined along the hyperbolic shape), cortical areas display a smooth progression evenly distributed across hierarchical positions (see Fig. 4d blue trace), in contrast to the Euclidean distance where a significant fraction of cortical areas are concentrated around the hierarchy value 0.8 (see Fig. 3d brown trace). Therefore, we used the hyperbolic distance to define the hierarchical position. Strikingly, after remapping each brain area’s hyperbolic hierarchical position into the ellipsoid’s position (see Fig. 4a), we found that the hierarchical position increases along the major axis of the ellipsoid, similar to the hierarchy of the mammalian cortex along the caudo-rostral axis (Fig. S5a). The alignment of hierarchical positions with the ellipsoid’s major axis likely reflects the underlying network connectivity and embedding geometry. This network structure drives the primary gradient of the diffusion map to align with the longest axis, corresponding to the caudo-rostral axis of the mammalian cortex.

Using the above defined hierarchy, we built a simplified yet biologically realistic model of the neocortex. Each brain area is modeled as a local canonical circuit of recurrently connected excitatory and inhibitory populations (Fig. 4e and Methods). Consistent with the macroscopic gradient of excitation observed in the cortex (Wang 2020), the excitatory connection strength *J* increases with the hierarchical location *h* of areas (inset in Fig. 4f). When decoupled, each cortical area exhibits a low firing rate resting state when the synaptic excitation level is below a threshold corresponding to *J*_threshold_ ≈ 1.32 (*J < J*_threshold_) and exhibits bistability between a resting state and an elevated persistent activity state when *J > J*_threshold_. In the scenario where the maximal value of *J* at the top of the hierarchy is smaller than *J*_threshold_ (Fig. 4f), isolated areas are unable to generate elevated persistent activity, but working memory representation may emerge from long-distance area-to-area connection loops.

The connected network of interacting areas exhibits active states (where some cortical areas display persistent high firing rates while others display low firing rates) suitable for working memory that coexist with a resting state (where all areas of the network exhibit a low firing rate state ~ 1 *Hz*). An example is shown in Fig. 4g. For this active state, those areas engaged in persistent activity are located higher in the hierarchy and are separated from those with low activity by a firing rate gap, indicating a transition zone. The size of this transition zone systematically shrinks with the increasing size of the network (Fig. S6b). We expect that in the limit of infinitely large networks the transition zone shrinks to a point that we denote as the bifurcation location in hierarchical space (Fig. 4g, Fig. S6b). Interestingly, at the bifurcation point, there is a firing rate gap reminiscent of a first-order phase transition in statistical physics (Landau and Lifshitz 2013).

Mapped into our generative model’s ellipsoid where connectivity was originally embedded, the activity of an area increases along its major axis in both active and resting states (Fig. 4h). In contrast to the resting state, where the firing rate increases smoothly along the hierarchy (Fig. 4h, lower panel), in the persistent activity state, there is a firing rate gap between the caudal and rostral areas (Fig. 4h, upper panel). Furthermore, the persistent firing rate increases along the minor axis *z* (Fig. 4h).

We quantified the time scale of persistent activity fluctuations in the generative network model, using the same autocorrelation analysis as for the connectome-based cortex models (Fig. 2). We found that the time scale of neural fluctuations increases from milliseconds to tens of seconds for areas near the transition point (Fig. 4i), displaying the critical slowing down phenomenon. The time scale profile along the hierarchy is of an inverted V-shape with fast fluctuations for areas low and high in the hierarchy and very slow fluctuations for areas in the bifurcation region. This inverted V-shape divergence in time scale is characteristic of critical slowing down and is only observed for the mnemonic active state. In contrast, in the resting state, the time scale increases monotonically with the hierarchy (left panel of Fig. S6h), reproducing the experimental result in the resting state (Chaudhuri et al. 2015; Li and Wang 2022).

Why do areas high in the hierarchy, with long time constants in the resting state, now exhibit short time constants? To gain an intuition for this surprising observation, imagine that each area could be described by an effective energy potential with its minimum corresponding to the firing rate of persistent activity (schematically illustrated in Fig. 4i). Following a brief perturbation, the time constant quantifies the speed at which the system returns to the local attractor. The potential gradient is steep for an area near the top of the hierarchy that displays very stable mnemonic activity, thus with short time constants. By contrast, it would be very shallow for an area near the transition that recovers slowly from perturbations (Tredicce et al. 2004). Although mathematically incorrect because areas are not isolated from each other, this simplified explanation helps comprehend the inverted-V shaped pattern of timescales along the cortical hierarchy during working memory.

#### Normal form analysis of bifurcation in space

We observed (Fig. 5a) that the firing rate gap shown in Fig. 4g depends on the gain parameter *d* of the input-output transfer function of our model as Eq. (8) (see Methods): it is nearly zero when *d* = 0.157, increases progressively with *d* and saturates at large *d* values where the transfer function becomes threshold-linear (see Methods, Eq. (9)) (Abbott and Chance 2005). With *d* = 0.157, in the spatial profile of persistent activity firing rate is constant and low in areas below a hierarchical location, at the onset of a spatial bifurcation, then starts to increase (without a discrete jump) in areas higher in the hierarchy, reminiscent of a second-order phase transition in statistical physics (Landau and Lifshitz 2013). Nevertheless, the working memory state continues to be characterized by an inverted-V shaped time scale profile (Fig. 5b), suggesting that critical slowing down does not require the presence of a firing gap in the persistent activity state.

**Figure 5.**
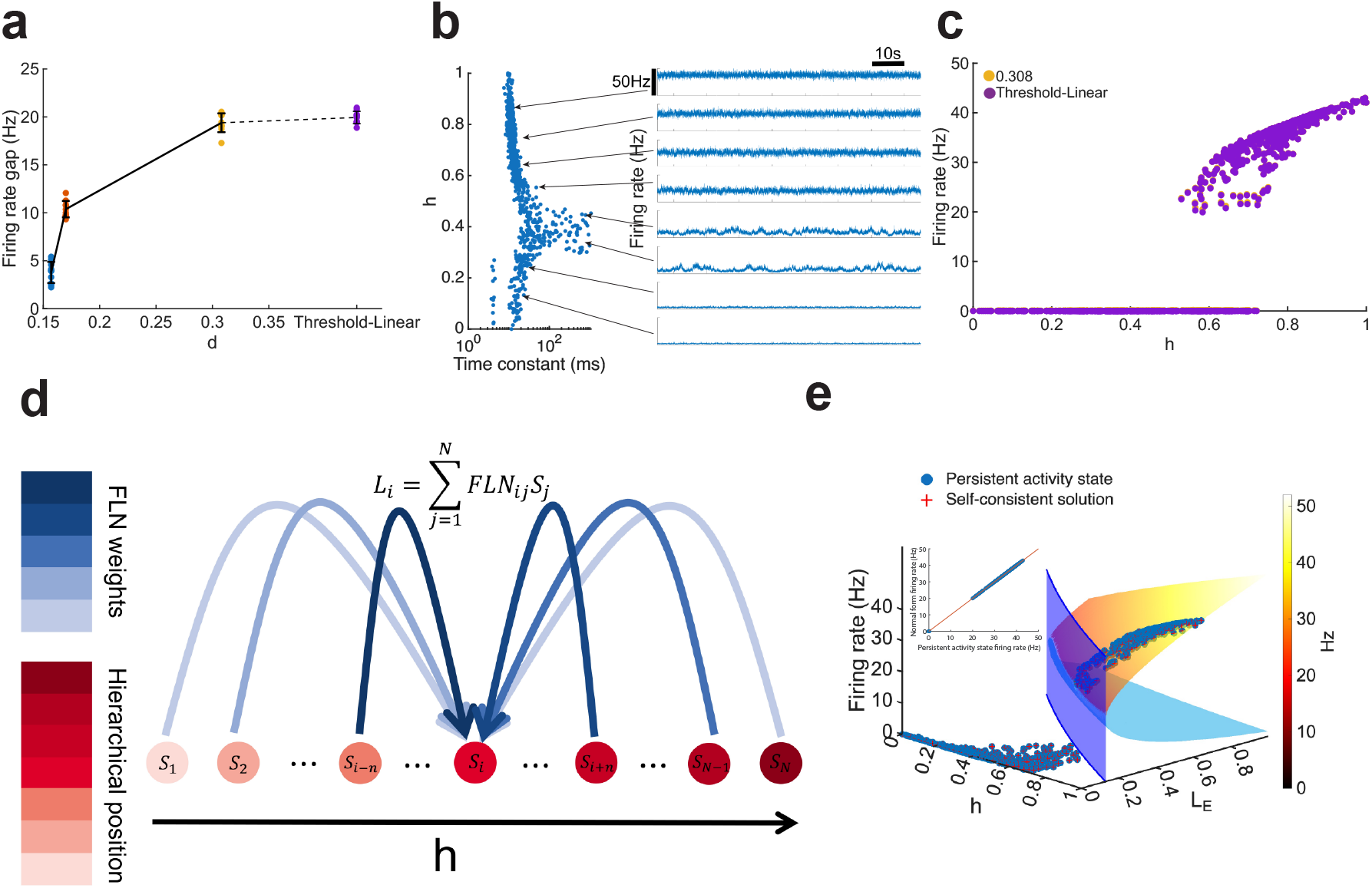
Normal form analysis that establishes mathematically the concept of bifurcation in space. (a) As the gain parameter *d* increases from 0.157 to ∞, the firing rate gap in the spatial profile of persistent activity progressively widens from zero. (b) Model behavior with the gain parameter d=0.157. When there is no rate gap, firing rates of persistent activity (right) of 8 selected brain areas show a smooth variation, yet the time constant still exhibits an inverted-V pattern (left). (c) With a sufficiently large gain parameter *d* of the input-output transfer function, persistent activity as a function of the hierarchy (orange) is similar to that of the model with a threshold-linear transfer function (purple) in the generative cortex model. (d-e) Normal form analysis with threshold-linear transfer function. (d) Illustration of the mean long-range input current to an area *i* as a sum of synaptic inputs weighted by *FLN*_*j*_ from all source areas *j*. The gradient of red and blue colors corresponds to hierarchical positions of areas and the weight of long-distance connections *FLN*_*ij*_ from area *j* to area *i*, respectively. (e) The bifurcation in hierarchical space occurs at the critical line in the plane of hierarchy *h* and weighted long-range excitatory input *L*_*E*_. Inset: The firing rate from network simulations versus the predicted firing rate from the normal form analysis shows perfect agreement.

The model behavior with large *d* values is approximately reproduced when the transfer function is threshold-linear (Fig. 5c). In this case, the model is much simplified and allowed us to undertake analytical calculation of a “normal form” according to mathematical bifurcation theory (Kuznetsov et al. 1998), in order to elucidate rigorously the transition observed in Figs. 2, 4 as a new form of bifurcation. In our model, cortical areas indexed by *i* = 1, 2, …, *N* receive long-range excitatory input currents from the network’s recurrent interactions, 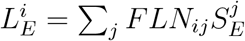 where *FLN*_*ij*_ is the connection weight from area *j* to *i* and 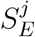 is the output synaptic variable of area *j* (see illustration in Fig. 5d). The quantity 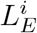 is a weighted average long-range input; if its value is known, the equation for area *i* can be solved. However, the input current for each area *i* depends on its connections (the *FLN*_*ij*_ values) and the collective activity state (all 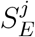 values). Indeed, the firing rates are driven by 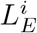, which in turn depend on the firing rates themselves; the two must be solved in a self-consistent manner for the entire system (not separately for each area). At the limit of infinite gain, equivalent to a threshold-linear transfer function (see Methods, Eq. (9)), the critical line that separates the regions with and without bistability in the *h*−*L*_*E*_ space can be calculated using Eq. (23) (see Fig. 5e, blue wall). We carried out Taylor expansion around the bifurcation line for the entire system, yielding the normal form of the bifurcation in the hierarchical space (see Methods, in particular, Eq. (29) and (33)). We solved the steady states of the normal form dynamical system Eq. (29), and identified both the stable steady states (Fig. 4, orange-yellow surface) and unstable steady states (Fig. 4, light blue surface). Using the normal form, we solved the self-consistent equation (Eq. (30)) for the *N* firing rates and *N L*_*E*_ variables. The results of the self-consistent equation match our numerical simulations perfectly (inset of Fig. 4e). In conclusion, the concept of bifurcation in space is rigorously established mathematically by the normal form analytical calculations.

To further test our model, we considered two “null” models. First, we randomly shuffled area-to-area interactions. This leads to modularity being absent and all areas participate in the persistent activity state (Fig. S8a, blue), even though the firing rate increases along the hierarchy as a result of the macroscopic gradient of excitation. There is no clear pattern for the time scale of neural fluctuations in the persistent state (Fig. S8b) or the resting state (Fig. S8c). Second, the macroscopic gradient of excitation is abolished by randomly shuffling the parameter *J* between areas. In this case, all the salient behavioral characteristics of our model disappear (Fig. S8d-f). These results demonstrate that the desired functional modularity depends critically on both the connectomic properties and the macroscopic gradient.

#### Coexistence of multiple functional modules (spatial attractors)

A multi-regional cortex model exhibits coexistence of numerous spatial attractors, each corresponding to persistent activity in a distinct subset of areas. Fig. 6a-c shows a list of coexisting spatial attractors, with a given set of fixed parameter values for the macaque, mouse and generative models, respectively. Note that spatial attractors are only defined by participating areas, independent of the number of selective neural populations within each area. We shall refer to these spatial attractors as functional modules, as they are well-defined activity patterns, and each is self-sustained among a selective subset of cortical areas.

**Figure 6.**
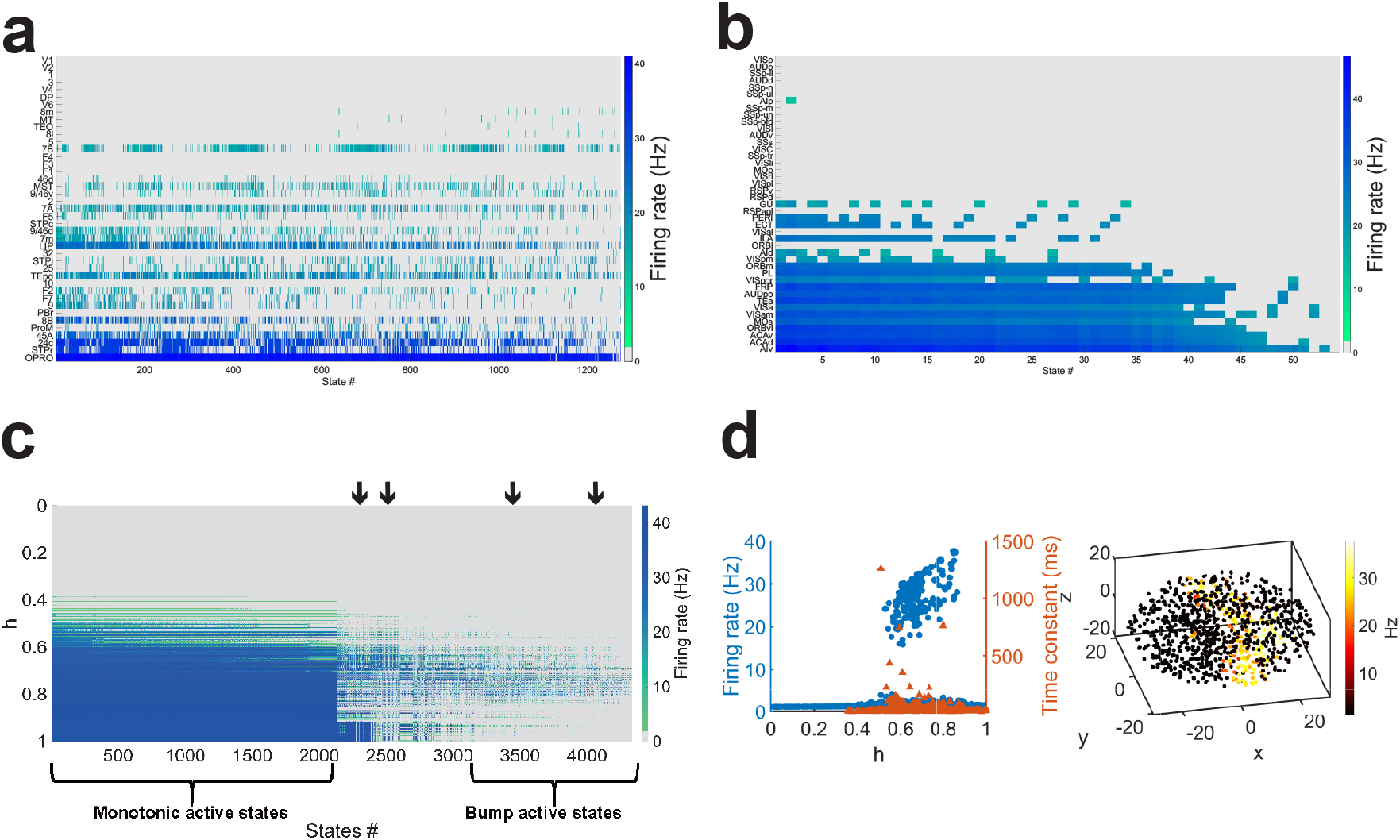
A diversity of spatially distributed attractor states. (a) A non-exhaustive list (x-axis) of coexisting self-sustained activity patterns, with activity of different areas (the hue of blue color) plotted as a function of the hierarchy (y-axis) in the macaque cortex model. (b) Similarly, the mouse cortex exhibits numerous spatial attractors. (c) More than 4,000 spatial attractors are present in the generative model. Distributed working memory states display either a monotonic pattern or a localized bump pattern with elevated activity only in the middle region of the hierarchy. The x-axis corresponds to the rank of all persistent activity states according to the number of areas with firing rates larger than 10 *Hz* (see Methods). All spatially distributed persistent states coexist in a single realization of the generative network model. (d) A localized bump spatial attractor in the cortical hierarchy of a generative model. Right: activity pattern in the 3D cortical space with firing rate indicated by color. Left: persistent firing rate (blue) and time scale of neural fluctuations (orange) along the hierarchy *h*. This state ranks 2489th among the 4333 active states in panel (c).

Interestingly, some of these self-sustained states do not show monotonic increase of firing rate along the hierarchy as in Fig. 4g (see also Fig. S9a left panel). Instead, they display a localized *bump* of activity in hierarchy space (Fig. 6d, and Fig.S9c, left panel). Strikingly, unlike the monotonic active states where persistent firing roughly follows the caudo-rostral axis of the model ellipsoid, localized bump active states generally display more scattered spatial patterns of persistent activity (Fig. 6d and Fig. S9c, right panel). The time constant of each area’s neuronal fluctuations in those localized bump states is maximal and exceptionally long at both edges of the localized bump (Fig. 6d and Fig. S9c). This double-peaked pattern of timescales further demonstrates critical slowing down as a signature of bifurcation in space, which in the case of a localized bump attractor takes place at two locations.

An interesting question is whether *functional modules* defined here can be predicted by *structural modules* defined by network analysis of connectomic data (Sporns 2014; Fornito et al. 2016). In graph theory, structural modules are identified as communities corresponding to mutually exclusive subsets of nodes (cortical areas) with many more connections among themselves than with the rest of the graph. To assess the relationship between functional modules defined by spatial attractors and structural modules, we quantified the latter by applying the Louvain community detection algorithm (Blondel et al. 2008) to the connectivity matrix, revealing 11, 5 and 4 structural modules for the generative model (Fig. 7a), macaque model (Fig. 7b) and mouse model (Fig. 7c), respectively. Note that for the macaque model, the 5 communities are identified only for the subgraph of 41 areas (out of 91 parcellated cortical areas (Markov et al. 2014)) for which the FLN data are currently available. Fig. 7d-f illustrate how relations between a functional module and 5 structural modules are quantified. For each of two sample spatial attractors (upper and lower plots of Fig. 7d), a functional module consists of areas with persistent firing rates above a threshold of 10 Hz. In Fig. 7e, areas (x-axis) are divided according to their memberships to each of 5 communities (structural modules) (y-axis), shown in green or purple depending on whether they belong to a given spatial attractor (functional module). This way, a structural module is described by a binary vector of areas (0 or 1) defined by whether each area belongs to a given community; a functional module is described by a binary vector of areas (0 or 1) depending on whether each participates in a given spatial attractor. The Hamming distance between the two (Fig. 7f) defines their similarity. For instance, all members of community 3 are part of the attractor 1, therefore, the Hamming distance is 0; for the attractor 2, no member of community 1 shows persistent activity, and the Hamming distance is 1. As is clear in these two examples, a spatial attractor typically includes areas in multiple communities.

**Figure 7.**
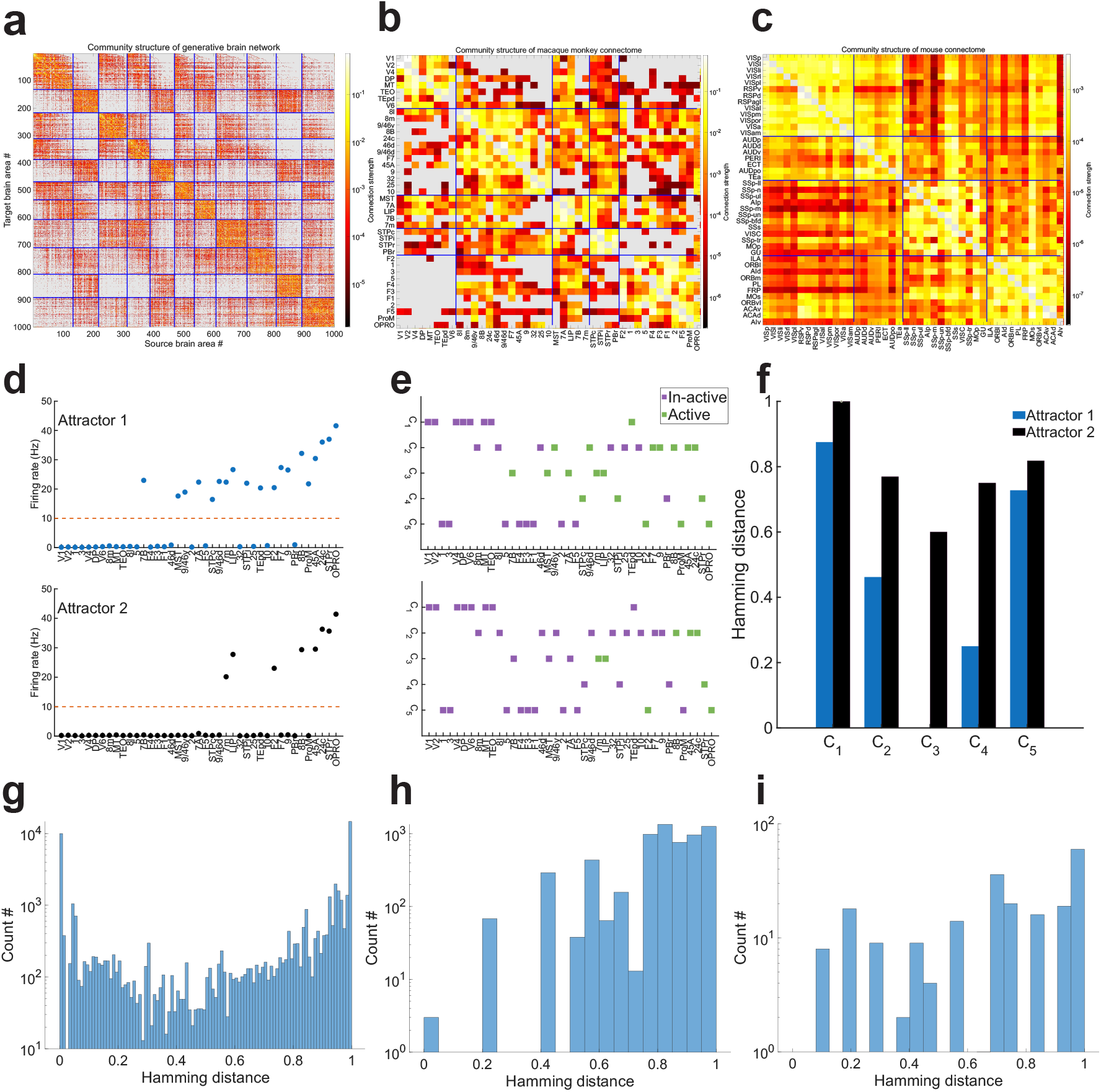
Comparison between functional and structural models. (a-c) Structural modules are defined by the Louvain community detection algorithm applied to the connectivity matrix for the generative model (a), macaque model (b) and mouse model (c). (d-e) Hamming distance between a functional module and each of five structural modules (communities) in the macaque cortex model. (d) Mnemonic persistent activity in the hierarchical space for two sample spatial attractors. A functional module consists of areas with firing rate above a threshold of 10 *Hz* (dashed horizontal line). (e) Membership to structural modules (y-axis) of individual cortical areas (x-axis). Green indicates participation in the functional module. (f) Hamming distance between a functional module and each of the five structural modules. (g-i) Histogram of Hamming distances of all functional modules (spatial attractors in the top row) with structural modules for the generative model (g), macaque model (h) and mouse model (i). There is no simple relationship between communities and spatial attractors, as functional modules are determined by complex recurrent neural circuit dynamics in the multiregional cortex, which cannot be inferred from structural modules alone.

**Figure 8.**
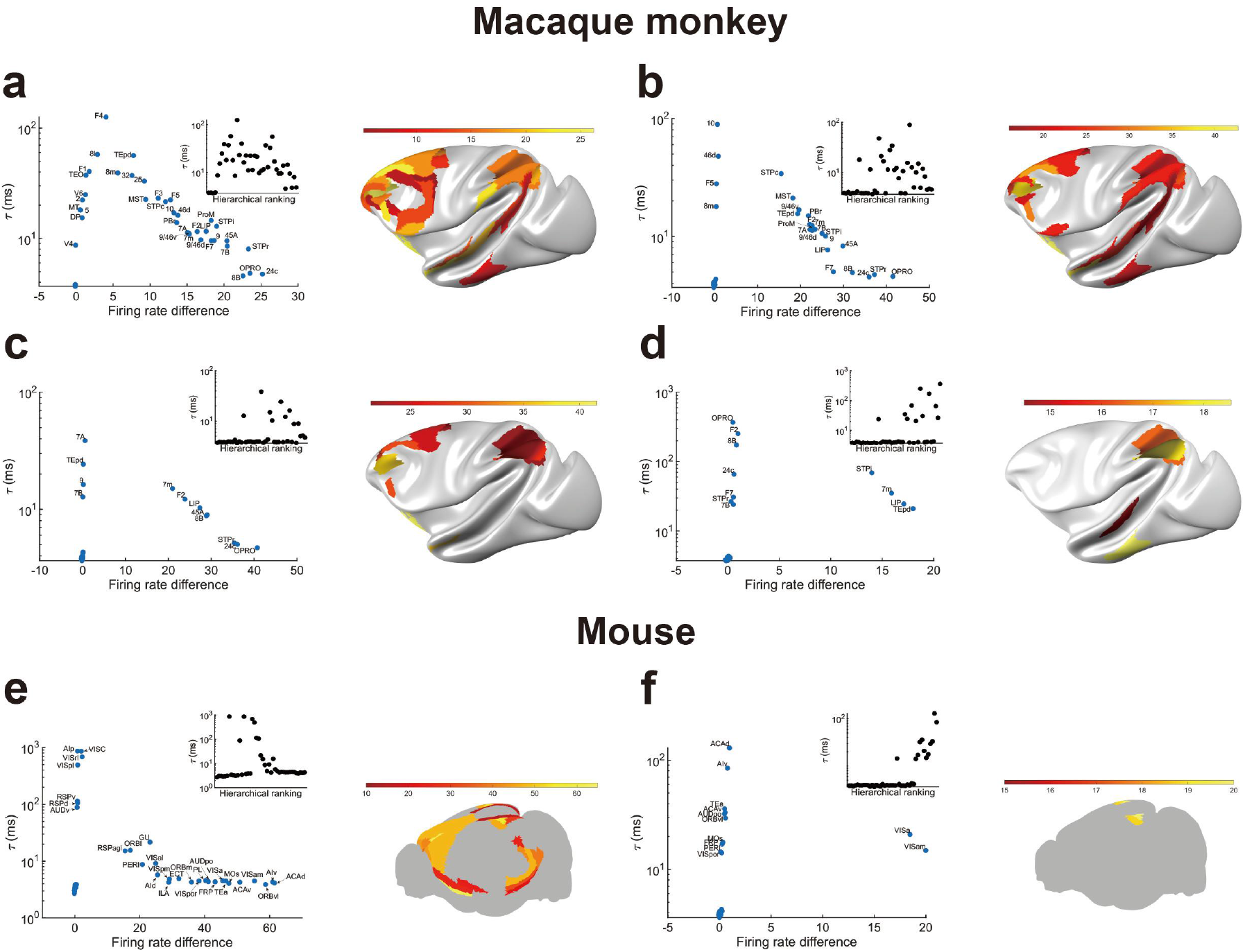
A plethora of bifurcations in space coexisting in a single cortical system. (a-d) Four sample spatial attractors of a macaque cortex model; (e-f) two sample spatial attractors of a mouse cortex model. Right: activity map showing a subset of areas that defines a functional module. Left: time scale of neural fluctuations as a function of the differential firing rate with respect to the resting state, with a peak at relatively small firing rate differences indicative of critical slowing down as a manifestation of bifurcation in space associated with an individual spatial attractor. Inset: timescale as a function of hierarchy.

Figure 7g-i displays the histograms of Hamming distances between spatial attractors and communities for the generative model, macaque and mouse connectome-based models, respectively. A peak around 0 for the generative model is indicative of numerous spatial attractors that engage all areas of some communities; but this peak is absent for the macaque and mouse models, which means that when a structural module partakes in a functional module, only some but not all of its member areas are involved. The wide distributions demonstrate that structural modules are insufficient to account for functional modules that are generated by complex largescale cortical dynamics. Likely, persistent activity patterns depend on reverberatory interactions through short as well as long loops among several areas that are not captured by communities defined by monosynaptic interareal connections.

#### A diversity of bifurcations in space

Fig. 8 shows 4 sample spatial attractors of the macaque cortex model and 2 spatial attractors of the mouse cortex model, each engaging a subset of areas (right panels: activity map). The time scale *τ* of neural fluctuations is plotted as a function of the difference in the firing rate *δr* = *r*_active_ − *r*_rest_ between the persistent activity *r*_active_ and baseline resting state *r*_rest_, revealing a similar pattern (left panels, blue), while its dependence on the hierarchy varies considerably from one to another (inset, black). Importantly, long time scales correspond to relatively small firing rate differences which correspond to a spatial location separating those areas not involved in the functional module (with *δr* ≃ 0) from those that are, demonstrating bifurcation in space in each of the 6 examples. This plot can be used to uncover a bifurcation in space even when the estimated time scale from neural activity displays complex variations along the cortical hierarchy.

In summary, bifurcation in space is defined separately for each of numerous spatial attractors of distributed persistent activity. In other words, a single network has many bifurcations in space, each for the emergence of a subset of areas (a functional module) engaged in the corresponding persistent activity state and manifested by critical slowing down at its transition spatial location. A *diversity* of time scale patterns along the cortical hierarchy highlights a rich repertoire of observable manifestations of bifurcations in space.

## Discussion

Motivated by the experimental advances, theoretical modeling of a multi-regional cortex has come to the fore (Perich and Rajan 2020; Wang 2022). Connectome-based models were developed for distributed working memory (Froudist-Walsh et al. 2021; Mejias and Wang 2022; Ding et al. 2024), and more recently are beginning to be applied to perceptual decision-making (Zou et al. 2023; Klatzmann et al. 2025).

The present work aims at uncovering a specific mechanism for the emergence of functional specialization, which is highly non-trivial under the assumptions that cortical areas share a common architecture but differ quantitatively from each other and that the cortical connectome exhibits an abundance of long-range inter-areal projections. The main findings are fivefold. First, our macaque cortex model incorporates new connectivity data of the area MST and reproduces the experimental observation of a sharp emergence of mnemonic persistent activity between the monosynaptically connected MT and MST areas (Mendoza-Halliday et al. 2014), validating the model. It offers a large-scale circuit account of choice correlates in sensory areas like MT that result from top-down signaling (Wimmer et al. 2015). Second, we show that a connectome-based model of macaque monkey cortex displays functional modularity for working memory and subjective choice in decision-making. In both macaque and mouse models, as well as the generative model, a transient input to the primary visual cortex triggers self-sustained persistent activity distributed across a subset of areas; timescales of mnemonic neural fluctuations display an inverted-V shaped pattern along the cortical hierarchy. Third, a mathematical analysis using the normal form theory of the generative model rigorously establishes the novel concept of bifurcation in space. The transition occurs locally in spite of the fact that the spatial attractor is a collective phenomenon arising from widespread long-range area-to-area interactions. Fourth, our model is not *à priori* prescribed with a modular network structure; instead, it is built on the measured connectomic data. We propose a quantitative similarity measure for comparison between structural modules and functional modules. Numerous spatial attractors coexist in a single cortical system, each defining a functional module which in general cannot be inferred from structural modules alone. Fifth, a plethora of bifurcations in space, each corresponding to a spatial attractor, gives rise to a variety of timescale patterns which may potentially be observed in different experiments. For example, one spatially distributed attractor could store sensory information, while another could maintain a behavioral rule that guides sensorimotor mapping, etc.

Recent reports of widespread neural representations in mice warrant a critical assessment. Hypothetically, a decision signal could be generated in one area or several selective areas, then propagates to many other areas and thus is decodable everywhere (Bondy et al. 2025). In the primates, there is ample evidence that working memory is distributed over a selective subset of areas (Leavitt et al. 2017; Christophel et al. 2017). In our models, the existence of functional modularity is not a given; it is absent when the experimentally measured connection weights are randomly shuffled (Fig. S8)). Also, parameter changes that enhance inter-area excitatory reverberations could give rise to an “everything everywhere” scenario, in which persistent activity during a mnemonic delay engages all areas (Fig. 3a and Fig. 3b). We examined how functional modularity may emerge and found that the mechanism is mathematically described as a bifurcation occurring at a transition location in the spatially embedded cortex. It it worth noting that this phenomenon should be distinguished from the idea of “computation near criticality”, which refers to critical dynamics for an entire neural system that requires fine-tuning parameters (Shew and Plenz 2013). By contrast, a bifurcation in space is characterized by criticality locally for areas close to the transition, and is robust against parameter changes, which would merely move the spatial location of bifurcation.

What are the implications of this theoretical work for experimental studies of distributed representations and functional modularity? (1) A cortex may or may not display functional modularity, possibly depending on animal species and task requirements. As a signature of functional modularity, the surprising and specific prediction of an inverted-V pattern of timescales (Figs. 2–4) is directly testable with recordings from many cortical areas in a working memory task. (2) Counterintuitively, certain cortical areas including the prefrontal cortex exhibit short time constants as an indication of robust mnemonic activity against perturbations (Suzuki and Gottlieb 2013; Murray et al. 2017; Froudist-Walsh et al. 2021). This raises the possibility that while playing a major role in sustaining working memory representations, they may be sensitive only to sufficiently strong optogenetic inactivation in behaving animals. (3) In our model, during a mnemonic delay, areas engaged in stimulus-selective persistent activity send sustained top-down synaptic inputs to both excitatory and inhibitory neurons in sensory areas that do not show noticeably elevated spiking activity. Since Blood Oxygen Level Dependent (BOLD) signals reflect directly synaptic currents rather than neural spiking activity (Logothetis et al. 2001), our work offers an alternative interpretation of the observation (which has been interpreted as evidence against functional specialization) that the information content of working memory can be decoded in primary sensory areas such as V1 by fMRI measurements (Harrison and Tong 2009; Sreenivasan and D’Esposito 2019). (4) An inverted-V shaped timescale profile during decision processes would explain why areas at intermediate hierarchical levels are particularly suitable to slow time integration of evidence in favor of choice options, whereas areas near the top of the hierarchy operate in a winner-take-all manner leading to a categorical decision (Gold and Shadlen 2007; Brody and Hanks 2016; Murray et al. 2017). (5) When structural and functional data are both available, for instance with structural connectivity measured by diffusion tensor imaging and resting state functional connectivity from fMRI, the functional and structural modules can be compared using the similarity measures introduced in Fig. 7. (6) In a specific experiment and during a particular behavioral epoch, the timescale may display a complex pattern across the hierarchy (e.g., Khilkevich et al. (2024)). In such cases, a plot as a function of the differential firing rate with respect to a baseline (Fig. 8) provides a test for revealing a bifurcation in space associated with the emergence of functional specialization required for ongoing behavior. (7) We demonstrated distributed coding and processing under the condition that they emerge from interareal interactions as a clear case of collective phenomenon. Alternatively, intrinsic recurrent excitation may be sufficiently strong in some areas in isolation to subserve decision-making and working memory (Wang 2002), which we considered in model simulations (Mejias and Wang 2022). Future experiments and computational work are needed to differentiate these two scenarios.

There is no contradiction between the inverted-V shaped pattern of time constants during working memory and our previous report that the dynamical timescale roughly increases along the cortical hierarchy during a resting state(Chaudhuri et al. 2015; Wang 2022). This is because time constants are uniquely defined mathematically only for a linear dynamical system; they critically depend on the internal state under consideration in a strongly recurrent neural circuit described by a highly non-linear system. The present work thus extends our previous finding of a hierarchy of time constants. We propose that an inverted-V shaped pattern of time constants during working memory represents a sensitive test of the absence or presence of functional modularity. Testing this model prediction warrants the following considerations. First, as noted above, time constant estimates may vary in different behavioral epochs and tasks. Second, there is heterogeneity of time constants across single cells within an area, therefore sufficient statistical power is needed for a cross-area comparison (Cavanagh et al. 2018; Soltani et al. 2021; Shi et al. 2025). The new prediction is that the *average* of time constants over recorded neurons in each area displays an inverted-V shaped pattern during working memory. Third, a mnemonic delay period must be much longer than to-be-assessed autocorrelation time constants. Fourth, critical slowing down is manifested near a spatial location, but the number of recorded cortical areas is limited; it remains to be seen in experiments how close one can get to a bifurcation locus in the cortical system.

This work focuses on spatial patterns of modular neural representations that are mathematically described as attractor states. The concept is not limited to steady state attractors, however. Generally, attractors can display complex temporal dynamics such as chaos rather than steady states (Strogatz 2016), and the attractor paradigm can be consistent with considerable temporal variations of neuronal delay period activity and cell-to-cell heterogeneities (Baeg et al. 2003; Stokes et al. 2013; Murray et al. 2017; Miller et al. 2018; Barbosa et al. 2020; Wang 2021; Panichello et al. 2024). Future research is needed to extend the concept of bifurcation in space beyond steady states.

A limitation of this study is the consideration of the neocortex alone. A previous work (Ding et al. 2024) examined the impact of thalamic inputs into a thalamocortical mouse model, suggesting that the thalamocortical connection loop effectively enhances corticocortical connections. Neural correlates of working memory and decision making are present in subcortical structures including the thalamus, basal ganglia, amygdala and hippocampus (Guo et al. 2017; Khilkevich et al. 2024; Chen et al. 2024; International Brain Laboratory et al. 2025). The quantitative connectomic analysis of the interplay between cortical and subcortical brain regions is still fairly incomplete and warrants future efforts. A combination of neurophysiological experiments and computational modeling would be required for advances in this direction.

In contrast to local interactions through diffusion or chemical reactions, interareal cortical interactions involve long-range connections, making it all the more remarkable that criticality can occur in a spatially restricted fashion in a multi-regional cortex. A *recurrent* deep neural network (a hierarchical cortex with numerous feedback loops) and macroscopic gradients are sufficient to generate multiregional functionally modules in our model. Indeed, we found that bifurcation unfolds in space and time, underlying subjective judgment in perceptual decision-making (Figure 1). Research along these lines should broadly help us understand the emergence of novel functional capabilities such as reasoning, mathematics and music, that are instantiated in certain parts of the brain as a result of quantitative changes of properties, providing a mechanistic foundation for functional modularity in the brain.

## Acknowledgments

We thank Jorge Mejias and Xingyu Ding for help with the cortical model codes of the macaque and mouse, respectively. We thank Loïc Magrou for help with macaque brain maps.

## Author contributions

X.-J.W. was responsible for the concept of bifurcation in space, designed the project, was actively involved in all details throughout the work, and wrote the bulk of the main text. X.-J.W., J.J., and U.P.-O. designed modeling; J.J., R.Z., U.P.-O., and A.B. performed the research with supervision and inputs from X.-J.W.; J.V. and H.K. performed the histological and anatomical analysis of the MST data; all the authors contributed to writing the paper.

## Data and materials availability

All data and codes can be accessed at https://zenodo.org/records/19150862, and will be made publically available upon publication.

## Materials and Methods

### Anatomical data and numerical parameter sets of the connectome-based cortical model of macaque and mouse

The macaque connectome data covering 41 cortical areas are presented in Fig. S1a. Besides the latest macaque connectome adopted in Froudist-Walsh et al. (Froudist-Walsh et al. 2021), we incorporated previously unreported measured connections of the MST cortical area. This is significant because the MST region of the visual system receives direct monosynaptic inputs primarily from area MT. The former not the latter, exhibits persistent activity during a mnemonic delay (Mendoza-Halliday et al. 2014). Following the injection of a retrograde tracer into the target cortical area MST, we precisely quantified retrogradely labeled neurons throughout the brain (excluding labeled neurons in the injected target area) and assigned them to supra- and infra-granular layers of all the source areas projecting to the target, as described previously (Froudist-Walsh et al. 2021; Markov et al. 2014). This allowed us to define two important metrics: i) the fraction of labeled neurons (FLN) representing the normalized strength of the projections from source areas to the target area - MST, and ii) the proportion of supragranular labeled neurons (SLN), indexing hierarchical distance between all connected cortical areas relative to the target - MST. SLN is a continuous variable that gradually varies between 1 for long-distance feedforward projections and 0 for long-distance feedback projections, for short-distance projections both feedforward and feedback tend to 0.5. Additionally, the hierarchical position of each brain area can be estimated based on the feedback and feedforward connections among the 41 brain areas (Markov et al. 2014). In brief, we used a beta-binomial model, with SLN values of each measured connection of the 41 areas dataset as input, to determine hierarchical levels of all areas based on the model coefficients as detailed in (Vezoli et al. 2021). Importantly, we use the number of neurons in the model, thus generating a weighted hierarchy. The overall FLN values are shown in Fig. S1a. The resulting hierarchical positions are shown in Fig. S1b.

For the cortical model of the macaque monkey model, we systematically analyzed how model behavior depends on two parameters: 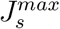 that denotes the largest strength of local excitatory-to-excitatory connections among cortical areas, and *Z*, which controls the counterstream inhibitory bias (Chaudhuri et al. 2015; Wang 2022) (the smaller the *Z*, the more top-down projections target inhibitory versus excitatory neurons in a recipient area). We found that, in the 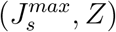 plane (Fig. S3a), the parameter set that satisfies the requirements for decision-making tasks constitutes a subset of the parameter space that meets the criteria for the working memory tasks (Fig. S3b,c). Consequently, we selected the parameter set that is tailored to the decision-making task and inherently satisfies the requirements for the working memory task as well. The “reference parameter” values of 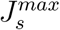 and *Z* were chosen so that the model’s performance fits well with the monkey’s psychometric function (Fig. 1c).

All the other parameters are the same as in Mejias et al. (Mejias and Wang 2022), except the 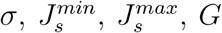, and *Z*. For Fig 1, Fig 2 a-c, d upper panel, Fig. 8a, Fig. S2a-e, and Fig. S4, *σ* = 24*pA*, 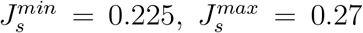, *G* = 0.48, and *Z* = 0.82. For Fig 2d lower panel, *σ* = 44*pA*, 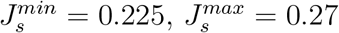, *G* = 0.48, and *Z* = 0.82. For Fig 5b,k, *σ* = 0*pA*, 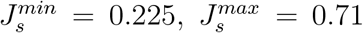, *G* = 0.48, and *Z* = 0.25. For Fig. 5g, upper panel, and Fig 8b, *σ* = 0*pA* and *σ* = 16*pA*, respectively. The other parameters’ are 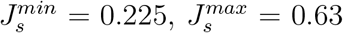, *G* = 0.48, and *Z* = 0.42. For Fig. 5g, lower panel, and Fig 8c, *σ* = 0*pA* and *σ* = 18*pA*, respectively. The other parameters’ are 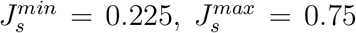, *G* = 0.48, and *Z* = 0.27. For Fig 8d, *σ* = 10*pA*, 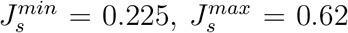, *G* = 0.48, and *Z* = 0.27. The AMPA noise strength for excitatory populations 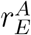 and 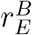 is the same. In Figure 3a, the parameters are the same as in Fig. 2a-d, except *σ* = 10*pA*, 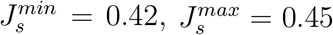, *G* = 0.3, and *Z* = 2. In Figure 3b, the parameters are the same as in Fig. 2e-f, except *σ* = 4*pA, G* = 0.05, and *Z* = 2.

For the RT task simulations (Fig. 1) where evidence favored option A, the RT was recorded once Δ(*t*) hit the positive threshold for choice A or the negative threshold for choice B. The threshold value was adjusted (to 0.525) for the model to capture RTs comparable to those observed in monkey experiments (Mazurek et al. 2003). The psychometric function is fitted with a Weibull function. Reaction times for correct and error trials are computed separately. Time-to-threshold values and activity maps are calculated from the average of the 500 correct and error trials.

The mouse connectome data covering 43 cortical areas shown in Fig. 2, Fig. 7c, and Fig. 8 are the same as those in Ding et al. (Ding et al. 2024). Besides, the model settings are the same, except for the *σ, g*_*E,self*_, *µ*_*IE*_, and *β*. For Fig. 2f, g, h upper panel, Fig. 8e, and Fig. S2f-g right panel, the parameters set are exactly the same as Ding et al. (Ding et al. 2024), except the *σ* = 70*pA*. For Fig. 5c,l, we choose *σ* = 0*pA, g*_*E,self*_ = 0.45, *µ*_*IE*_ = 0.25, and *β* = 1.4. For Fig. 8f, we choose *σ* = 2*pA, g*_*E,self*_ = 0.45, *µ*_*IE*_ = 0.25, and *β* = 1.4. The AMPA noise strength of excitatory populations 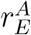 and 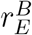 is identical.

### Auto-correlation function of excitatory firing rate and estimated time scales

We calculate the auto-correlation function of each cortical area based on the excitatory firing rate time series. The sample rate and total length of the firing rate time series are 200 *Hz* and 80 seconds, excluding the transient period. First, we compute the autocorrelation function using the *autocorr* function in MATLAB, with the maximum lag set to 50 seconds (corresponding to 10000 sample points). After that, we estimate the time scale of the brain area based on the auto-correlation function. Since the auto-correlation function could have more than one time scale, we fit the auto-correlation using both the single-exponential and double-exponential functions, which show as follows:

Single-exponential function:

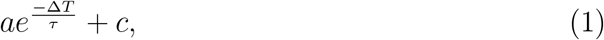

Double-exponential function:

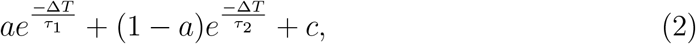

where Δ*T* is the time lag of the auto-correlation function, *τ* is the estimated time constant of a brain area with the single-exponential function, *τ*_1_ and *τ*_2_ are the two estimated time constants of each brain area with the double-exponential function. For the double-exponential function, we define a combined time constant *τ*_*c*_ = *aτ*_1_ + (1 − *a*)*τ*_2_. However, if *a <* 0.07 or *a >* 0.93, we will choose *τ*_2_ or *τ*_1_ as the final time constant of double-exponential fitting, respectively. Otherwise, we choose *τ*_*c*_ as the final time constant of the double exponential fitting. We fit the auto-correlation function with the single-exponential and double-exponential functions using the *fit* function in Matlab. For the *fit* function, we set the upper and lower bound for each parameter as *a* ∈ (0, 1), *τ* ∈ (1, ∞), *τ*_1_ ∈ (1, ∞), *τ*_2_ ∈ (1, ∞), *c* ∈ (−1, 1), the algorithm of fitting procedure is Levenberg-Marquardt.

In general, we determine each brain area’s final time constant based on the fitting’s root-mean-square error (RMSE). If the RMSE of the single exponential fitting is larger than two times the RMSE of the double exponential fitting, we choose the double exponential fitting *τ*_*c*_ as the final time constant of the brain area. Otherwise, we choose the single exponential fitting *τ* as the final time constant of the brain area. On the other hand, each double exponential fit exhibits a rapid temporal component approximately at 4 *ms*, aligning with the temporal dynamics characteristic of AMPA noise of the cortical model of the macaque monkey (Fig. S2d). Consequently, we disregard this rapid component and focus solely on the prolonged temporal scale in Fig. 2d,i in the main text. However, for some brain areas, such as *V* 1, the weight of the fast time scale is greater than 0.93, which means that this brain area only contains a fast time scale. For those brain areas, we will use this fast time scale.

### A generative model for the mammalian cortical connectivity

We use the model in (Song et al. 2014) to generate multiple cortical network realizations. In this model, we randomly choose the center of *N* brain areas in a three-dimensional ellipsoid. The ellipsoid is parcellated into *N* areas through a Voronoi partition. Axon growth starts at a randomly chosen source in the ellipsoid. The direction of growth is determined by the summing force of all area centers. Growth length is randomly chosen from an exponential distribution used for modeling the reported distance effects on the connectivity (Ercsey-Ravasz et al. 2013). Since the axon’s source, direction, and length are determined, we can locate the axon’s target position in the ellipsoid. We then add a connection from the source area to the target area and repeat the axon growth process *N ×* 2.1978 *×* 10^4^ times. With this procedure, the generative network exhibits an in-degree and out-degree distribution similar to that of the real macaque brain network (whose data were obtained via retrograde tract-tracing), as well as a matching triad distribution. Additionally, based on previous studies (Song et al. 2014), a three-dimensional ellipsoid fits the connectivity and motif distribution better than a two-dimensional spheroid. Moreover, the ellipsoid is a useful approximation to cortical geometry, reflecting a simple but meaningful structure that effectively captures the hierarchical organization of brain areas, such as the rostro-caudal gradient observed in the cortex.

### Diffusion map method for the embedding of generative mammalian cortical connectivity

We analyze the connectivity generated using the diffusion map method (Coifman and Lafon 2006). This class of non-linear dimensionality reduction has recently been applied to human and macaque monkey connectomes (Margulies et al. 2016). Briefly, this method assumes a hypothetical *diffusion process* on the nodes of the symmetric version of the generative network connectivity (that is, the matrix *FLN*). The symmetry of the network ensures the equilibrium and reversibility of the diffusion process. This diffusion process generates a *diffusion metric space* where the distance between the cortical areas is defined. In this diffusion space, closer areas share more loops, which connect them with stronger connections. On the other hand, areas further apart in diffusion space share fewer loops and weaker connections. When this method is used on connectivity, the cortical network is embedded in a few “principal gradients” of the diffusion process. These principal gradients are the principal components of the normalized graph Laplacian of the diffusion process. This process leads to embedding the connectivity matrix into a low-dimensional space. Its dimensionality is determined by the selected number of principal gradients (three for Fig. 4c). We applied this method in the symmetric version of the *FLN* matrix *L* = *FLN* + *FLN* ^*T*^.

We closely followed the method described in (Coifman and Lafon 2006). First, we define the following normalized matrix

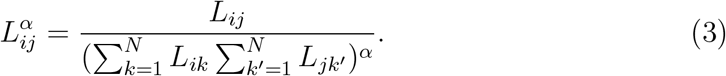

Then, we obtain the Markov transition matrix of the hypothetical Markov process on the connectivity as

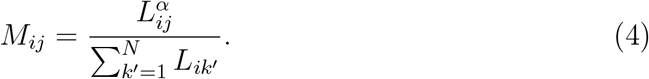

For the discrete Markov process defined on the connectivity at any time, *t*, the probability of jumping from the edge *j* to edge *i* is given by the matrix 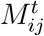. To define the principal gradients, we perform the eigenvalue decomposition of 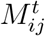, obtaining

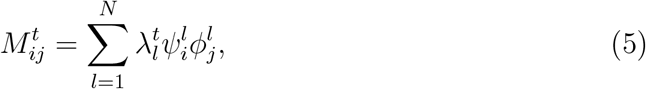

Then, the principal gradients are the set of vectors

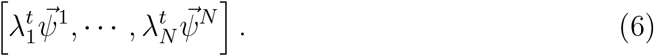

where the real eigenvalues are ordered from the largest to the smallest, with *λ*_1_ = 1, which corresponds to the stationary distribution 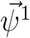 for *t* → ∞. For the dimensionality reduction presented in this work, we use the first three principal gradients 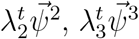, and 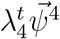. We examine the Markov process at very short time scales by taking the value *t* = 0. We choose the hyperparameter *α* = 0.5 since the underlying Markov process approximates the Fokker-Planck operator in this case (Coifman and Lafon 2006).

### Constructing cortical hierarchies from the generative mammalian cortical connectivity

We calculate two classes of hierarchies based on the three-dimensional embedding of the structural connectivity matrix through the diffusion map (see Fig. 4c): Euclidean and Hyperbolic hierarchies. To calculate either class of hierarchy value, we choose the cortical area with the smallest value in the first principal gradient as the first area in the hierarchy (origin area). This choice is arbitrary. We compute the distance in diffusion space between each cortical area and the origin area to determine the hierarchical position. For the Euclidean hierarchy, the hierarchical value of a cortical area *i* is computed using the normalized Euclidean distance 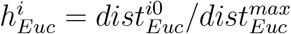. Here, the value 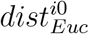 is the Euclidean distance between brain area *i* and the origin area, and the value 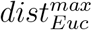 is the maximum Euclidean distance of all the brain areas to the origin area. The ranked Euclidean hierarchical positions of individual brain areas are represented by the brown circles in Fig. 4d.

For the hyperbolic distance, we estimate the distance from the origin area along the hyperbolic shape in the embedding space. To do that, we create a nearest-neighbor network for all the brain areas in the embedding space. All brain areas are interconnected when each area is linked to its neighbors within a Euclidean distance threshold *dist*^*thr*^. This threshold is defined as the maximum distance from each brain area to its nearest neighbor. The weight of each connection in the nearest-neighbor network equals the Euclidean distance between paired cortical areas. After constructing the network, we calculated the hyperbolic distance between brain area *i* and the origin by computing their shortest path. The length of the shortest path is the summation of the connection weights (i.e., Euclidean distances) within this shortest path. The shortest path finding is computed using the *dijkstra_path_length* function in the Python package of NetworkX. We define the hyperbolic hierarchical position of brain area 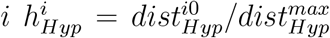, where 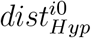 is the Hyperbolic distance between brain area *i* and the origin area. The value 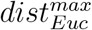 is the maximum hyperbolic distance of all the brain areas to the origin area. The ranked hyperbolic hierarchical positions of individual brain regions are marked by blue dots in Fig. 3d.

### Isolated cortical circuit of the generative model

The simplified nonlinear dynamical model is adopted from (Wong and Wang 2006), which approximates spiking neural networks with AMPA, GABA, and NMDA synapses (Wang 2002). The dynamical equations that describe the dynamics for a single cortical area are as follows:

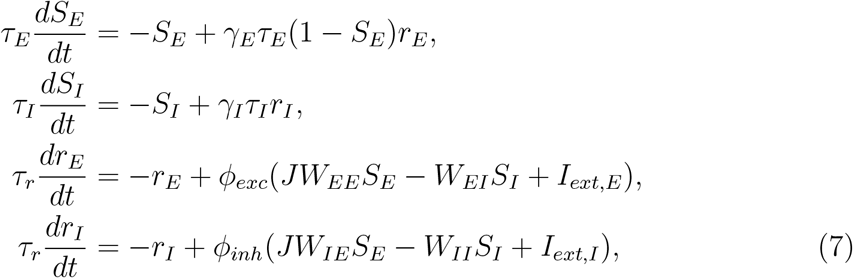

where *S*_*E*_ and *S*_*I*_ are the gating variables of the NMDA receptor of the excitatory population and the gating variable of the GABAergic receptor of the inhibitory population, respectively, the variables *r*_*E*_ and *r*_*I*_ are the mean firing rates of the excitatory and inhibitory populations, respectively. The functions *ϕ*_*exc*_ and *ϕ*_*inh*_ are the input-output transfer functions of the excitatory and inhibitory populations. The variable *J* is the cortical heterogeneity factor, which is proportional to the hierarchical value and differs for each cortical area. Unless specified, parameters are *τ*_*E*_ = 60*ms, τ*_*I*_ = 5*ms, τ*_*r*_ = 2*ms, γ*_*E*_ = 0.76, *γ*_*I*_ = 1, *W*_*EE*_ = 276.48*pA, W*_*EI*_ = 251*pA, W*_*IE*_ = 129.6*pA, W*_*II*_ = 54*pA, I*_*ext,E*_ = 329.5*pA, I*_*ext,I*_ = 260*pA*. The input-output transfer function *ϕ*(*I*) maps the average input current to a cortical circuit to the mean firing rate. We use two different functions for excitatory populations.

1. Abbott-Chance function (Abbott and Chance 2005)

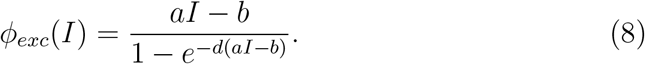
2. Threshold-linear function

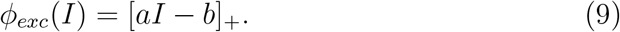

The notation [*•*]_+_ denotes rectification, i.e., *ϕ*_*exc*_(*I*) = *aI* − *b* when *aI* − *b >* 0 and *ϕ*_*exc*_(*I*) = 0 when *aI* − *b* ≤ 0.

The parameters for Abbott-Chance and threshold-linear functions are *a* = 0.27*Hz/pA, b* = 108*Hz*. The parameter *d* is the gain in the Abbott-Chance function. For a very large gain *d*, i.e., in the limit when *d* → ∞, the Abbott-Chance function becomes equal to the threshold-linear function.

For the inhibitory population, the transfer function is threshold-linear

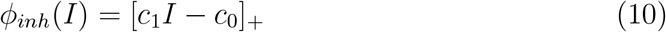

where the parameters are *c*_1_ = 0.308*Hz/pA, c*_0_ = 77*Hz*.

### Dynamical model of the generative model

We connect cortical areas with local neural dynamics described by Eqs. (7–10) using the connectivity from our generative model of the mammalian neocortex. In our model, long-range projections exist between excitatory populations (Chaudhuri et al. 2015; Froudist-Walsh et al. 2021; Mejias and Wang 2022). We detail our large-scale model below:

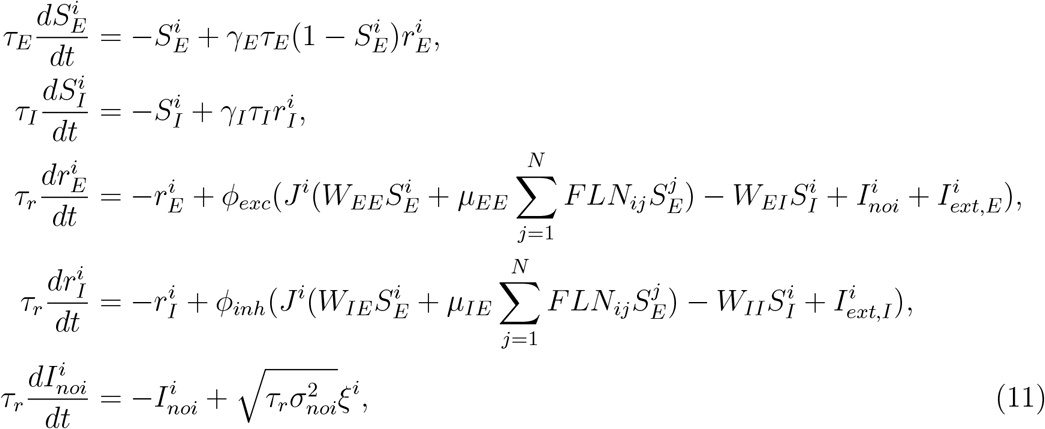

where the parameters *µ*_*EE*_ and *µ*_*IE*_ are the long-range coupling strength. Unless specified, *µ*_*EE*_ = 69.12*pA, µ*_*IE*_ = 62.809*pA*. The *FLN*_*ij*_ is the long-range connection strength from the source brain area *j* to the target cortical area *i*, generated as described in the previous section. The parameter *J*^*i*^ corresponds to the excitation gradient. This cortical heterogeneity factor scales excitation for each cortical area *i*, which is linearly related to the hierarchical position *h*^*i*^ of cortical area *i* as *J*^*i*^ = 1+*ηh*^*i*^ (see inset of Fig. 4f). We assume that all the brain areas have the same external input current 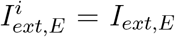 and 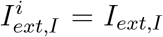. The noise term *I*_*noi*_ is an Ornstein-Uhlenbeck process, representing the AMPA synaptic noise with a short time constant *τ*_*r*_ = 2*ms* (Wong and Wang 2006). The parameter *σ*_*noi*_ is the standard deviation of the noise, and *ξ* is Gaussian white noise with zero mean and unit variance. Unless specified, all other parameters are the same as in the isolated brain area.

### Steady states for an isolated cortical area of the generative model

For solving the steady state of an isolated brain area, we set 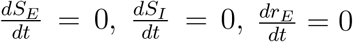 and 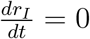. After that, we have the steady-state equations as follows:

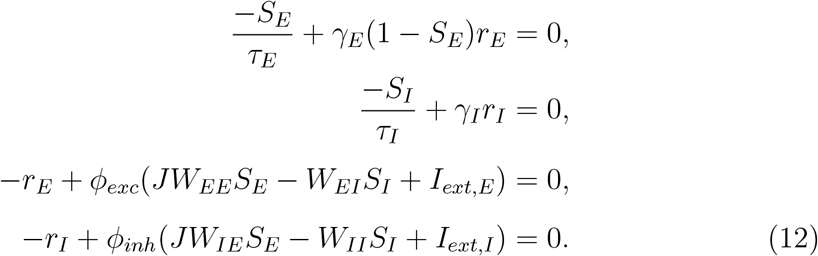

A meaningful steady state of the brain area must have positive firing rates *r*_*E*_ ≥ 0, *r*_*I*_ ≥ 0. Thus, we will have the steady state 0 ≤ *S*_*E*_ ≤ 1 and *S*_*I*_ ≥ 0. Therefore, we could reduce our steady-state equation to

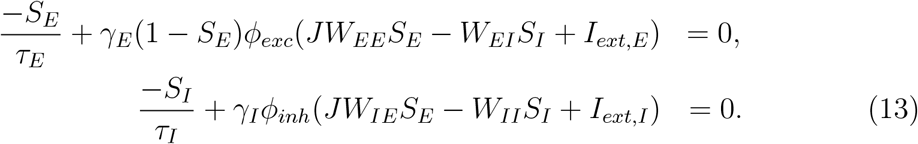

We reorganize the above expression and obtain an expression for *S*_*I*_ given by

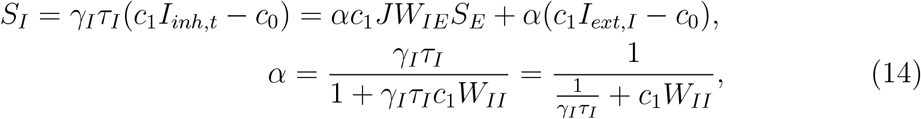

where we define *I*_*inh,t*_ as a total current input to inhibitory population, and *α* = 4.6*ms* by using the parameters of Table 1. Then, we plug-in Eq. (14) into the steady state *S*_*E*_ Eq. (13). After this manipulation, the steady state of the single cortical area is determined by the NMDA gating variable *S*_*E*_ as follows:

**Table 1.**
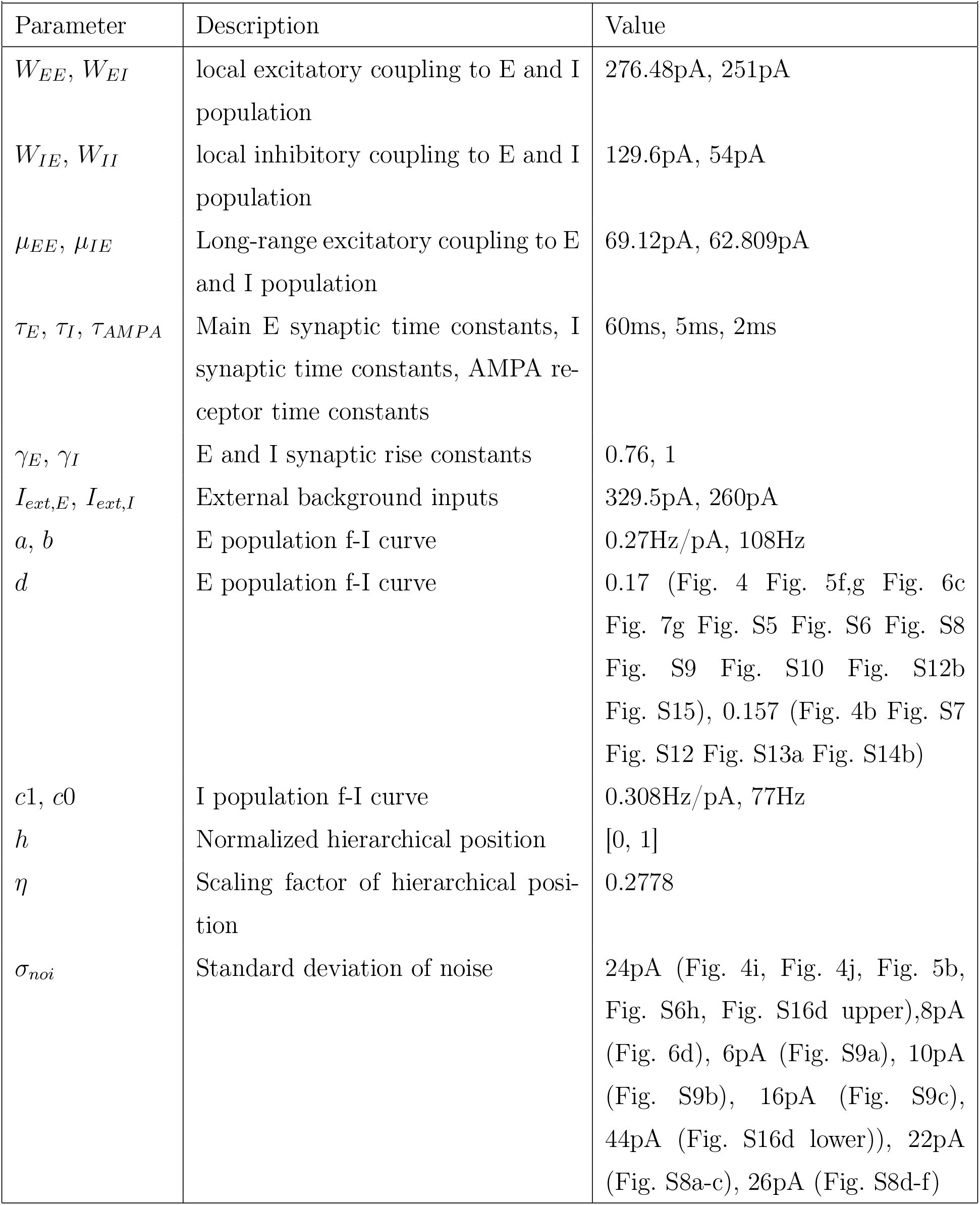
Parameters for numerical simulations of the generative model.

| Table 1. Parameters for numerical simulations of the generative model |  |  |
| --- | --- | --- |
| Parameter | Description | Value |
| $W_{EE}, W_{EI}$ | local excitatory coupling to E and I population | 276.48pA, 251pA |
| $W_{IE}, W_{II}$ | local inhibitory coupling to E and I population | 129.6pA, 54pA |
| $\mu_{EE}, \mu_{IE}$ | Long-range excitatory coupling to E and I population | 69.12pA, 62.809pA |
| $\tau_E, \tau_I, \tau_{AMPA}$ | Main E synaptic time constants, I synaptic time constants, AMPA receptor time constants | 60ms, 5ms, 2ms |
| $\gamma_E, \gamma_I$ | E and I synaptic rise constants | 0.76, 1 |
| $I_{ext,E}, I_{ext,I}$ | External background inputs | 329.5pA, 260pA |
| $a, b$ | E population f-I curve | 0.27Hz/pA, 108Hz |
| $d$ | E population f-I curve | 0.17 (Fig. 4 Fig. 5f,g Fig. 6c Fig. 7g Fig. S5 Fig. S6 Fig. S8 Fig. S9 Fig. S10 Fig. S12b Fig. S15), 0.157 (Fig. 4b Fig. S7 Fig. S12 Fig. S13a Fig. S14b) |
| $c1, c0$ | I population f-I curve | 0.308Hz/pA, 77Hz |
| $h$ | Normalized hierarchical position | [0, 1] |
| $\eta$ | Scaling factor of hierarchical position | 0.2778 |
| $\sigma_{noi}$ | Standard deviation of noise | 24pA (Fig. 4i, Fig. 4j, Fig. 5b, Fig. S6h, Fig. S16d upper), 8pA (Fig. 6d), 6pA (Fig. S9a), 10pA (Fig. S9b), 16pA (Fig. S9c), 44pA (Fig. S16d lower), 22pA (Fig. S8a-c), 26pA (Fig. S8d-f) |

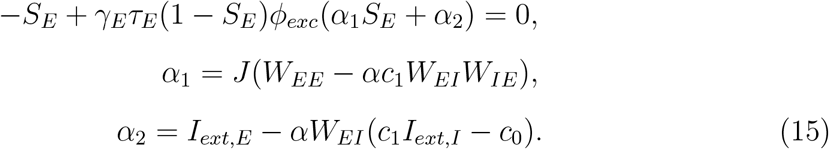

Where *α*_1_ = 230.2305*JpA* and *α*_2_ = 301.1294*pA* by using the parameters of Table 1. For the threshold-linear transfer function in Eq. (9), we immediately noticed that *S*_*E*_ = 0, *S*_*I*_ = *α*(*c*_1_*I*_*ext,I*_ − *c*_0_) is one of the steady states solution with −*W*_*EI*_(*α*(*c*_1_*I*_*ext,I*_ − *c*_0_)) + *I*_*ext,E*_ *<* 400*pA*. This solution applies to the resting state and is independent of the cortical heterogeneity factor *J*. This implies that, with the threshold-linear transfer function, the resting state consistently exists across the cortical hierarchy and satisfies *r*_*E*_ = 0.

However, for the other steady states *S*_*E*_, they obey the following quadratic equation:

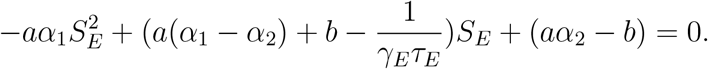

By solving the quadratic equation, we obtain two steady-state

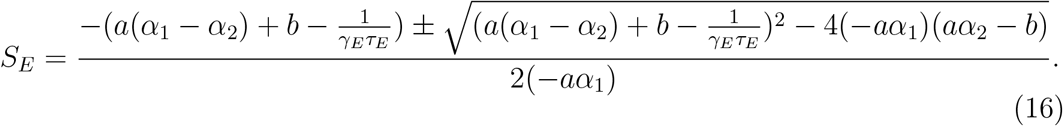

Therefore, the isolated brain area has a saddle-node bifurcation of *S*_*E*_, and the bifurcation point at 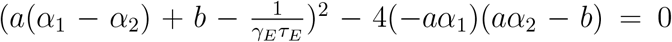. At the bifurcation point, we have 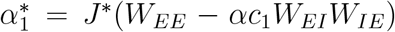 and 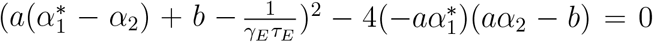, Thus 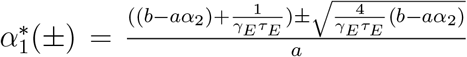. If we consider the solution 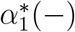 then we have that the <u>cortical</u> heterogeneity factor is given by 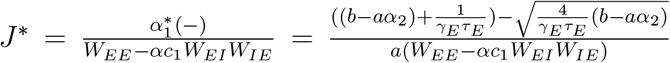. For our parameter setting, 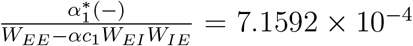, which means that *J*^∗^ ≪ 1. However, this is not possible since *J*_*min*_ = 1. Therefore, the bifurcation cortical heterogeneity factor is given by 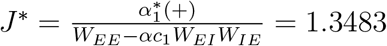.

We performed a similar analysis for the Abbott-Chance transfer function in Eq. (8). By combining Eq. (8) and Eq. (15) the steady state is given by

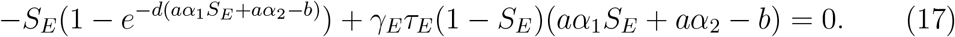

The steady state Eq. (17) is highly nonlinear, and we can not provide an analytic solution. Instead, we solve Eq. (17) using the Matlab numerical solver *fsolve*. The steady state in Eq. (17) depends on the gain parameter *d* of the Abbott-Chance function, and the bi-stable region enlarges when *d* increases. This is shown by comparing Fig. 4f, Fig. S5e, and Fig. S5f.

### Steady states of the generative model

As for the large-scale network model, we write the steady state equation as follows:

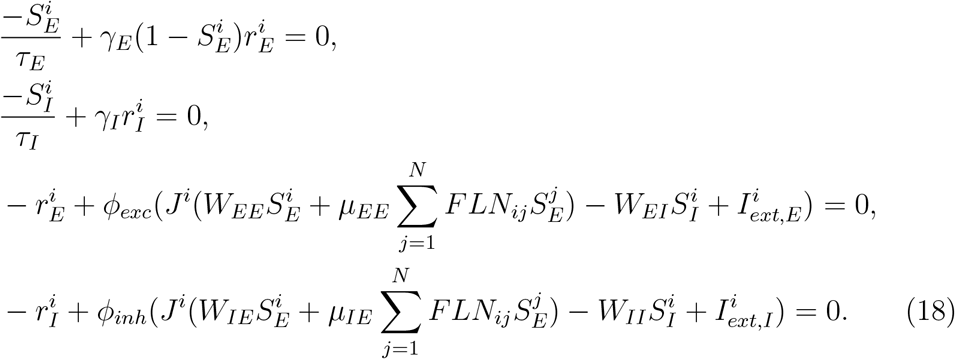

We assume that stable steady states of the large-scale network model are attractor states. Therefore, in the steady states, the long-range excitatory input for the *i*^*th*^ brain area 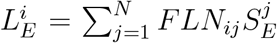 is a fixed number. The exact value of long-range excitatory inputs 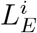 depends on the connectivity structure of *FLN*. Based on this assumption, we could rewrite the steady-state equation of the large-scale network model as

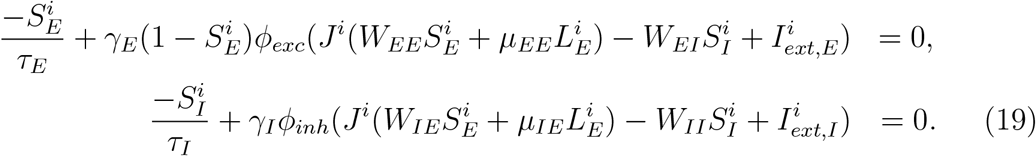

After some manipulations, we obtain the expression for 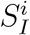 given by

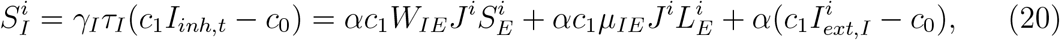

where the definition of *α* is same as in Eq. (14). Therefore, for the *i*^*th*^ cortical area the excitatory gating variable 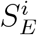 obeys the following steady state equation

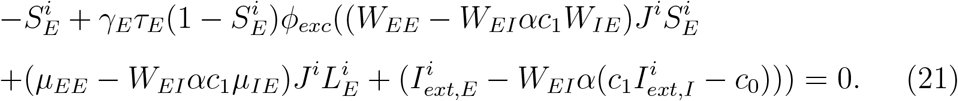

First, we analyzed the steady state Eq. (21) when the transfer function is threshold-linear. The above Eq. (21) can be written as the steady state of the following set of dynamical equations

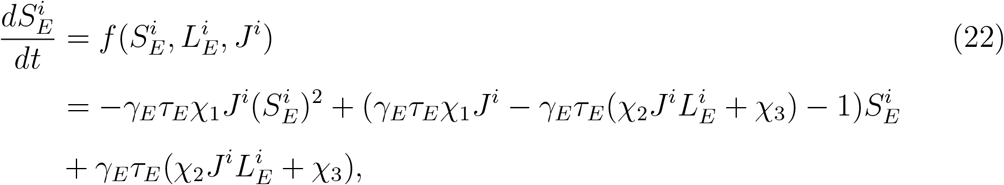

with

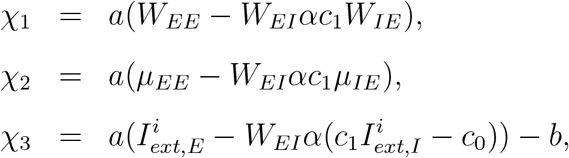

where *χ*_1_ = 62.1622*Hz, χ*_2_ = 12.6106*Hz, χ*_3_ = −19.9985*Hz* by using the parameters of Table 1, and steady states value of the synaptic variable of the *i*^*th*^ cortical area 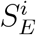 obeys the above quadratic equation equal to zero. Importantly, the steady state of 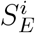 depends on the hierarchy value through *J*^*i*^ and the long-range excitatory inputs 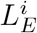.

Since the steady state equation for the synaptic variables of each cortical area 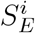 in Eq. (22) is given by a quadratic equation, the steady state can be calculated by using the quadratic formula. This calculation is similar to the steady state calculations for an isolated cortical area above (see Eq. (16)). However, in our large-scale network model, the quadratic formula of the network model is also dependent on the hierarchy value through *J*^*i*^ and the long-range excitatory inputs 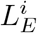. Therefore, the bifurcation happening in the hierarchical space is determined by the following expression:

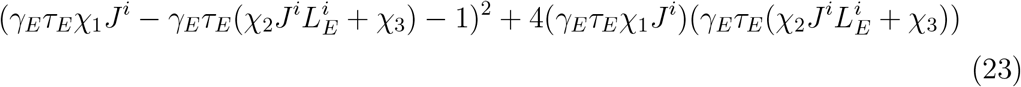

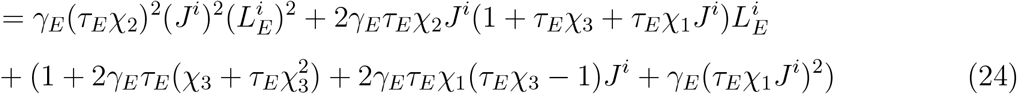

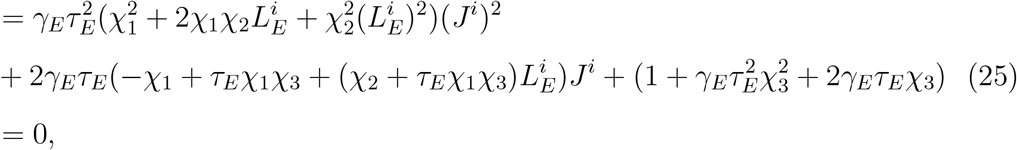

where *J*^*i*^ = 1 + *ηh*^*i*^ and 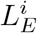 are the scaled hierarchy value and long-range excitatory inputs of *i*^*th*^ brain area, respectively. The Eq. (23) is a constraint equation in the two-dimensional space of hierarchical position *h* and weighted long-range excitatory inputs *L*_*E*_. Therefore, Eq. (23) determines where the bifurcation in space is happening in the two-dimensional hierarchy and long-range excitatory inputs space. We refer to this curve as the critical line. For example, the *i*^*th*^ brain area with scaled hierarchical value *J*^*i*^ will give a specific quadratic equation (see Eq. (24)), which determines the bifurcation long-range excitatory inputs 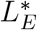. Therefore, *J*^*i*^ and 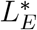 determine one of the bifurcation points in the two-dimensional space. The *i*^*th*^ brain area will be active only when it has long-range excitatory inputs such as 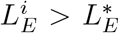. From another viewpoint, the bifurcation equation could be a quadratic equation of the scaled hierarchical value *J*^*i*^ (Eq. (25)). For a *i*^*th*^ brain area with long-range excitatory inputs 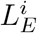, only when it has a hierarchical position *J*^*i*^ *> J*^∗^ displays non-zero firing rates. The critical line given by Eq. (23) is shown in Fig. 5e. We perform the same analysis for the Abbott-Chance transfer function. The steady-state equation for the large-scale model reads as follows:

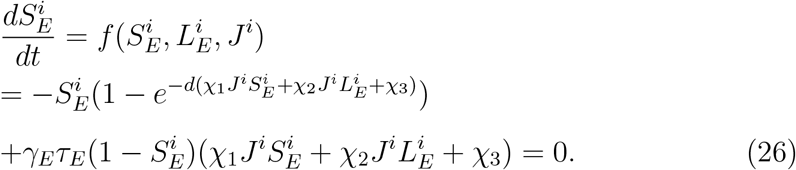

We use numerical methods to solve Eq. (26). Numerically solving the Eq. (26) will give a steady state surface shown in Fig. S10, Fig. S11, Fig. S12, and Fig. S13. We refer to this surface as *the r*(*h, L*_*E*_) *surface*. Any steady-state solution to the network’s dynamics will lay on this surface.

Remarkably, our network’s *r*(*h, L*_*E*_) surface has a very similar geometry to the cusp bifurcation normal form *r*(*h, L*_*E*_) surface (Kuznetsov et al. 1998). The cusp normal form is given by 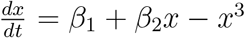, where *β*_1_ and *β*_2_ are two independent parameters (Kuznetsov et al. 1998; Strogatz 2016). The set of solutions to the steady state equation *β*_1_ + *β*_2_*x* − *x*^3^ = 0 gives the cusp normal form *r*(*h, L*_*E*_) surface in the (*β*_1_, *β*_2_) parameter space. We refer to this surface as the cusp surface. The cusp surface determines the possible bifurcations that the cusp normal form undergoes (Kuznetsov et al. 1998; Strogatz 2016), and with this, its bifurcation diagram. The cusp bifurcation point is given by *β*_1_ = *β*_2_ = 0. In Fig. S13, for illustration purposes, we overlay *β*_1_ and *β*_2_ axes to highlight the resemblance of our network’s *r*(*h, L*_*E*_) surface with the cusp surface (Kuznetsov et al. 1998).

Similarly to the cusp surface, in our network’s *r*(*h, L*_*E*_) surface, the bi-stable region is the region in the *J*^*i*^ and 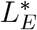 parameter space where, for a given active state, brain areas have two stable states: one with low and another with high firing rates. For the *r*(*h, L*_*E*_) surface, the bi-stable region increases with the increase of the transfer function gain *d*, and when *d* → ∞, the bi-stable region is the maximum. Thus, the *r*(*h, L*_*E*_) surface structure depends on the gain parameter *d*.

### Normal form analysis of a bifurcation in hierarchy space of the generative model

We derived a reduced equation for the dynamics of our large-scale neocortical network. We refer to this equation as the normal form of bifurcation in hierarchy space. Similar to classical normal forms in dynamical systems (Kuznetsov et al. 1998), this is a reduced dynamical equation derived from the network dynamical system, which qualitatively captures the network’s nonlinear dynamics close to the bifurcation in hierarchy space. We performed the derivation of this equation analytically for a network with a threshold-linear transfer function.

To calculate this equation, we first calculate the bifurcation points in the network dynamics. Based on the steady-state equation for the threshold-linear transfer function in Eq. (23), we have the bifurcation point 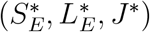 fulfill the following equation.

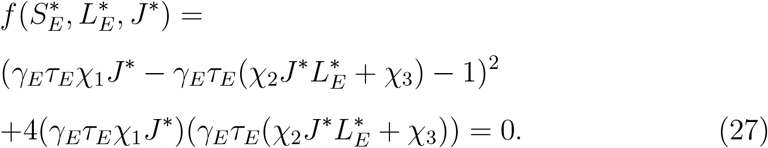

The bifurcation point in our multi-regional network with a threshold-linear transfer function is defined as the point in parameter space where the solutions of the quadratic equation in Eq. (23) change from complex conjugate to real. This point in parameter space corresponds to the appearance of bi-stability at the single-area level. Areas below the bifurcation point exhibit a single stable state with a low firing rate. Beyond this bifurcation point, cortical areas possess two stable states: one characterized by a low firing rate and the other by a high firing rate. Notably, during the emergence of the high firing rate state, each brain region undergoes an equivalent “saddle-node” bifurcation.

As we demonstrate in Eq. (19), if the final state of all brain regions corresponds to a steady state, then 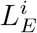 will be a fixed value-one that, however, depends on the connectivity structure of the *FLN*. For nonlinear dynamical systems characterized by complex interactions, it is typically necessary to compute center manifolds (Kuznetsov et al. 1998). These manifolds are not only dependent on the complex network structure (and thus vary across different cases) but also extremely difficult to express in a concise, elegant form.

Given the properties of 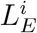 and *J*^*i*^, we therefore chose to treat these quantities as variable parameters and to consider each brain region in isolation during the derivation process. Consequently, to calculate the bifurcation within the hierarchical space normal form, it is essential to expand the function *f* around the bifurcation point 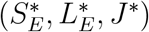-accounting for both the variables and the parameters in this expansion. We shift the variable and parameters to the bifurcation point and define 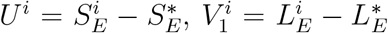, and 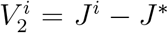. Additionally, we have 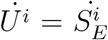.

The expanded function reads as follows:

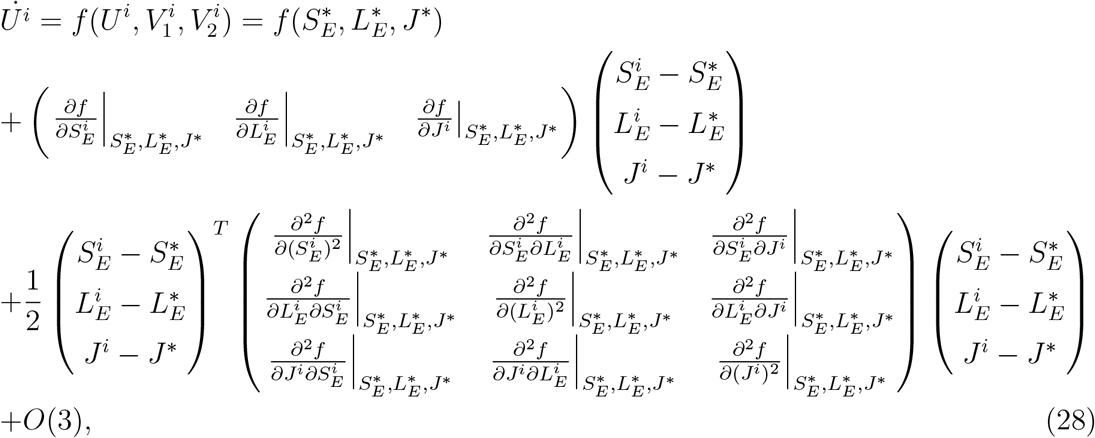

where we have:

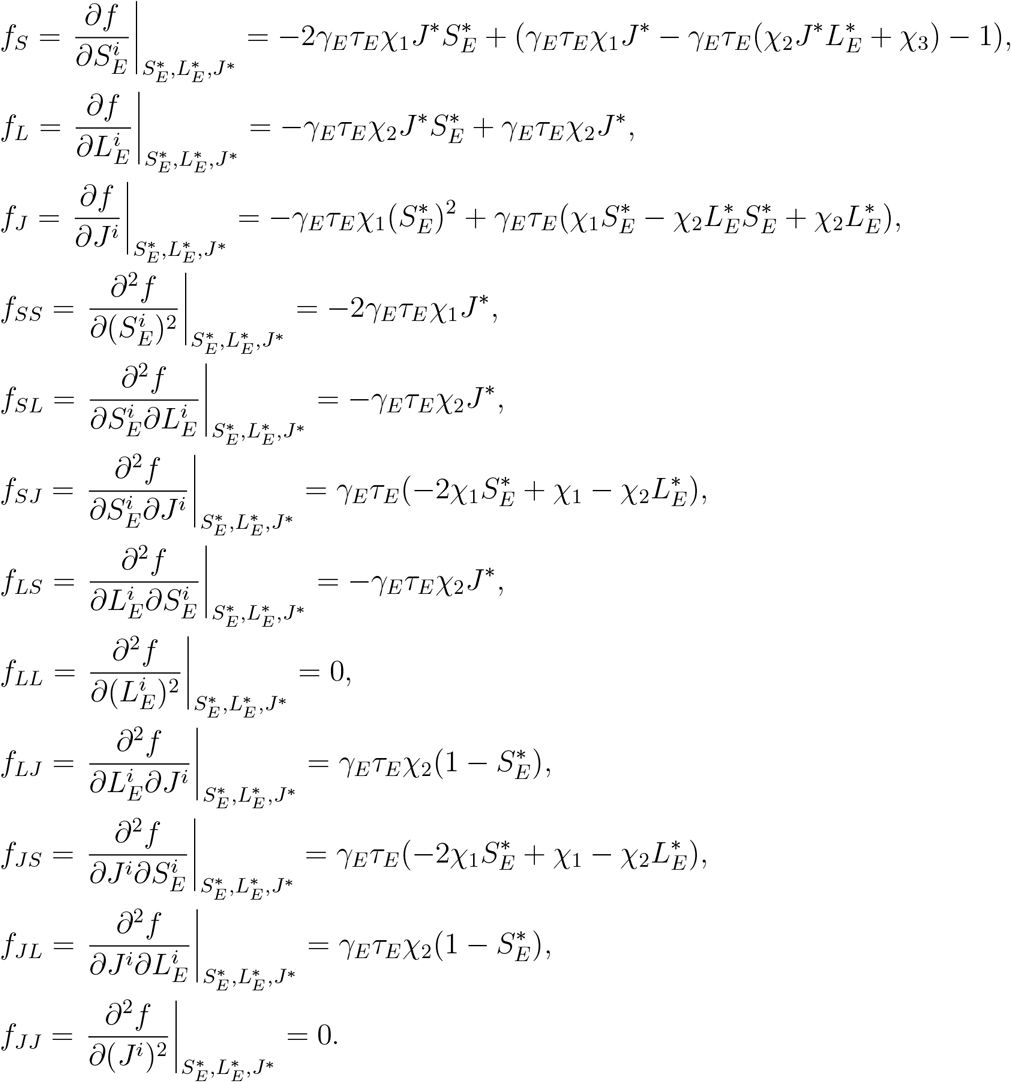

We have *f*_*SL*_ = *f LS, f*_*SJ*_ = *f JS, f*_*LJ*_ = *f JL*. After writing out each term in Eq. (28), we simplify the entire expression and obtain the simplified form as follows:

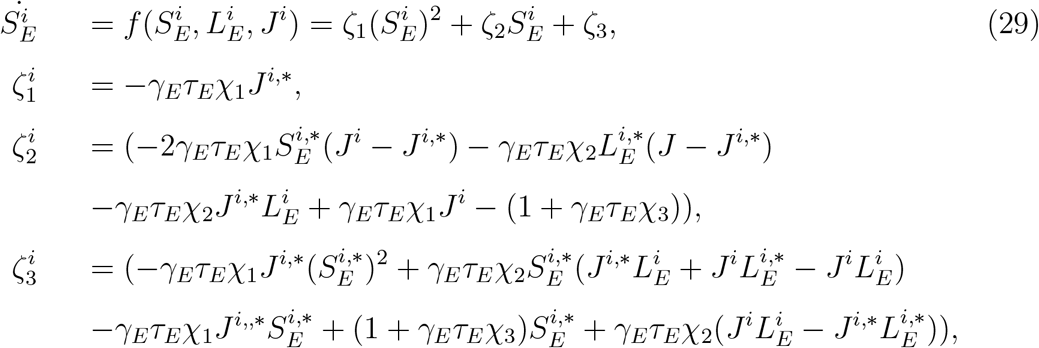

The aforementioned Eq. (29) is equivalent to the expanded normal form of bifurcation in the hierarchy space. Based on this ordinary differential equation, for each value of *J* = *J*^∗^, we can solve Eq. (25) to obtain the bifurcation long-range current 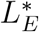, and then calculate the bifurcation 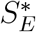. After determining the bifurcation point 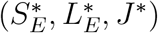, we can use the MATLAB function *fzero* to find the fixed point of 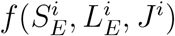. By following this process and iterating over all values of *J*, we can obtain all fixed points in the *J*-*L*_*E*_ space; these fixed points are naturally divided into two branches, as shown in Fig. 5e. The upper branch corresponds to the active states, while the lower branch corresponds to the unstable steady states.

We can also solve the steady state of the above Eq. (29) self-consistently and predict the firing rate of the delay period working memory states. The self-consistent equations read as

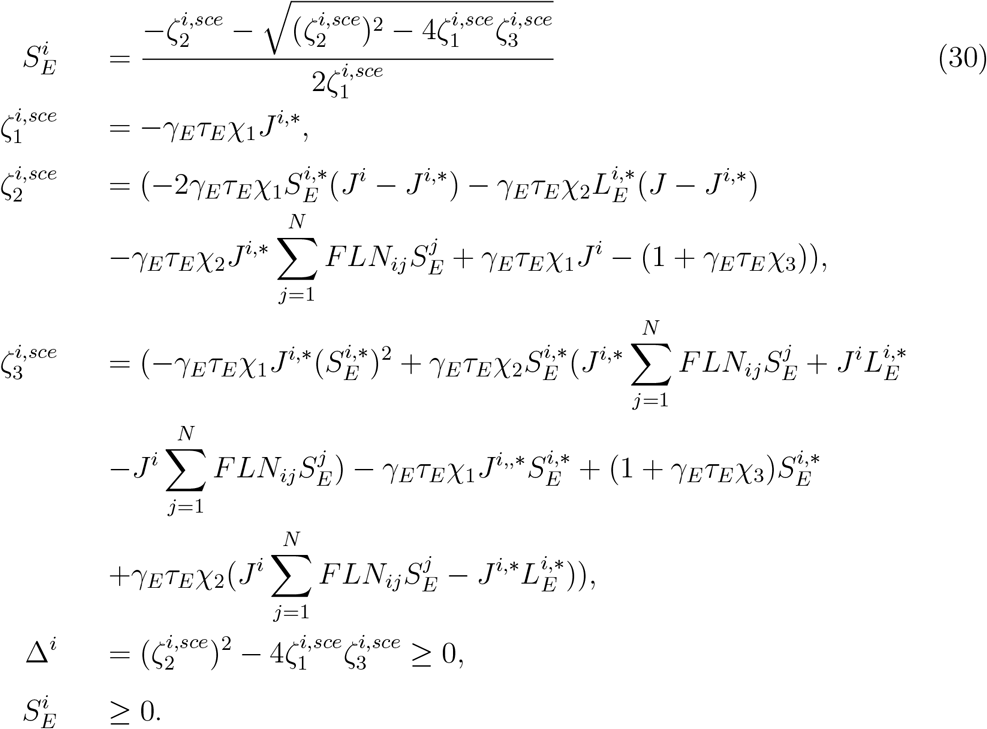

To solve the self-consistent equations, first, we solved Eq. (25) numerically and obtained *J*^*i*,∗^ and 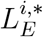 for the bifurcation point of the *i*^*th*^ brain area. Second, we determine the excitatory gating variable 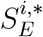 by inserting *J*^*i*,∗^ and 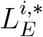 into Eq. (22) and solving numerically the Eq. (22). Third, we insert 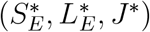 into the self-consistent equations in Eq. (30). Lastly, we solve the self-consistent equations in Eq. (30) and then predict the firing rate pattern of activity during an active state as shown in Fig. 4e.

The normal form in Eq. (29) can be further reduced to a more general expression, as follows:

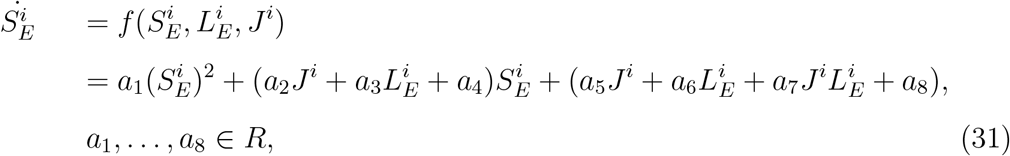

where *a*_1_, …, *a*_8_ are parameters calculated re-arranging terms in Eq. (29). The parameter values are *a*_1_ = −3.3, *a*_2_ = 0.08, *a*_3_ = −0.67, *a*_4_ = 3.11, *a*_5_ = 0.1, *a*_6_ = 0.3, *a*_7_ = 0.3127, *a*_8_ = −1.017 for a state like Fig. 5e. Remarkably, the bifurcation in the hierarchy space normal form has a similar mathematical form (up to a translation) as the saddle-node bifurcation (Kuznetsov et al. 1998). Actually, we could also transfer the expression of Eq. (28) into saddle-node canonical form. The Eq. (28) could be rewritten as follows:

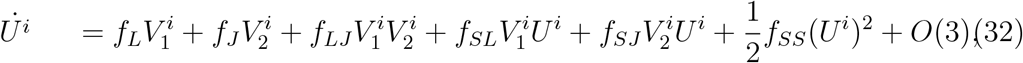

to eliminate the cross term 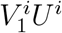 and 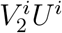, we define 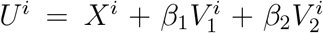. This definition yields the values *β*_1_ = −*f*_*SL*_*/f*_*SS*_ and *β*_2_ = −*f*_*SJ*_ */f*_*SS*_. After this transformation, the coefficient of the 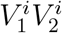 term changes to 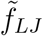, where 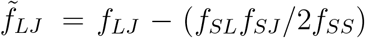. Additionally, the transformation causes the terms 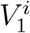 and 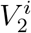 to appear with coefficients 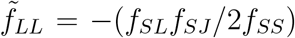 and 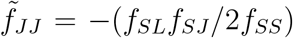, respectively. Finally, we obtain the canonical normal form of the bifurcation in the hierarchy space, which is given as follows:

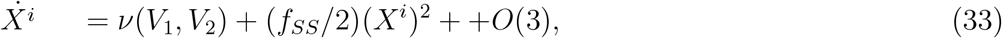

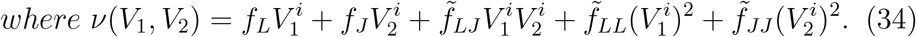

However, unlike the traditional saddle-node normal form, this model has cortical areas coupled via constant and linear coefficients. The coefficients *f*_*L*_ and *f*_*SL*_ depend on the hierarchy via *J*^*i*^, while *f*_*J*_ and *f*_*SJ*_ depend on the long-range excitatory input current 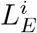. These coefficients denote both the heterogeneity of individual brain regions and the influence exerted by the network. Consequently, the normal form equation presented above demonstrates that bifurcation in space is determined by two key factors: the macroscopic gradients of neuronal properties and the structural characteristics of the neocortical network.

### Numerical methods for searching for spatial attractor states of generative models and connectome-based cortex models of monkey and mouse

In a network with between 1000 and 10000 brain areas, it is infeasible for our computational resources to find all the possible active states. Therefore, we try to determine as many unique active states as possible by using as many different initial conditions as possible. In practice, we ranked all 1000 cortical areas by their hierarchical level and then partitioned them into 20 groups according to this hierarchy. Accordingly, each group contained 50 cortical areas that were contiguous in the hierarchical order. To minimize variations in initial conditions, we assigned identical initial conditions to all brain areas within the same group.

We obtain the steady state by re-writing Eq. (19) as self-consistent equations for the variables 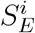 and 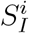 and iterating these equations to find the steady state solutions for the neural dynamics. The iterated equations read

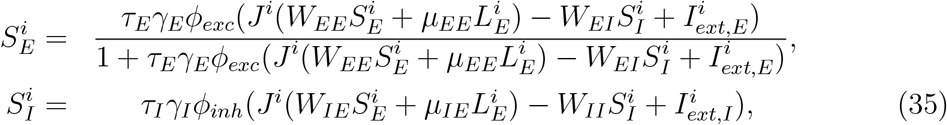

where the long-range excitatory inputs for the *i*^*th*^ brain area is given by 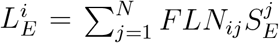. In any given initial condition, we use only two different initial values for all areas within a group: *S*_*E*_ = 1 or *S*_*E*_ = 0. Each group may take different values of *S*_*E*_. Using the above self-consistent equations, we search for the steady states from 2^20^ = 1, 048, 576 different initial conditions.

We iterate the Eq. (35) until the mean absolute difference between two consecutive iterations is smaller than 10^−10^, or the iteration number is larger than 10,000. However, in practice, no initial condition has more than 10,000 iterations for 1000 brain areas. After generating an active state, we determine whether it is unique by computing the absolute difference between it and all the unique active states we obtained previously. Once the sum of the absolute difference of all the brain areas is larger than 0.05, we retain it as a new unique state. Through this process, we obtained 4333 (one of the states in Fig. 6c is the resting state) distinct active states after trying 1,048,576 initial states. For all the distinct active states, we checked the local stability of the state by calculating all the eigenvalues of the Jacobian matrix. We found that the real part of all eigenvalues in all the active states is negative. Therefore, all the states are locally stable.

Since we lack a set of self-consistent equations for the monkey and mouse connectome models, we have adopted an alternative search strategy: specifically, we simulate the connectome model under different initial conditions. In practice, we first ranked all 41 cortical areas of the monkey and 43 cortical areas of the mouse based on their hierarchical positions. We then divided these areas into 21 groups (for the monkey) and 22 groups (for the mouse), respectively, following their hierarchical order. Consequently, the brain area with the highest hierarchical position in both the monkey and the mouse constitutes a group of its own. For each group, we tested two initial conditions: one high and one low. As a result, we ran the monkey connectome model with 2^21^ = 2, 097, 152 distinct initial conditions, and the mouse connectome model with 2^22^ = 4, 194, 304 distinct initial conditions. The connectome model simulations were performed using the Euler method, with a total simulation time of 35 seconds and a time step of 10^−4^ seconds. We retained only the final states where the difference in the gating variable between two consecutive steps was smaller than 10^−7^. After obtaining an active state, we verified its uniqueness by calculating the absolute difference between this state and all previously identified unique active states. If the sum of the absolute differences across all brain areas exceeded 0.01, the state was retained as a new unique active state. Through this process, we identified 1275 distinct active states for the monkey connectome model (Fig. 6a) and 54 distinct active states for the mouse connectome model (Fig. 6b).

## Supplementary figures

**Figure S1:**
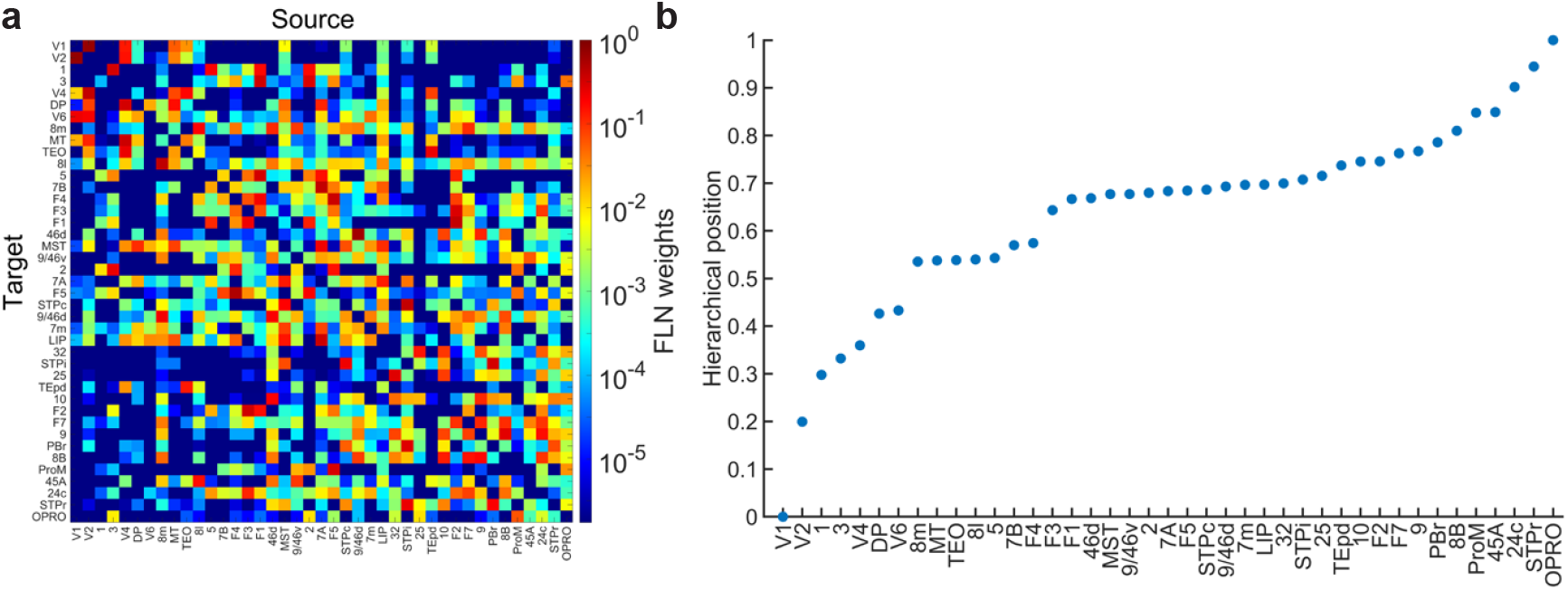
FLN Matrices and Hierarchical Positions of Brain areas in Macaque Connectome Data. (a) The FLN matrix of macaque connectome with 41 brain areas through retrograde tracing (Markov et al. 2014). (b) The hierarchical position is estimated based on the 41 brain areas’ percentage of supragranular labeled neurons (SLN); the actual method can be found in (Markov et al. 2014).

**Figure S2:**
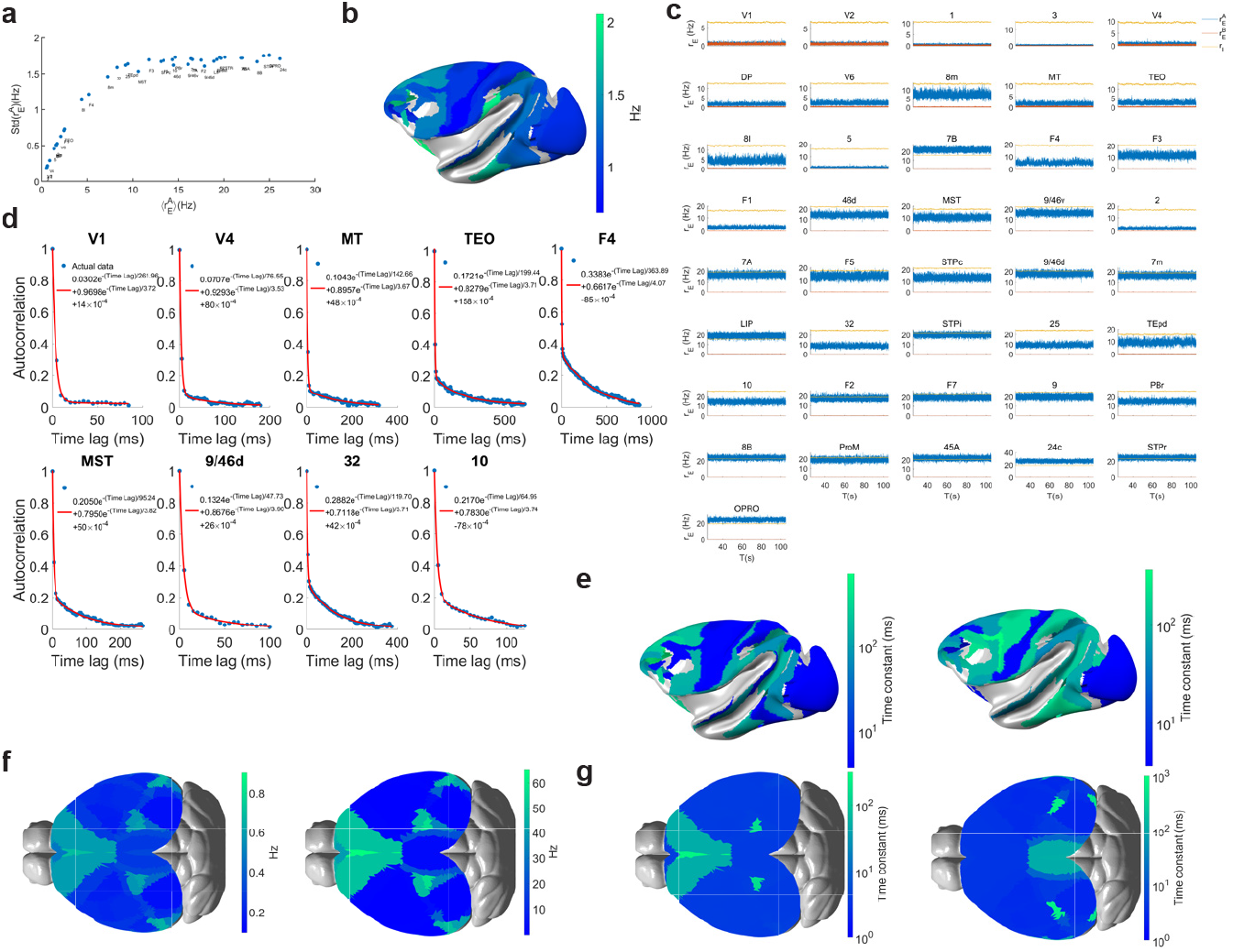
This is the supplementary figure illustrating bifurcation in the hierarchical space of connectome-based cortical models of macaque monkey and mouse. (a) The average firing rate versus the standard deviation of each brain area of the macaque neocortex with 41 brain areas. (b) Spatial activity map of the resting state of the macaque neocortex model with 41 brain areas with the model in (Mejias and Wang 2022). (c) The firing rate of excitatory population A (blue), B (brown), and inhibitory population (yellow) of the 41 brain areas. (d) The auto-correlation function and the related double exponential fitting function of the nine chosen brain areas are in Fig. 2d in the main text. (e) The spatial time constant map of 41 brain areas for resting state (left) and delay period working memory state (right) corresponds to the states of panel b and Fig. 8a right panel. (f) Spatial activity map of resting state (left) and the delay period working memory state (right) of the large-scale mouse brain model with 43 brain areas with the model in (Ding et al. 2024). (g) The spatial time constant map of 43 brain areas for resting state (left) and delay period working memory state (right) corresponds to the states of panel f.

**Figure S3:**
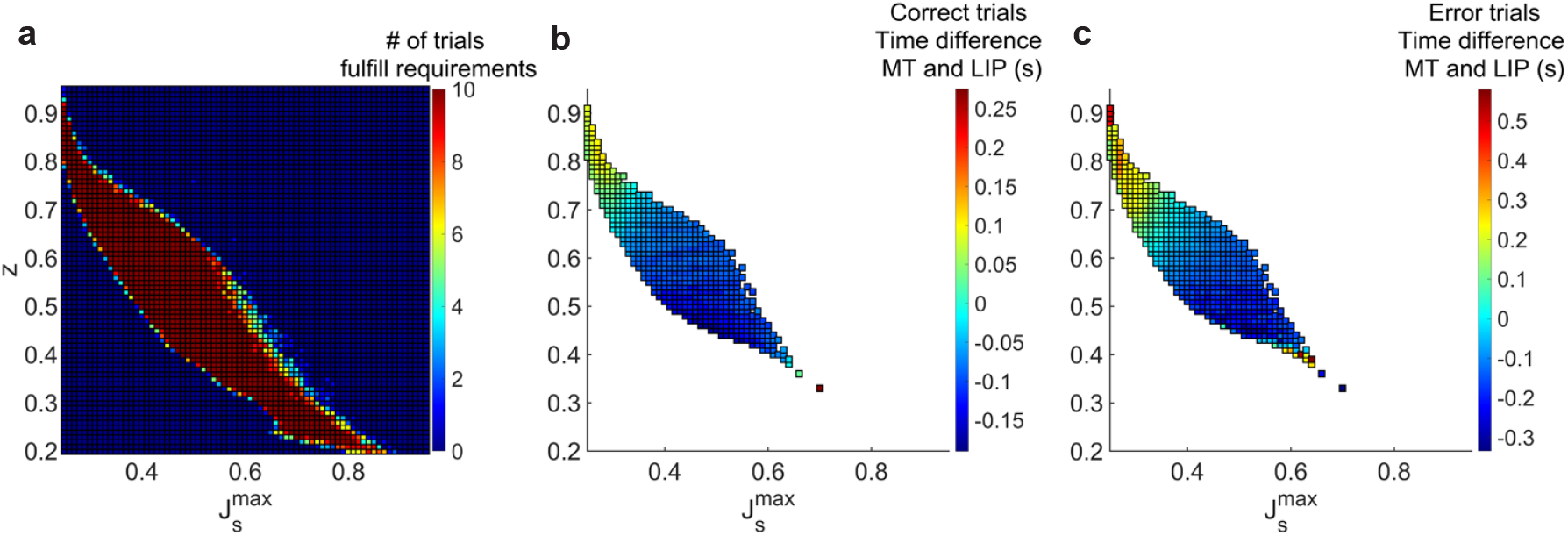
The parameter region in 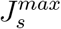 and *Z* space that fulfills all the requirements of working memory and decision-making of connectome-based cortical models of the macaque monkey with 41 brain areas. We updated Mejias’ model (Mejias and Wang 2022) with 41 brain areas and their corresponding hierarchical values. After keeping all the other parameters consistent with the original model (Mejias and Wang 2022), we searched the parameter sets of 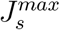 and *Z* that meet the following requirements: 1. the resting state must be stable across all brain areas; 2. persistent activity emerges in some brain areas to encode working memory; 3. the brain region MST can exhibit persistent activity, while the brain region MT cannot, which is consistent with experimental findings (Mendoza-Halliday et al. 2014). (a) shows the number of trials that fulfill all the requirements of the parameter set of 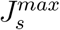 and *Z*. We did 10 trials for each parameter set. If one parameter set with all those trials fulfills all the requirements, it has a value of 10. If no trial fulfills all the requirements, it has a value of 0. (b) shows the mean difference in time to threshold =0.55 between *MT* and *LIP* for correct trials during decision-making across parameter sets in A. We simulated 5000 trials for each pair of 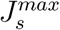 and *Z*, identifying parameters showing both correct and some error trials. The distribution highlights parameter sets with the highest positive difference between *MT* and *LIP* during correct trials. (c) shows the mean difference in time to threshold =0.55 between *MT* and *LIP* for error trials using the same set of parameters as panel b. The parameters 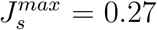 and *Z* = 0.82 were chosen for further analysis based on their optimal performance in correct trials and maintaining 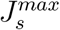 below the critical bistability value of 0.4655. For these values, 200 correct and error trials were generated for additional analyses.

**Figure S4:**
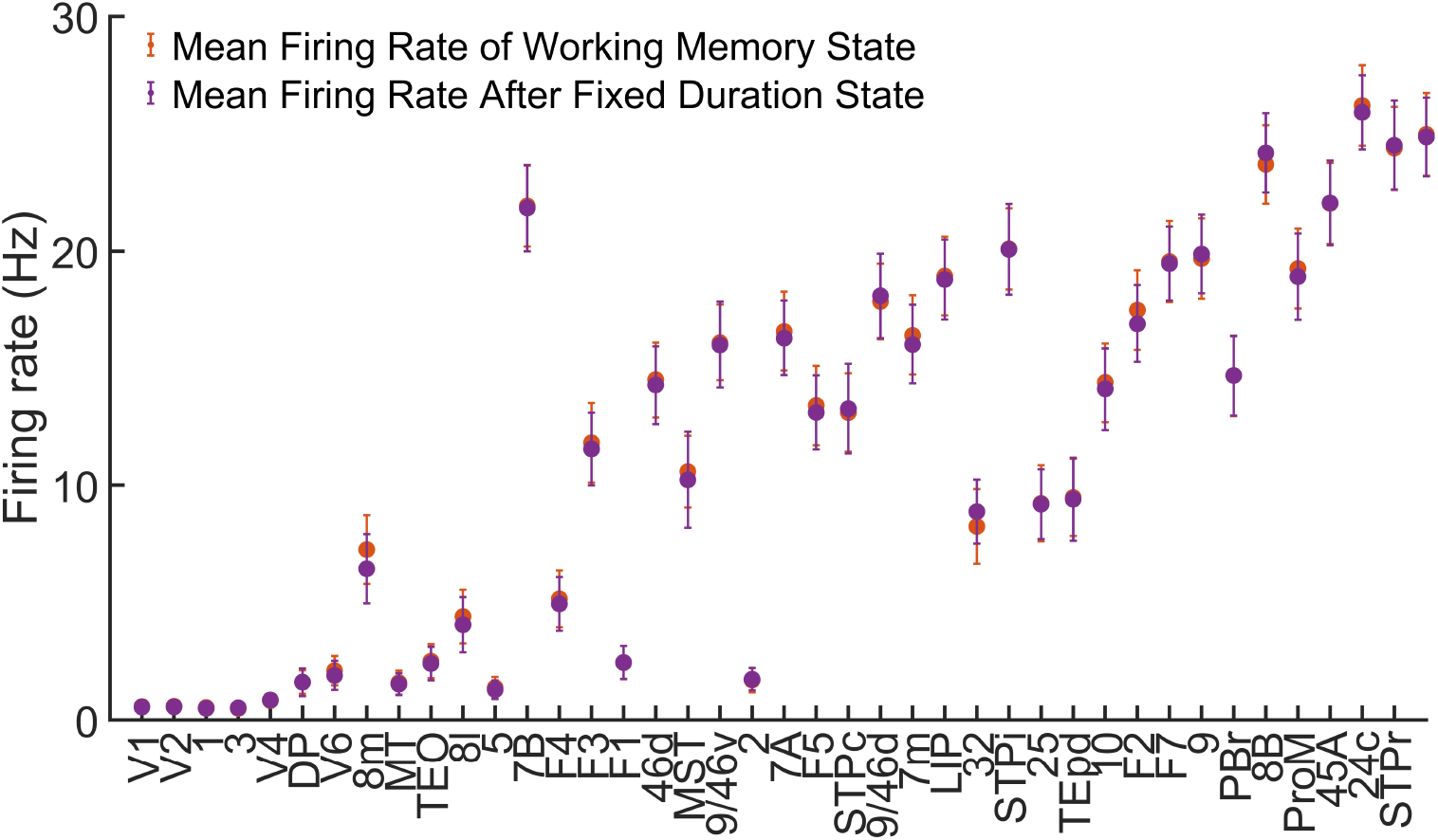
The delay-period activity for the fixed-duration decision-making task and the working memory state is depicted. The dots indicate the average firing rate, while the error bars represent the standard deviation of the firing rate.

**Figure S5:**
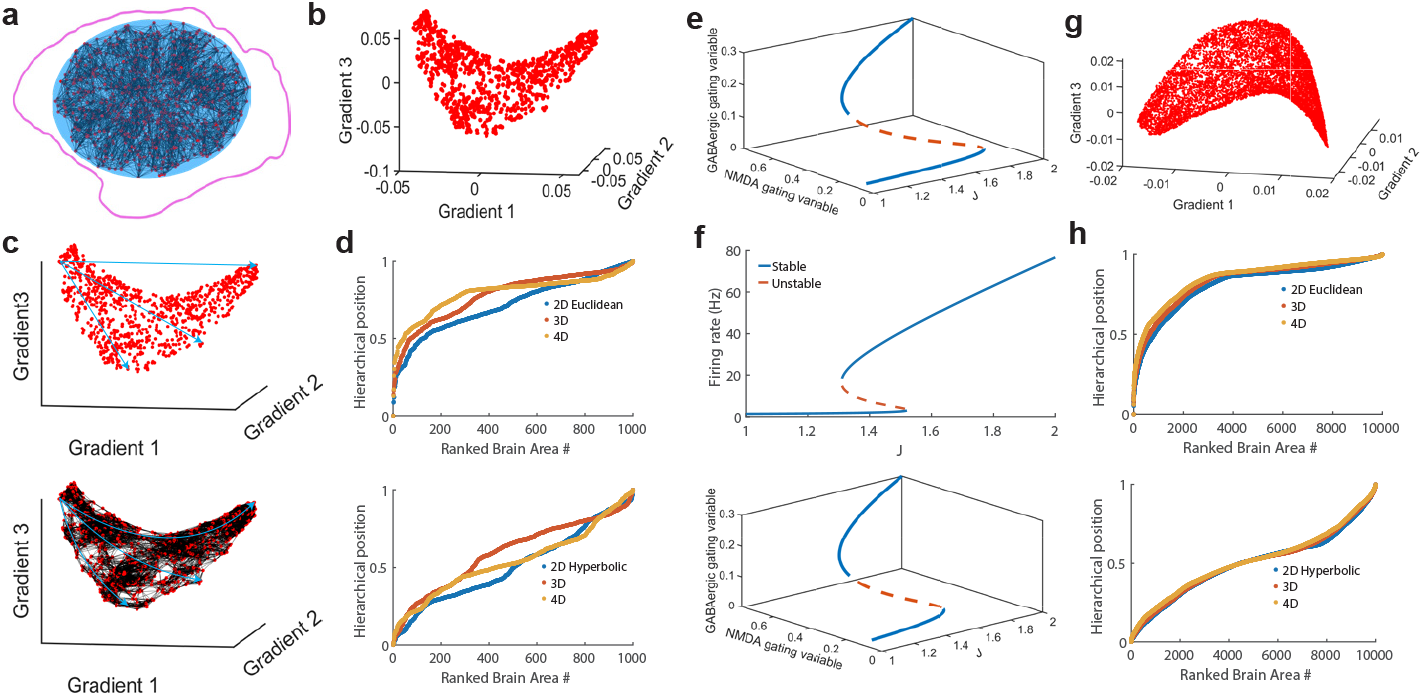
Supplementary figure showing connectivity, hierarchy and local circuit models generated in the study. (a) The generated super neocortex network with 1000 brain areas is consistent with the statistics of the macaque neocortex network. (b) The three-dimensional embedding of the super neocortex network (Panel a) shows hyperbolic characteristics. (c) Illustrates Euclidean (top) and hyperbolic (bottom) distances in the 3D embedding of the super neocortex network. (d) The Euclidean (top) and hyperbolic (bottom) hierarchical positions of all brain areas. The blue, brown and yellow lines represent the 2D, 3D and 4D embeddings of the constructed super neocortex model. (e) The NMDA and GABAergic gating variables’ steady states vary with the hierarchical position of an isolated brain area with simplified dynamics. This isolated brain area also has the same parameter settings as Fig. 4g in the main text. (f) The upper panel shows the bifurcation diagram of an isolated brain area with simplified dynamics, where the gain parameter is set to *d* = 0.157. The lower panel illustrates how the steady states of NMDA and GABAergic gating variables in this brain area change with its hierarchical position. (g) The generated super neocortex with 10,000 brain areas presents a hyperbolic shape in its 3D embedding. (h) Euclidean (upper) and hyperbolic (lower) hierarchical positions of all regions in the 10,000-area brain network. Blue, brown and yellow lines represent the 2D, 3D and 4D embeddings of the synthetic 10,000 areas super neocortex model.

**Figure S6:**
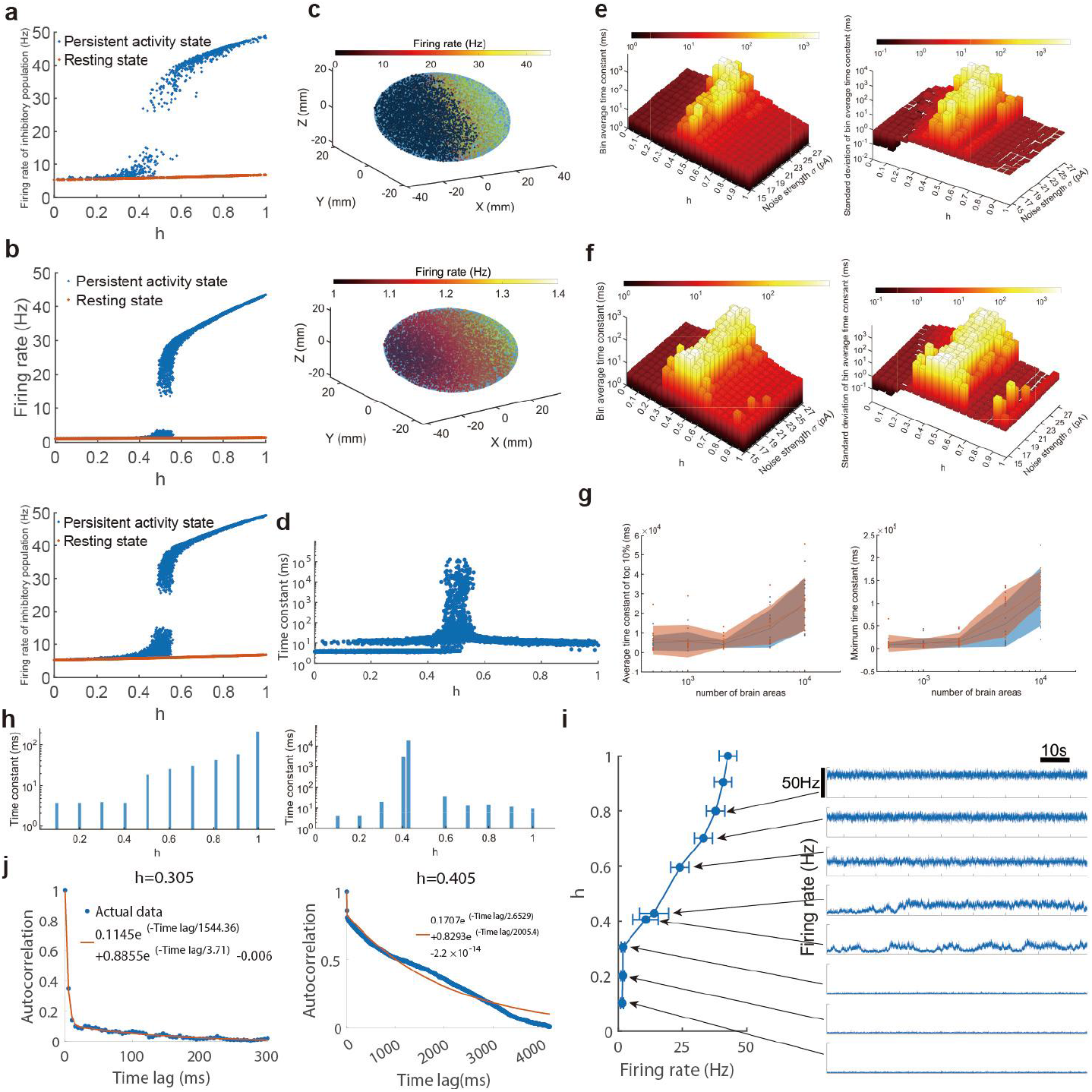
Supplementary Figure of bifurcation in hierarchical space. (a) The firing rates of all inhibitory populations under the same parameter set as panel a in Fig. 4g in the main text. (b) The firing rates of excitatory (upper) and inhibitory (lower) populations in active (blue) and resting (brown) states across all 10,000 brain areas in the generated super-neocortex network. (c) The spatial distribution of persistent firing rates in monotonic active states (top) and resting-state firing rates (lower) across an ellipsoidal space containing 10,000 brain areas. (d) The time constant of all the brain areas at the monotonic active state of panel b with 10,000 brain areas and noise strength *σ* = 24*pA*. (e) The distribution of bin average (left) and standard deviation of bin-averaged time constant (right) along the hierarchical position changes with the noise strength, which increases from 15*pA* to 27*pA*, of 5 noise ensemble. In this panel, we adopt the bin-averaged time constants from Panel a, where the bin size corresponds to a hierarchical position interval of 0.05. For each bin of time constants, the values are averaged across 5 ensemble realizations. (f) Across hierarchical positions, the distributions of binned averages (left) and standard deviations of bin-averaged time constants (right) vary with noise strength (15pA to 27pA). The bin size matches that in Panel e. All averages were computed over 5 noise ensemble realizations and 10 distinct super-neocortex networks. (g) The average of the 10% largest time constants (left) and the maximum time constant (right) vary with network size. The ten dots corresponding to a given number of brain areas represent ten distinct super neocortex networks. The shaded area indicates the range within one standard deviation. (h) The time constants of 10 cortical areas in the persistent activity state (right) and 10 areas in the resting state (left). The states correspond to the states in panel g of Fig. 4. (i) The mean and standard deviation of firing rates and their corresponding time series for ten selected cortical regions during persistent activity. (j) The autocorrelation and double exponential fitting were performed for two selected brain regions with h=0.305 and h=0.405, respectively. These two brain areas correspond to the bottom and top points of the inverted V-shaped profile of the time constant during active states.

**Figure S7:**
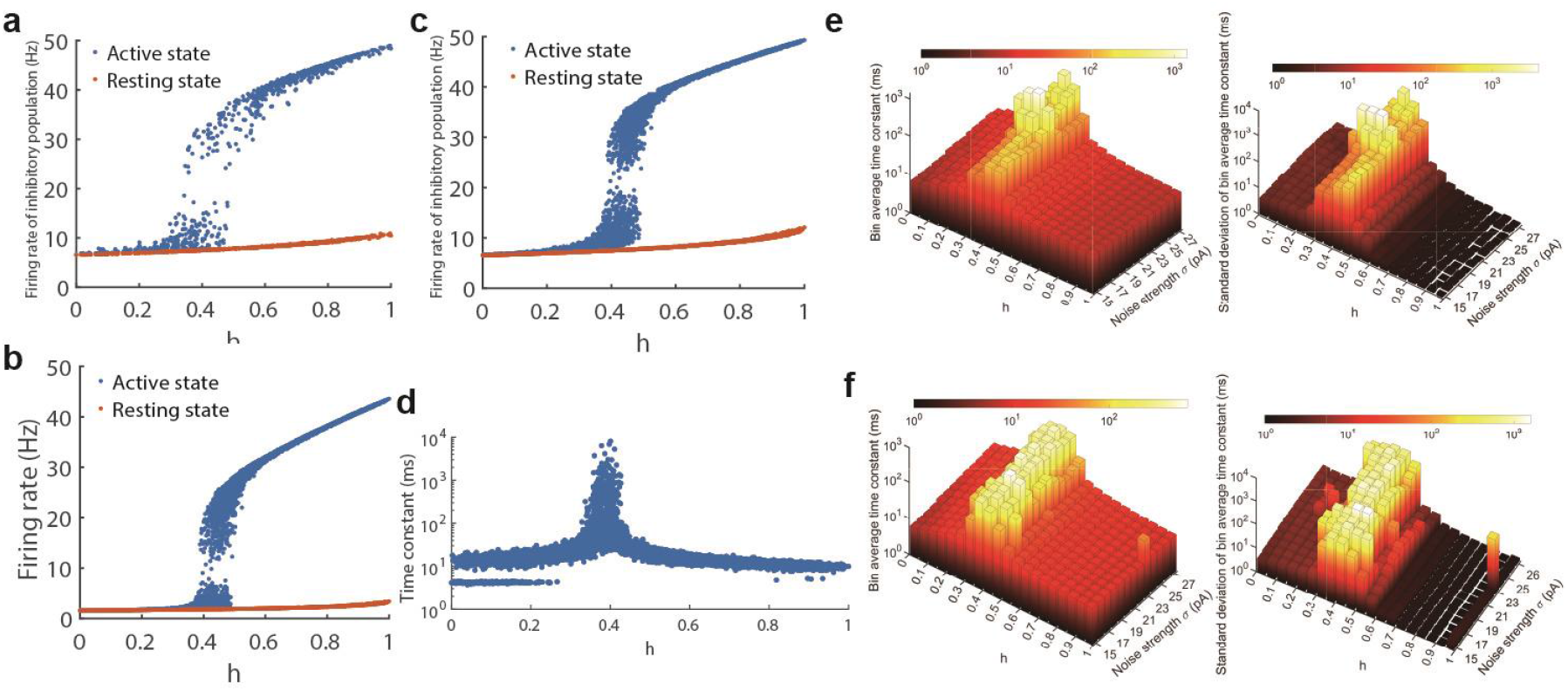
The delay activity state and time constant of brain networks with 10,000 brain areas (d=0.157) (a) The firing rates of all inhibitory populations, corresponding to panel b in Fig. 5 of the main text, with d=0.157. (b) The firing rates of excitatory populations in both active (blue) and resting (brown) states are shown for all brain areas in the synthetic super neocortex network. This network contains 10,000 brain areas and uses the same parameter settings as Panel a. (c) The firing rates of inhibitory populations in all brain areas during active and resting states are consistent with the settings in panel b. (d) The time constants of all 10,000 brain regions are measured under the monotonic active state in Panel b, with a noise strength of *σ* = 24*pA*. (e) The distributions of bin averages (left) and standard deviations of bin-averaged time constants (right) vary with noise strength (ranging from 15 pA to 27 pA) across five noise ensembles. In this panel, we adopt the bin-averaged time constants from Panel e of Fig. S6, with a bin size corresponding to a hierarchical position interval of 0.05. For each bin of time constants, we take the average across 5 ensemble realizations. All other parameters remain identical to those in Panel d. (f) The distributions of bin averages (left) and standard deviations of bin-averaged time constants (right) across hierarchical positions change with noise strength (ranging from 15pA to 27pA). Results are obtained from 5 noise ensembles and 10 network ensembles. The bin size is consistent with that in panel e. Averages were calculated across 10 distinct super-neocortex networks. For each bin of time constants, we performed averaging over 5 noise ensemble realizations and 10 different super-neocortex networks.

**Figure S8:**
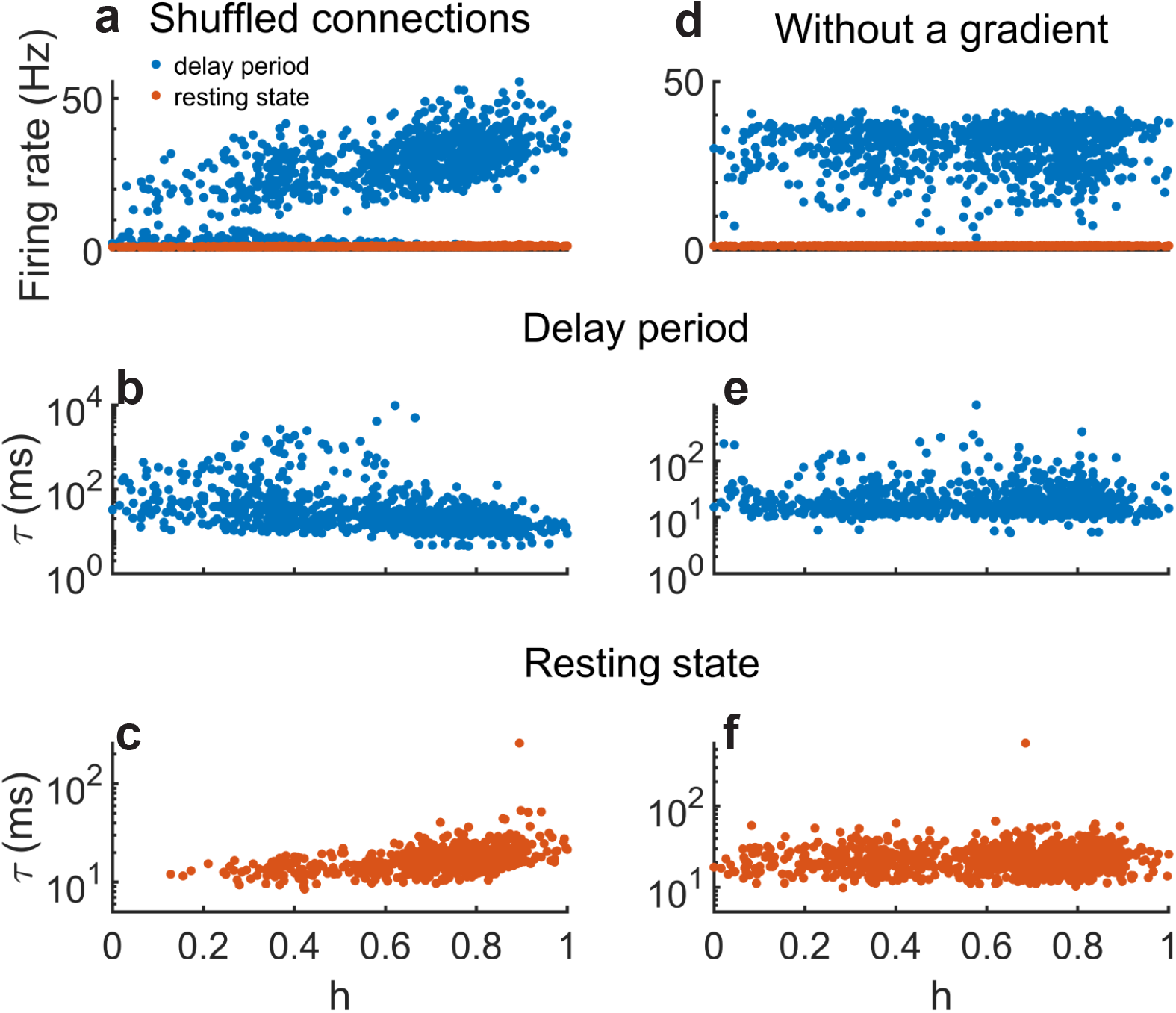
The firing rates and time constants of distributed attractor states when the network connection weights are randomly shuffled (a-c), or the macroscopic gradient of excitation is abolished by randomly shuffling the parameter J (d-f). In either case, the model still displays the bistability of a resting state (orange) and an elevated persistent activity state (blue). However, modularity disappears because all the areas participate in the persistent activity state, and there is no inverted-V shaped profile of time constants in the active state.

**Figure S9:**
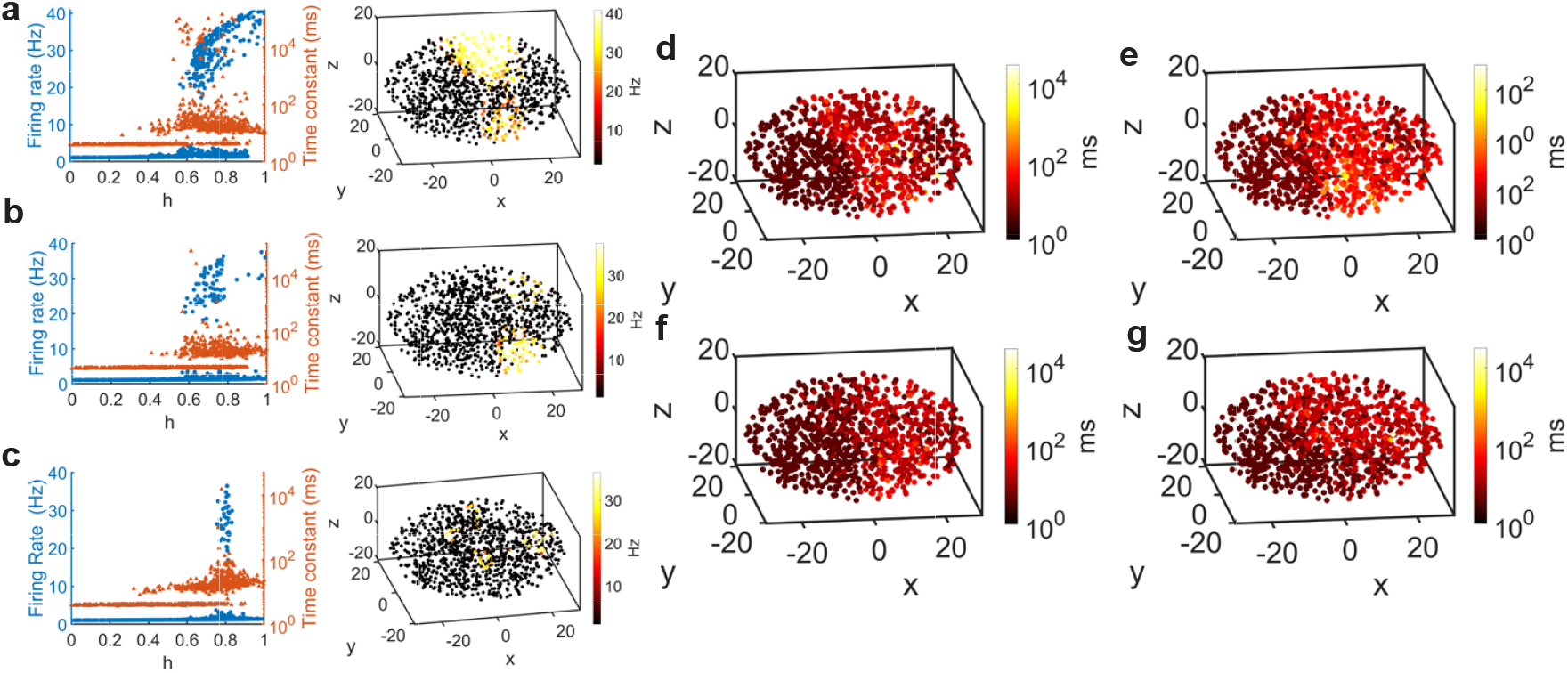
The supplementary figure of the diversity of distributed working memory states. (a-c) The firing rate (blue) and time constant (brown) of the active state correspond respectively to the 2300th, 3454th and 4068th states among the 4333 active states shown in Fig. 6c. The spatial distributions of the firing rates of the 2300th, 3454th and 4068th active states (right) in the generative model ellipsoid are shown respectively. In the generative model ellipsoid, the spatial distributions correspond to the 2300th, 2489th, 3454th and 4068th time constants of active states respectively. The firing rate and time constant of the 2489th active state are shown in Fig. 6d.

**Figure S10:**
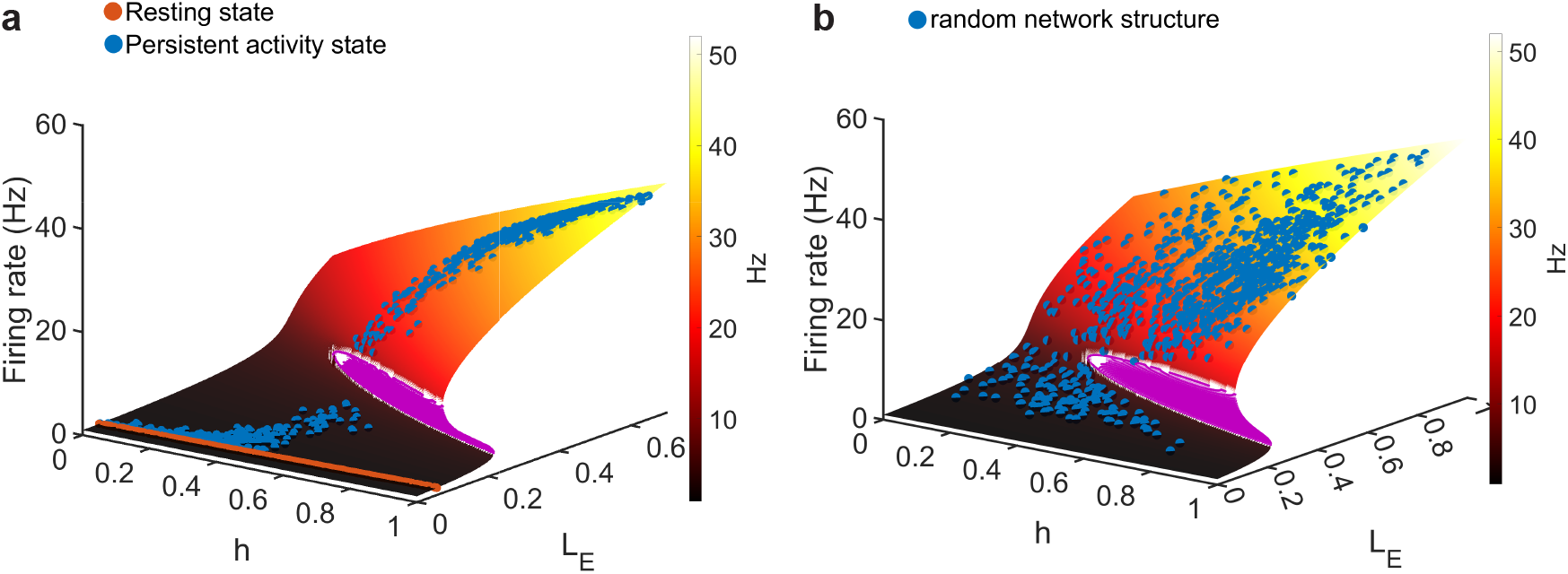
The geometry of distributed attractor states in parameter space. (a) The neocortex model’s resting (brown) and persistent activity state (blue) lie on top of the *r*(*h, L*_*E*_) surface with *d* = 0.17 (corresponding to Fig. 4g). (b) Mapping of active delay states of randomly shuffled generative network connections onto the *r*(*h, L*_*E*_) surface (corresponding to Fig. S8a).

**Figure S11:**
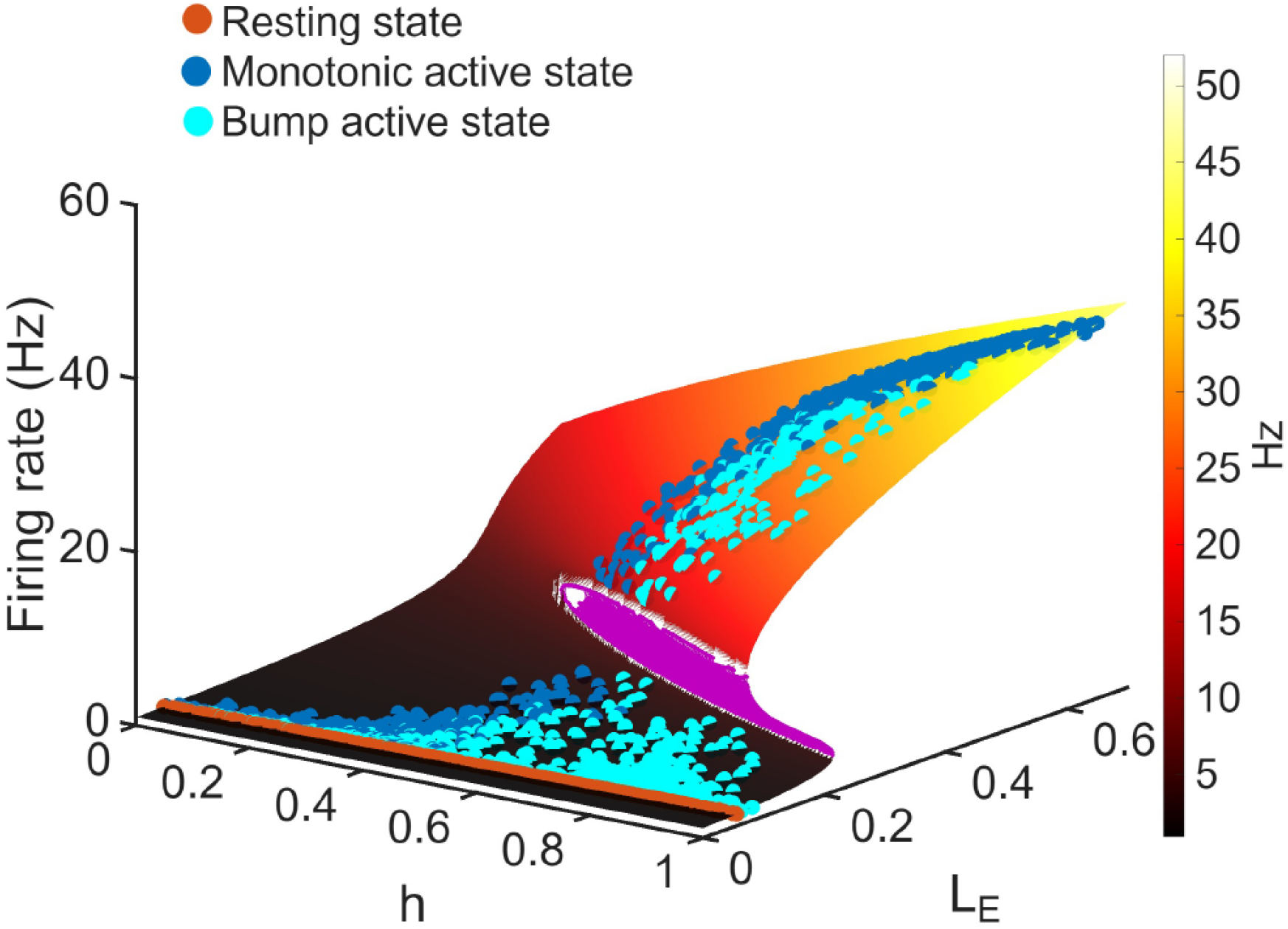
The *r*(*h, L*_*E*_) surface of *d* = 0.17. Firing rates of all the areas in the resting state, a monotonic state, and a localized bump state lie on the *r*(*h, L*_*E*_) surface.

**Figure S12:**
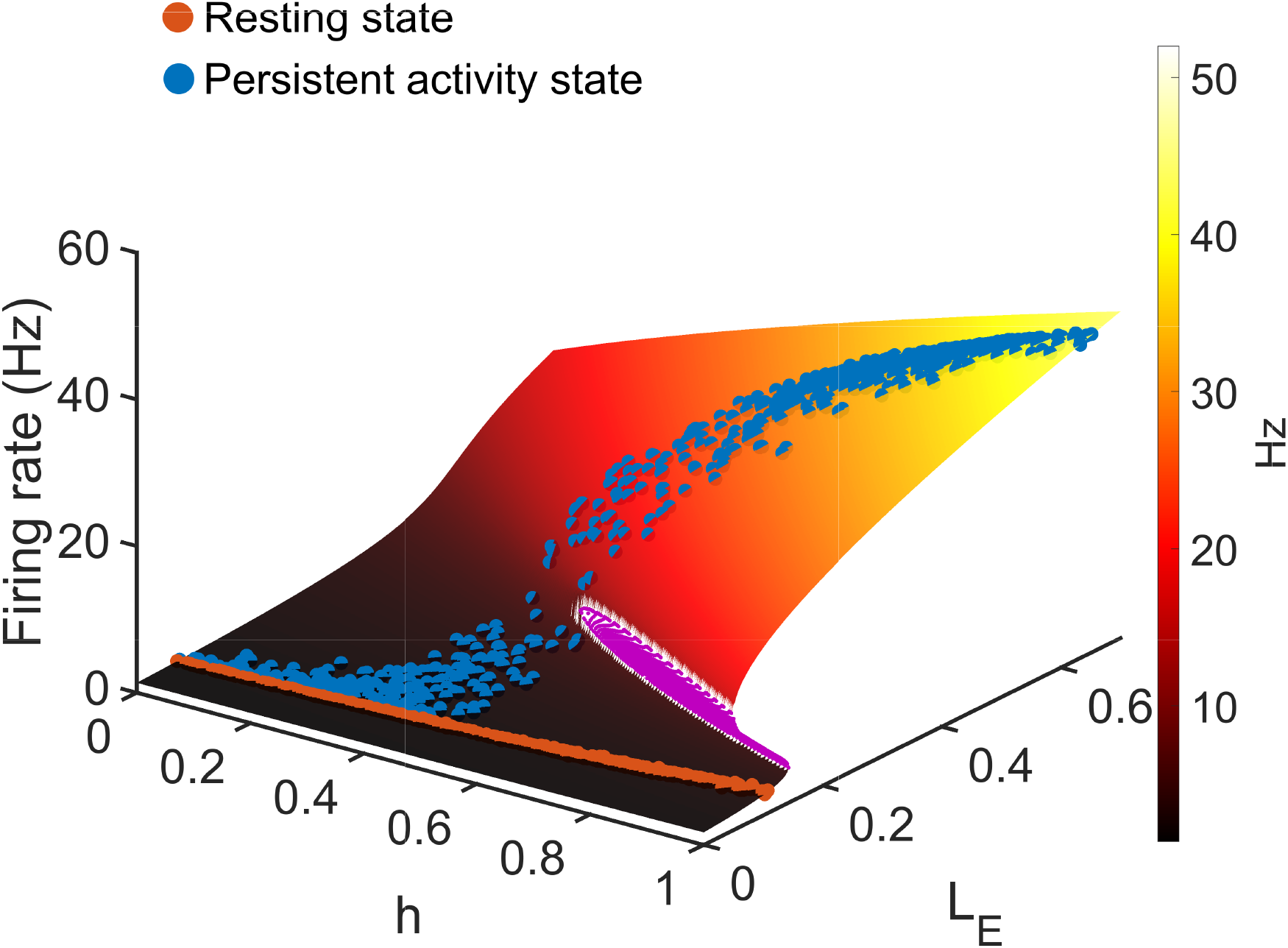
The *r*(*h, L*_*E*_) surface of *d* = 0.157. The neocortex model’s resting state (brown) and persistent activity state (blue) lie on top of the *r*(*h, L*_*E*_) surface (*d* = 0.157). Here, the active state transitions continuously with no gap in firing rate.

**Figure S13:**
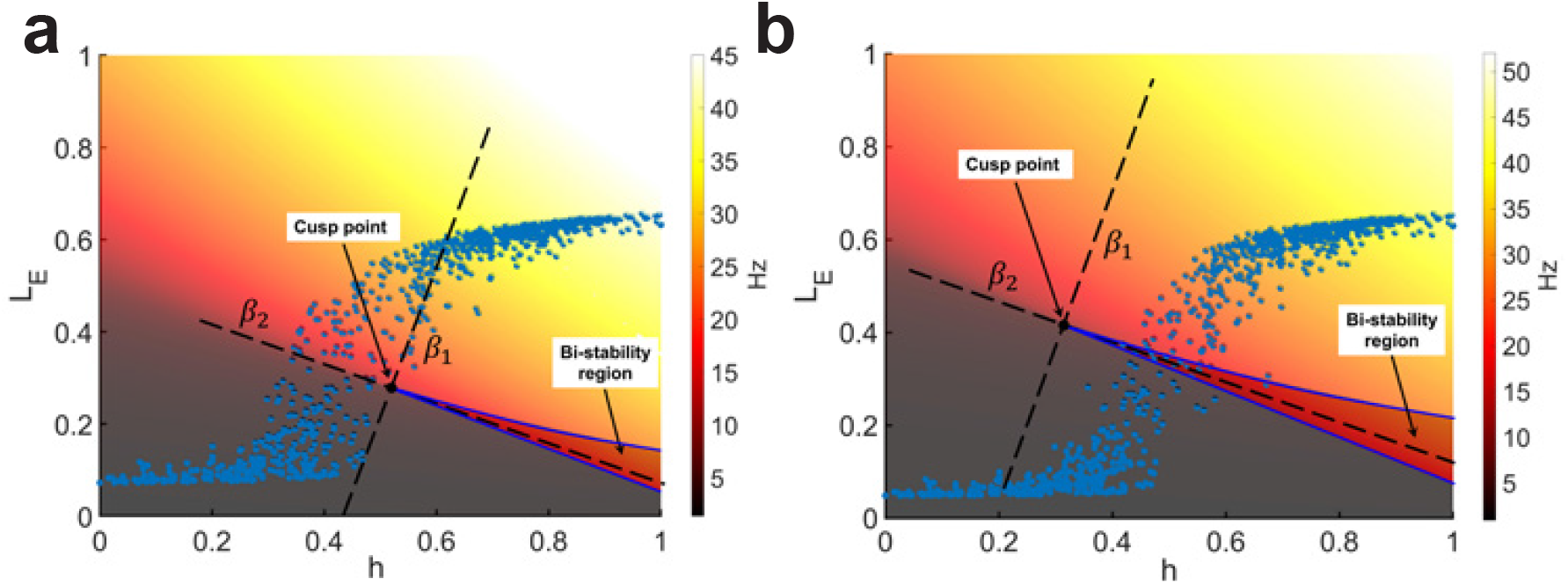
Cusp geometry determines the bifurcation in hierarchical space. (a and b) The geometry of the *r*(*h, L*_*E*_) surface and the cusp point determine the bifurcation in hierarchical space. The blue dots correspond to the persistent activity states of the neocortex model with *d* = 0.157 (panel a) and *d* = 0.17 (panel b), respectively. For comparison purposes, the axes *β*_1_ and *β*_2_, which correspond to the two control parameters in the cusp bifurcation normal form (see Methods), are overlaid on the *r*(*h, L*_*E*_) surface.From the figure, we can see that for regions at lower hierarchical levels, the firing rate increases smoothly with *h* and *L*_*E*_. After this cusp point, the surface of *r*(*h, L*_*E*_) becomes folded when *h* (hierarchy level) ranges from 0.6 to 1 and *L*_*E*_ (weighted long-range excitatory input current) ranges from 0.1 to 0.2, respectively.

**Figure S14:**
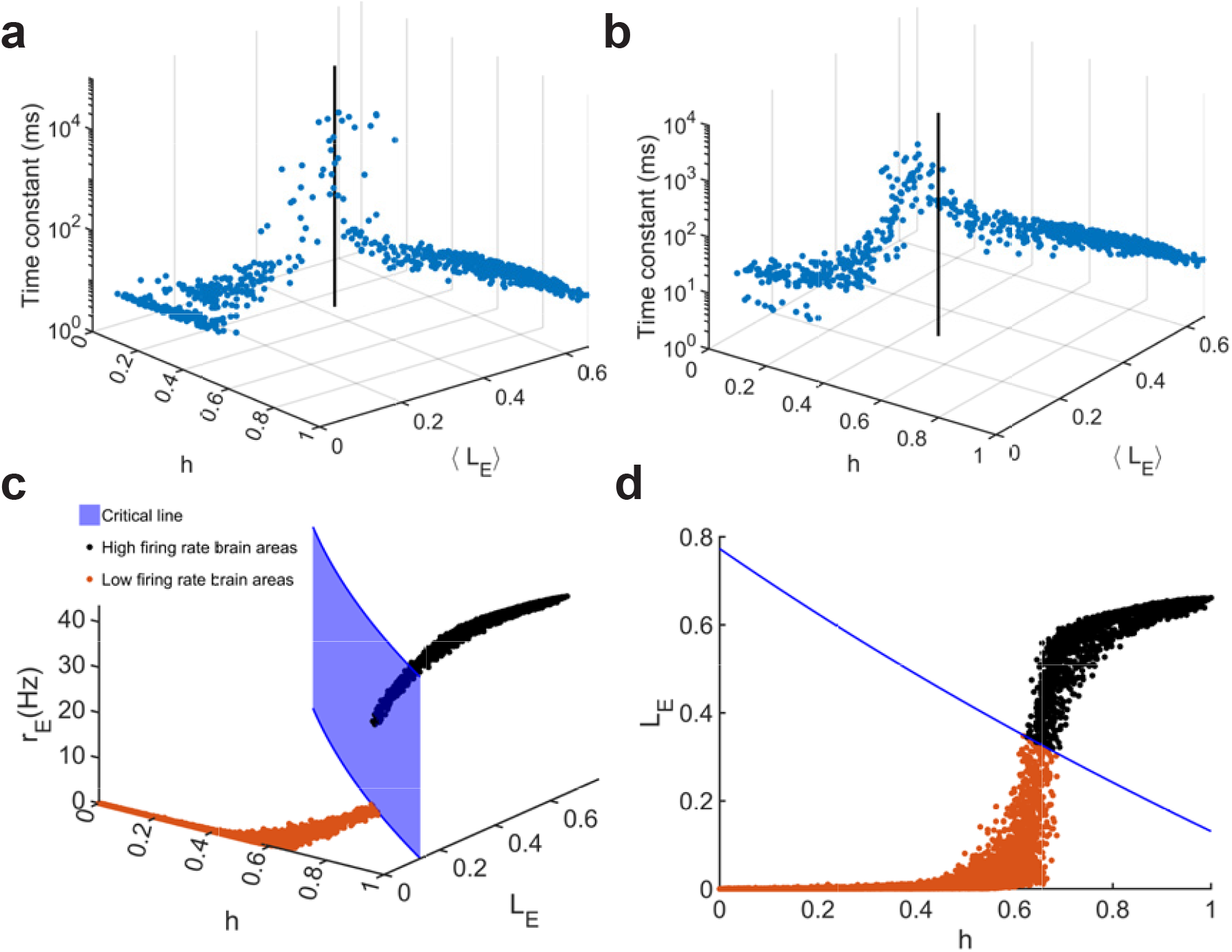
Time Constant Near the Cusp Point and Geometry of the *r*(*h, L*_*E*_) Surface for a Network of 10,000 Brain Areas with Threshold-Linear Transfer Function. (a-b) The time constants of the 1000 brain regions are plotted in the *h* - ⟨*L*_*E*_⟩ space for d=0.17 and d=0.157, respectively. ⟨*L*_*E*_⟩ denotes the average long-range gating variable calculated over a duration of 80 seconds. The noise strengths for panels a and b match those for panel i in Fig. 4 and panel b in Fig. 5, respectively. The black line indicates the *h* and *L*_*E*_ values of the estimated “cusp point.” (c) The critical line and firing rate distribution in *h* and *L*_*E*_ space for the threshold-linear transfer function with 10000 brain areas. (d) the top view of panel c.

**Figure S15:**
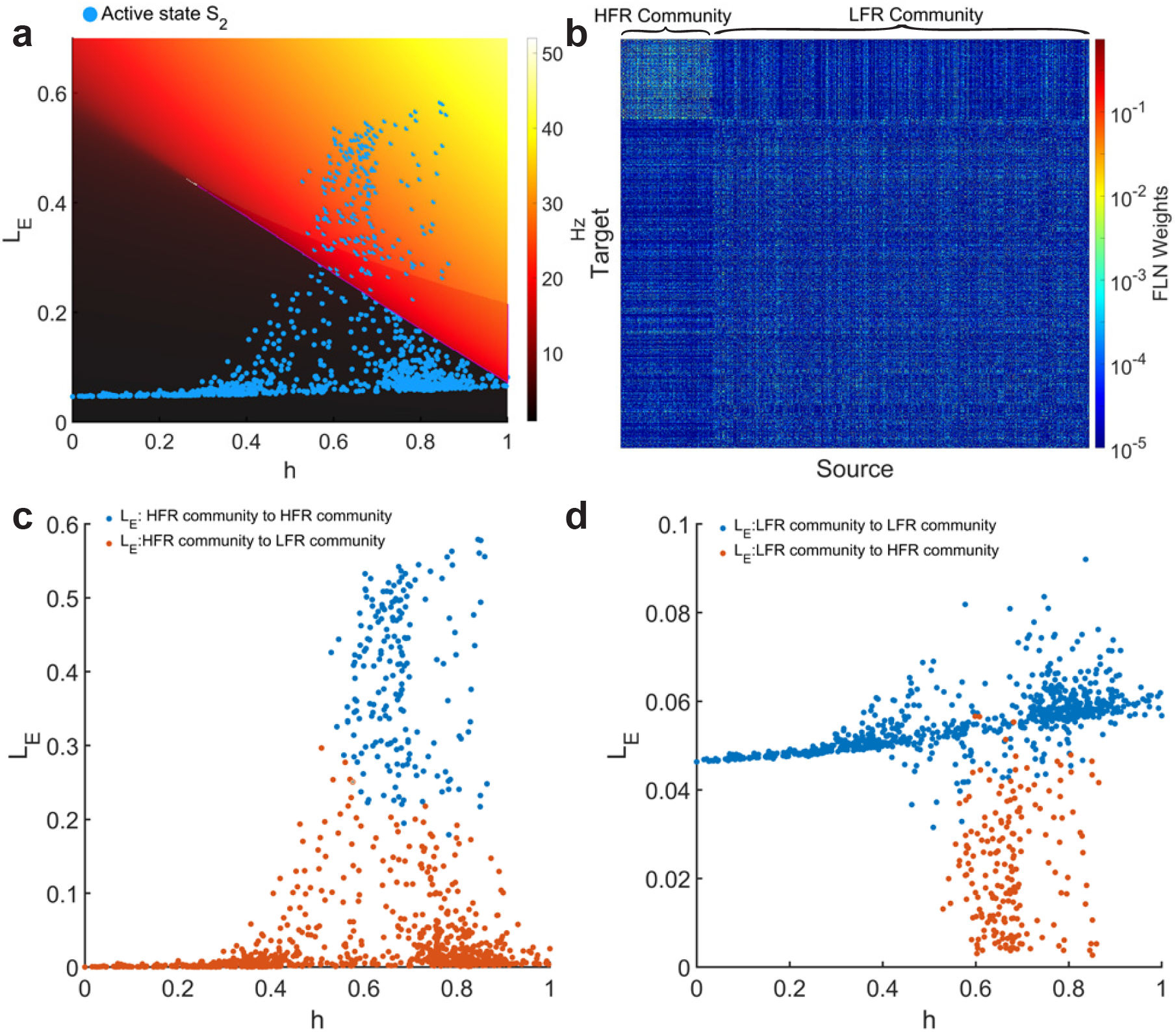
The supplementary figure of *L*_*E*_ of the 2489th localized bump active state (Fig. 6d). In the localized bump state, the weighted long-range excitatory input (*L*_*E*_) is widely distributed within the hierarchical localized bump region. Brain areas receiving a sufficiently large *L*_*E*_ will exhibit a high firing rate, whereas those with a small *L*_*E*_ will only achieve a low firing rate state. The high firing rate (HFR) community also has a greater average connection strength than the low firing rate (LFR) community. Due to the community structure of the connections, the long-range excitatory input within the HFR community exceeds the input from the HFR community to the LFR community. Conversely, the long-range excitatory input within the LFR community is greater than the input from the LFR community to the HFR community. This pattern is a result of the normalization of input FLN weights. (a) A top-down view of all brain areas on the *r*(*h, L*_*E*_) surface for the 2489th localized bump active state is illustrated in Fig. 6d. (b) The FLN matrix is organized by the HFR and LFR communities. Brain areas in *S*_2_ with a firing rate exceeding 10 Hz are included in the HFR community, while the remaining areas are included in the LFR community. (c) The long-range excitatory input from the HFR community to brain areas within the HFR community is represented by blue dots, and the input from the HFR community to brain areas within the LFR community is depicted by brown dots. (d) Similarly, the long-range excitatory input from the LFR community to brain areas within the LFR community is shown by blue dots. In contrast, the input from the LFR community to brain areas within the HFR community is represented by brown dots.

**Figure S16:**
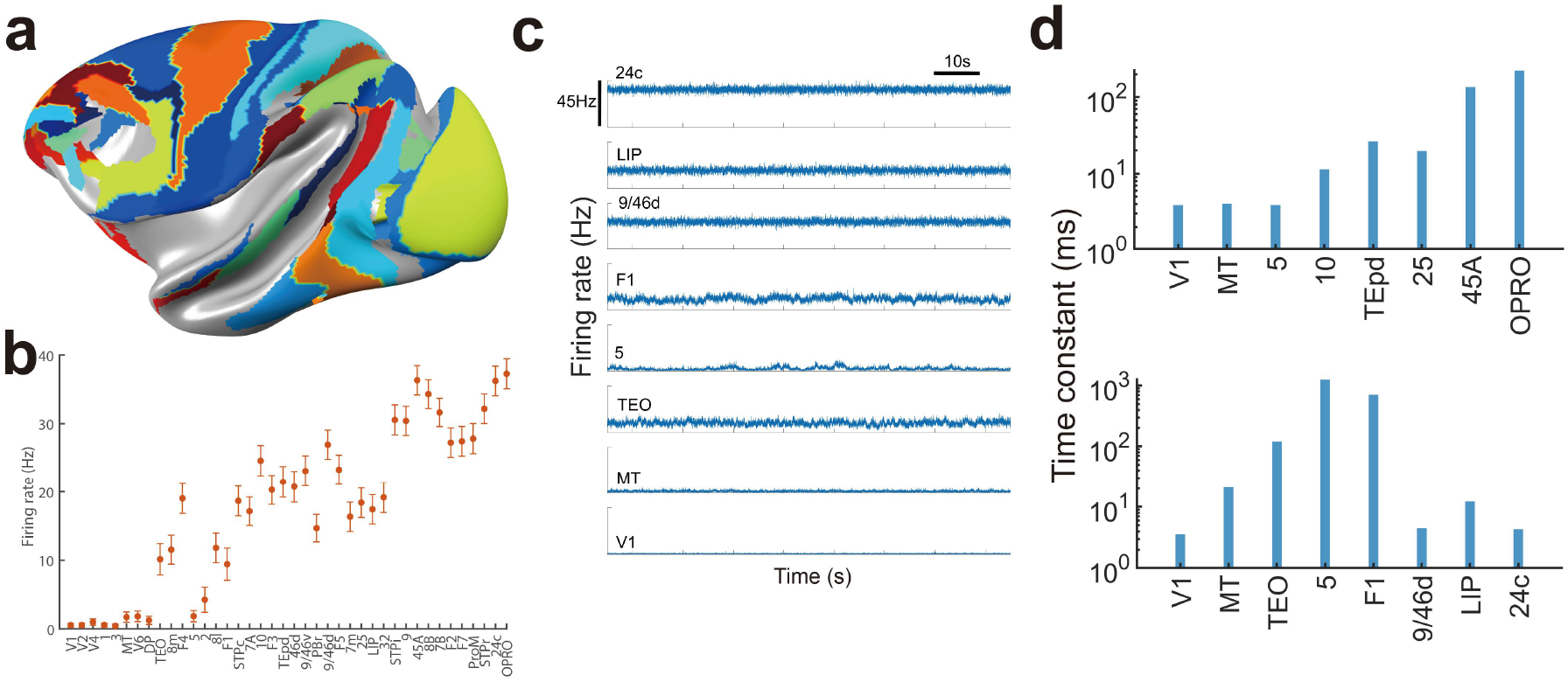
Bifurcation in the space of the connectome-based cortical models of the macaque monkey (a-d) (Mejias and Wang 2022). (a) Lateral view of the macaque neocortex surface with 40 model areas shown in color. (b) The firing rates of 40 brain regions are ranked according to their hierarchical positions, consistent with the classification used in Froudist-Walsh et al. (Froudist-Walsh et al. 2021). (c) Firing rate time series of 8 chosen brain areas when neocortex model is in a delay period working memory state for the model in b. (d) Bar chart of time constants in 8 selected brain areas during the resting state (upper panel) and delay-period working memory state (lower panel).

## References

Abbott, L. F. and F. S. Chance (2005). Drivers and modulators from push-pull and balanced synaptic input. Progress in Brain Research 149, 147–155.

Amit, D. J. (1995). The Hebbian paradigm reintegrated: local reverberations as internal representations. Behav. Brain Sci. 18, 617–626.

Baeg, E. H., Y. B. Kim, K. Huh, I. Mook-Jung, H. T. Kim, and M. W. Jung (2003). Dynamics of population code for working memory in the prefrontal cortex. Neuron 40, 177–188.

Barbosa, J., H. Stein, R. L. Martinez, A. Galan-Gadea, S. Li, J. Dalmau, K. C. Adam, J. Valls-Solé, C. Constantinidis, and A. Compte (2020). Interplay between persistent activity and activity-silent dynamics in the prefrontal cortex underlies serial biases in working memory. Nature Neuroscience 23, 1016–1024.

Blondel, V. D., J.-L. Guillaume, R. Lambiotte, and E. Lefebvre (2008). Fast unfolding of communities in large networks. Journal of Statistical Mechanics: Theory and Experiment 2008, P10008.

Bondy, A. G., J. A. Charlton, T. Z. Luo, C. D. Kopec, W. M. Stagnaro, S. J. C. Venditto, L. Lynch, S. Janarthanan, S. N. Oline, T. D. Harris, et al. (2025). Brain-wide coordination of decision formation and commitment. bioRxiv, 2024–08.

Britten, K. H., W. T. Newsome, M. N. Shadlen, S. Celebrini, and J. A. Movshon (1996). A relationship between behavioral choice and the visual responses of neurons in macaque mt. Visual neuroscience 13 (1), 87–100.

Brody, C. D. and T. D. Hanks (2016). Neural underpinnings of the evidence accumulator. Current Opinion in Neurobiology 37, 149–157.

Cavanagh, S. E., J. P. Towers, J. D. Wallis, L. T. Hunt, and S. W. Kennerley (2018). Reconciling persistent and dynamic hypotheses of working memory coding in prefrontal cortex. Nat Commun 9, 3498.

Chaudhuri, R., K. Knoblauch, M. A. Gariel, H. Kennedy, and X.-J. Wang (2015). A large-scale circuit mechanism for hierarchical dynamical processing in the primate cortex. Neuron 88, 419–431.

Chen, S., Y. Liu, Z. A. Wang, J. Colonell, L. D. Liu, H. Hou, N.-W. Tien, T. Wang, T. Harris, S. Druckmann, et al. (2024). Brain-wide neural activity underlying memory-guided movement. Cell 187 (3), 676–691.

Christophel, T. B., P. C. Klink, B. Spitzer, P. R. Roelfsema, and J. D. Haynes (2017). The distributed nature of working memory. Trends Cogn. Sci. 21, 111–124.

Coifman, R. R. and S. Lafon (2006). Diffusion maps. Applied and computational harmonic analysis 21 (1), 5–30.

Ding, X., S. Froudist-Walsh, J. Jaramillo, J. Jiang, and X.-J. Wang (2024). Cell type-specific connectome predicts distributed working memory activity in the mouse brain. elife 13, e85442.

Douglas, R. J. and K. A. Martin (2007). Mapping the matrix: the ways of neo-cortex. Neuron 56, 226–238.

Douglas, R. J. and K. A. C. Martin (2004). Neuronal circuits of the neocortex. Annu Rev Neurosci 27, 419–451.

Ercsey-Ravasz, M., N. T. Markov, C. Lamy, D. C. Van Essen, K. Knoblauch, Z. Toroczkai, and H. Kennedy (2013). A predictive network model of cerebral cortical connectivity based on a distance rule. Neuron 80, 184–197.

Findling, C., F. Hubert, I. B. Laboratory, L. Acerbi, B. Benson, J. Benson, D. Birman, N. Bonacchi, E. K. Buchanan, S. Bruijns, et al. (2025). Brain-wide representations of prior information in mouse decision-making. Nature 645, 192–200.

Fodor, J. A. (1983). The Modularity of Mind: An Essay on Faculty Psychology. MIT Press: Cambridge, MA.

Fornito, A., A. Zalesky, and E. Bullmore (2016). Fundamentals of Brain Network Analysis. Academic Press: London.

Froudist-Walsh, S., D. P. Bliss, X. Ding, L. Jankovic-Rapan, M. Niu, K. Knoblauch, K. Zilles, H. Kennedy, N. Palomero-Gallagher, and X.-J. Wang (2021). A dopamine gradient controls access to distributed working memory in monkey cortex. Neuron 109, 3500–3520.

Gămănuţ, R., H. Kennedy, Z. Toroczkai, M. Ercsey-Ravasz, D. C. Van Essen, K. Knoblauch, and A. Burkhalter (2018). The mouse cortical connectome, characterized by an ultra-dense cortical graph, maintains specificity by distinct connectivity profiles. Neuron 97, 698–715.

Gold, J. I. and M. N. Shadlen (2007). The neural basis of decision making. Annu. Rev. Neurosci. 30, 535–574.

Goldman-Rakic, P. S. (1995). Cellular basis of working memory. Neuron 14, 477– 485.

Guo, Z. V., H. K. Inagaki, K. Daie, S. Druckmann, C. R. Gerfen, and K. Svoboda (2017). Maintenance of persistent activity in a frontal thalamocortical loop. Nature 545, 181–186.

Harris, J. A., S. Mihalas, K. E. Hirokawa, J. D. Whitesell, H. Choi, A. Bernard, P. Bohn, S. Caldejon, L. Casal, A. Cho, et al. (2019). Hierarchical organization of cortical and thalamic connectivity. Nature 575, 195–202.

Harrison, S. A. and F. Tong (2009). Decoding reveals the contents of visual working memory in early visual areas. Nature 458, 632–635.

International Brain Laboratory, B. Benson, J. Benson, D. Birman, N. Bonacchi, K. Bougrova, S. A. Bruijns, M. Carandini, J. A. Catarino, G. A. Chapuis, et al. (2025). A brain-wide map of neural activity during complex behaviour. Nature 645, 177–191.

Kanwisher, N. (2025). Animal models of the human brain: Successes, limitations, and alternatives. Current Opinion in Neurobiology 90, 102969.

Khilkevich, A., M. Lohse, R. Low, I. Orsolic, T. Bozic, P. Windmill, and T. D. Mrsic-Flogel (2024). Brain-wide dynamics linking sensation to action during decision-making. Nature 634 (8035), 890–900.

Kim, J.-N. and M. N. Shadlen (1999). Neural correlates of a decision in the dorsolateral prefrontal cortex of the macaque. Nature Neurosci. 2, 176–183.

Kim, Y., G. R. Yang, K. Pradhan, K. U. Venkataraju, M. Bota, L. C. Garcia Del Molino, G. Fitzgerald, K. Ram, M. He, J. M. Levine, P. Mitra, Z. J. Huang, X. J. Wang, and P. Osten (2017). Brain-wide maps reveal stereotyped cell-type-based cortical architecture and subcortical sexual dimorphism. Cell 171, 456–469.

Klatzmann, U., S. Froudist-Walsh, D. P. Bliss, P. Theodoni, J. Mejías, M. Niu, L. Rapan, N. Palomero-Gallagher, C. Sergent, S. Dehaene, et al. (2025). A dynamic bifurcation mechanism explains cortex-wide neural correlates of conscious access. Cell Reports 44, 115372.

Kuznetsov, Y. A., I. A. Kuznetsov, and Y. Kuznetsov (1998). Elements of Applied Bifurcation Theory, Volume 112. Springer.

Landau, L. D. and E. M. Lifshitz (2013). Statistical Physics. Elsevier.

Leavitt, M. L., D. Mendoza-Halliday, and J. C. Martinez-Trujillo (2017). Sustained activity encoding working memories: not fully distributed. Trends in Neurosci. 40, 328–346.

Li, S. and X.-J. Wang (2022). Hierarchical timescales in the neocortex: mathematical mechanism and biological insights. Pro. Natl. Acad. Sci. (USA) 119, e2110274119.

Logothetis, N. K., J. Pauls, M. Augath, T. Trinath, and A. Oeltermann (2001). Neurophysiological investigation of the basis of the fMRI signal. Nature 412, 150–157.

Margulies, D. S., S. S. Ghosh, A. Goulas, M. Falkiewicz, J. M. Huntenburg, G. Langs, G. Bezgin, S. B. Eickhoff, F. X. Castellanos, M. Petrides, E. Jefferies, and J. Smallwood (2016). Situating the default-mode network along a principal gradient of macroscale cortical organization. Proc. Natl. Acad. Sci. U.S.A. 113, 12574–12579.

Markov, N. T., M. M. Ercsey-Ravasz, A. R. Ribeiro Gomes, C. Lamy, L. Magrou, J. Vezoli, P. Misery, A. Falchier, R. Quilodran, M. A. Gariel, J. Sallet, R. Gamanut, C. Huissoud, S. Clavagnier, P. Giroud, D. Sappey-Marinier, P. Barone, C. Dehay, Z. Toroczkai, K. Knoblauch, D. C. Van Essen, and H. Kennedy (2014). A weighted and directed interareal connectivity matrix for macaque cerebral cortex. Cereb. Cortex 24, 17–36.

Markov, N. T., J. Vezoli, P. Chameau, A. Falchier, R. Quilodran, C. Huissoud, C. Lamy, P. Misery, P. Giroud, S. Ullman, P. Barone, C. Dehay, K. Knoblauch, and H. Kennedy (“2014”). Anatomy of hierarchy: feedforward and feedback pathways in macaque visual cortex. J. Comp. Neurol. 522, 225–259.

Mazurek, M. E., J. D. Roitman, J. Ditterich, and M. N. Shadlen (2003). A role for neural integrators in perceptual decision making. Cereb Cortex. 13, 1257–1269.

Mejias, J. F. and X.-J. Wang (2022). Mechanisms of distributed working memory in a large-scale model of the macaque neocortex. eLife 11, e72136.

Mendoza-Halliday, D., S. Torres, and J. C. Martinez-Trujillo (2014). Sharp emergence of feature-selective sustained activity along the dorsal visual pathway. Nat. Neurosci. 17, 1255–1262.

Miller, E. K., M. Lundqvist, and A. M. Bastos (2018). Working memory 2.0. Neuron 100, 463–475.

Murray, J. D., A. Bernacchia, D. J. Freedman, R. Romo, J. D. Wallis, X. Cai, C. Padoa-Schioppa, T. Pasternak, H. Seo, D. Lee, and X. J. Wang (2014). A hierarchy of intrinsic timescales across primate cortex. Nat. Neurosci. 17, 1661–1663.

Murray, J. D., A. Bernacchia, N. A. Roy, C. Constantinidis, R. Romo, and X. J. Wang (2017). Stable population coding for working memory coexists with heterogeneous neural dynamics in prefrontal cortex. Proc. Natl. Acad. Sci. U.S.A. 114, 394–399.

Murray, J. D., J. Jaramillo, and X. J. Wang (2017). Working memory and decision-making in a frontoparietal circuit model. J. Neurosci. 37, 12167–12186.

Newsome, W. T., K. H. Britten, and J. A. Movshon (1989). Neuronal correlates of a perceptual decision. Nature 341, 52–54.

Oh, S. W., J. A. Harris, L. Ng, B. Winslow, N. Cain, S. Mihalas, Q. Wang, C. Lau, L. Kuan, A. M. Henry, M. T. Mortrud, B. Ouellette, T. N. Nguyen, S. A. Sorensen, C. R. Slaughterbeck, W. Wakeman, Y. Li, D. Feng, A. Ho, E. Nicholas, K. E. Hirokawa, P. Bohn, K. M. Joines, H. Peng, M. J. Hawrylycz, J. W. Phillips, J. G. Hohmann, P. Wohnoutka, C. R. Gerfen, C. Koch, A. Bernard, C. Dang, A. R. Jones, and H. Zeng (2014). A mesoscale connec-tome of the mouse brain. Nature 508, 207–214.

Panichello, M. F., D. Jonikaitis, Y. J. Oh, S. Zhu, E. B. Trepka, and T. Moore (2024). Intermittent rate coding and cue-specific ensembles support working memory. Nature 636, 422–429.

Perich, M. G. and K. Rajan (2020). Rethinking brain-wide interactions through multi-region ‘network of networks’ models. Current opinion in neurobiology 65, 146–151.

Roitman, J. D. and M. N. Shadlen (2002). Response of neurons in the lateral intraparietal area during a combined visual discrimination reaction time task. J. Neurosci. 22, 9475–9489.

Scheffer, M. (2009). Critical Transitions in Nature and Society. Princeton University Press.

Shew, W. L. and D. Plenz (2013). The functional benefits of criticality in the cortex. The Neuroscientist 19, 88–100.

Shi, Y.-L., R. Zeraati, International Brain Laboratory, A. Levina, and T. Engel (2025). Brain-wide organization of intrinsic timescales at single-neuron resolution. bioRxiv, 2025–08.

Soltani, A., J. D. Murray, H. Seo, and D. Lee (2021). Timescales of cognition in the brain. Current opinion in behavioral sciences 41, 30–37.

Song, H. F., H. Kennedy, and X.-J. Wang (2014). Spatial embedding of similarity structure in the cerebral cortex. Proc. Natl. Acad. Sci. (USA). 111, 16580– 16585.

Song, M., E. J. Shin, H. Seo, A. Soltani, N. A. Steinmetz, D. Lee, M. W. Jung, and S.-B. Paik (2024). Hierarchical gradients of multiple timescales in the mammalian forebrain. Proceedings of the National Academy of Sciences 121, e2415695121.

Sporns, O. (2014). Contributions and challenges for network models in cognitive neuroscience. Nat. Neurosci. 17, 652–660.

Sreenivasan, K. K. and M. D’Esposito (2019). The what, where and how of delay activity. Nat. Rev. Neurosci. 20, 466–481.

Stokes, M. G., M. Kusunoki, N. Sigala, H. Nili, D. Gaffan, and J. Duncan (2013). Dynamic coding for cognitive control in prefrontal cortex. Neuron 78, 364–375.

Strogatz, S. H. (2016). Nonlinear Dynamics and Chaos: With Applications to Physics, Biology, Chemistry and Engineering (Second edition ed.). lOxford, Britain: Taylor & Francis Group.

Suzuki, M. and J. Gottlieb (2013). Distinct neural mechanisms of distractor suppression in the frontal and parietal lobe. Nat. Neurosci. 16, 98–104.

Tredicce, J. R., G. L. Lippi, P. Mandel, B. Charasse, A. Chevalier, and B. Picqué (2004). Critical slowing down at a bifurcation. American Journal of Physics 72, 799–809.

Vezoli, J., L. Magrou, R. Goebel, X.-J. Wang, K. Knoblauch, M. Vinck, and H. Kennedy (2021). Cortical hierarchy, dual counterstream architecture and the importance of top-down generative networks. Neuroimage 225, 117479.

Wang, X.-J. (2001). Synaptic reverberation underlying mnemonic persistent activity. Trends in Neurosci. 24, 455–463.

Wang, X.-J. (2002). Probabilistic decision making by slow reverberation in cortical circuits. Neuron 36, 955–968.

Wang, X.-J. (2020). Macroscopic gradients of synaptic excitation and inhibition in the neocortex. Nature Reviews Neuroscience 21, 169–178.

Wang, X.-J. (2021). 50 years of mnemonic persistent activity: Quo vadis? Trends in Neurosci. 44, 888–902.

Wang, X.-J. (2022). Theory of the multiregional neocortex: large-scale neural dynamics and distributed cognition. Ann. Rev. Neurosci. 45, 533–560.

Wimmer, K., A. Compte, A. Roxin, D. Peixoto, A. Renart, and J. de la Rocha (2015). Sensory integration dynamics in a hierarchical network explains choice probabilities in cortical area MT. Nat Commun 6, 6177.

Wong, K. F. and X.-J. Wang (2006). A recurrent network mechanism of time integration in perceptual decisions. J. Neurosci. 26, 1314–1328.

Zou, L., N. Palomero-Gallagher, D. Zhou, S. Li, and J. F. Mejias (2023). Distributed evidence accumulation across macaque large-scale neocortical networks during perceptual decision making. bioRxiv, 2023–12.

